# A connectomic study of a petascale fragment of human cerebral cortex

**DOI:** 10.1101/2021.05.29.446289

**Authors:** Alexander Shapson-Coe, Michał Januszewski, Daniel R. Berger, Art Pope, Yuelong Wu, Tim Blakely, Richard L. Schalek, Peter H. Li, Shuohong Wang, Jeremy Maitin-Shepard, Neha Karlupia, Sven Dorkenwald, Evelina Sjostedt, Laramie Leavitt, Dongil Lee, Luke Bailey, Angerica Fitzmaurice, Rohin Kar, Benjamin Field, Hank Wu, Julian Wagner-Carena, David Aley, Joanna Lau, Zudi Lin, Donglai Wei, Hanspeter Pfister, Adi Peleg, Viren Jain, Jeff W. Lichtman

## Abstract

We acquired a rapidly preserved human surgical sample from the temporal lobe of the cerebral cortex. We stained a 1 mm^3^ volume with heavy metals, embedded it in resin, cut more than 5000 slices at ∼30 nm and imaged these sections using a high-speed multibeam scanning electron microscope. We used computational methods to render the three-dimensional structure containing 57,216 cells, hundreds of millions of neurites and 133.7 million synaptic connections. The 1.4 petabyte electron microscopy volume, the segmented cells, cell parts, blood vessels, myelin, inhibitory and excitatory synapses, and 104 manually proofread cells are <u>available to peruse online</u>. Many <u>interesting and unusual features</u> were evident in this dataset. Glia outnumbered neurons 2:1 and oligodendrocytes were the most common cell type in the volume. Excitatory spiny neurons comprised 69% of the neuronal population, and excitatory synapses also were in the majority (76%). The synaptic drive onto spiny neurons was biased more strongly toward excitation (70%) than was the case for inhibitory interneurons (48%). Despite incompleteness of the automated segmentation caused by split and merge errors, we could automatically generate (and then validate) connections between most of the excitatory and inhibitory neuron types both within and between layers. In studying these neurons we found that deep layer excitatory cell types can be classified into new subsets, based on structural and connectivity differences, and that chandelier interneurons not only innervate excitatory neuron initial segments as previously described, but also each other’s initial segments. Furthermore, among the thousands of weak connections established on each neuron, there exist rarer highly powerful axonal inputs that establish multi-synaptic contacts (up to ∼20 synapses) with target neurons. Our analysis indicates that these strong inputs are specific, and allow small numbers of axons to have an outsized role in the activity of some of their postsynaptic partners.

## Introduction

While the functions carried out by most of the vital organs in humans are not remarkably different when compared to other animals, the human brain clearly separates us from the rest of life on the planet. It is a vastly complicated tissue, and to date, little is known about its cellular microstructure, and in particular its synaptic circuits. These circuits underlie the unparalleled capabilities of the human mind, and when disrupted, likely underlie incurable disorders of human brain function. One critical barrier that has prevented detailed knowledge of the cells and circuits of the human brain has been the access to high quality human brain tissue. Organ biopsies provide valuable information in many human organ systems, but biopsies are rarely done in the brain except to examine or excise neoplastic masses, and hence, most of them are of little value for the investigation of human brain structure. Moreover, for many organ systems, tissue from model organisms are useful for human disease: a mouse liver or lung can provide insights into human pathophysiology. The human brain is different. We don’t have many good models of cognitive and developmental disorders (although this may be changing with genetic manipulation of non-human primates ^1^). Also, the human brain is clearly not the same as a mouse’s brain, and if we go by functional repertoire, humans are dramatically different from other primates. We therefore must find ways to explore human brain tissue per se. One attempt has been to use brain organoids made from human cells. This is certainly a promising field ^2–4^ but at present they do not approximate brain tissue architectonics (e.g., cortical layers are not present) nor do they have circuits that resemble those in the human brain. An alternative to organoids is the direct approach: use state-of-the-art tools to map cells and circuits from human specimens. Human specimens are available, owing to neurosurgical interventions for neurological conditions where the cortex is discarded or destroyed, because it obstructs access. Human tissue that is a byproduct of neurosurgical procedures on patients can be leveraged to understand normal, and perhaps disordered human neural circuits. One source of such samples are individuals with drug-resistant epilepsy. These patients are sometimes treated by invasive surgical extirpation of the focus site determined by EEG ^5^. The most common type of focal epilepsy occurs in the medial part of the temporal lobe, and when these are treated via neurosurgery, it is by unilateral resection of medial structures often including the hippocampus. Medial structure access is usually by transection through the overlying temporal cortex, and significant volumes of temporal lobe (approaching one half a cubic cm) are sometimes removed. The number of patients with drug-resistant epilepsy is large (in the US, about 750,000 patients) but very few (0.2%) undergo epilepsy surgery per year (Natl. Assoc. of Epilepsy Centers, 2014). Nonetheless, 1,500 patients per year in the US potentially could provide tissue.

Here we describe such a sample, a cubic millimeter in volume, that extends through all cortical layers which we imaged at the ultrastructural level with serial high-throughput electron microscopy, and analyzed with computational approaches. The acquisition of digital human brain tissue at this large scale and this fine resolution, enables not only access to neuronal circuitry comprising thousands of neurons and millions of synapses, but such volume microscopy also provides a clear view of all the other tissue elements that comprise human brain matter including all glial cells, the blood vasculature, and the relations between these various cell types. A wide range of questions related to human brain biology are thus open to scrutiny from a single sample. Because the dataset is large and incompletely scrutinized, to aid in its analysis we are <u>sharing all of the data online</u>.

## Results

The sample we analyzed is from an anonymized 45 year old patient with drug-resistant epilepsy who had a left hippocampal resection via the anterior temporal lobe at Massachusetts General Hospital, Boston, MA. Traditional neuropathology showed that the ablated cortex was not abnormal, but the underlying hippocampus was sclerotic with neuron loss, as is typical in such epilepsy patients.

We ran the resected temporal lobe fragment (superficial to the hippocampus) through a pipeline of steps. We used rapid immersion fixation for excellent preservation of ultrastructure. The tissue was further preserved and stained with osmium and other heavy metals using a ROTO staining protocol ^6^. The sample was then embedded in a resin (epon) block and trimmed. Using an automatic tape-collecting ultramicrotome (ATUM; ^7^) we sectioned 5,292 sections at a section thickness that averaged 33 nm (range 30-40 nm) and imaged with a multibeam scanning electron microscope (**Fig. 1**). The images were acquired with pixels that were 4 x 4 nm^2^. The raw data size was up to 350 GB per section due to necessary overlap at the edges of the tiles, or for the entire set of sections ∼2.1 petabytes (PB). While the scope was acquiring images, a custom-built workflow manager assessed each tile for quality and flagged problems. The total throughput of the image acquisition ranged from 125 million (M) pixels per second to 190 M pixels per second. The majority of the data was acquired at 190 M pixels per second. The total imaging time for the 1 mm^3^ sample was 326 days.

**Figure 1.**
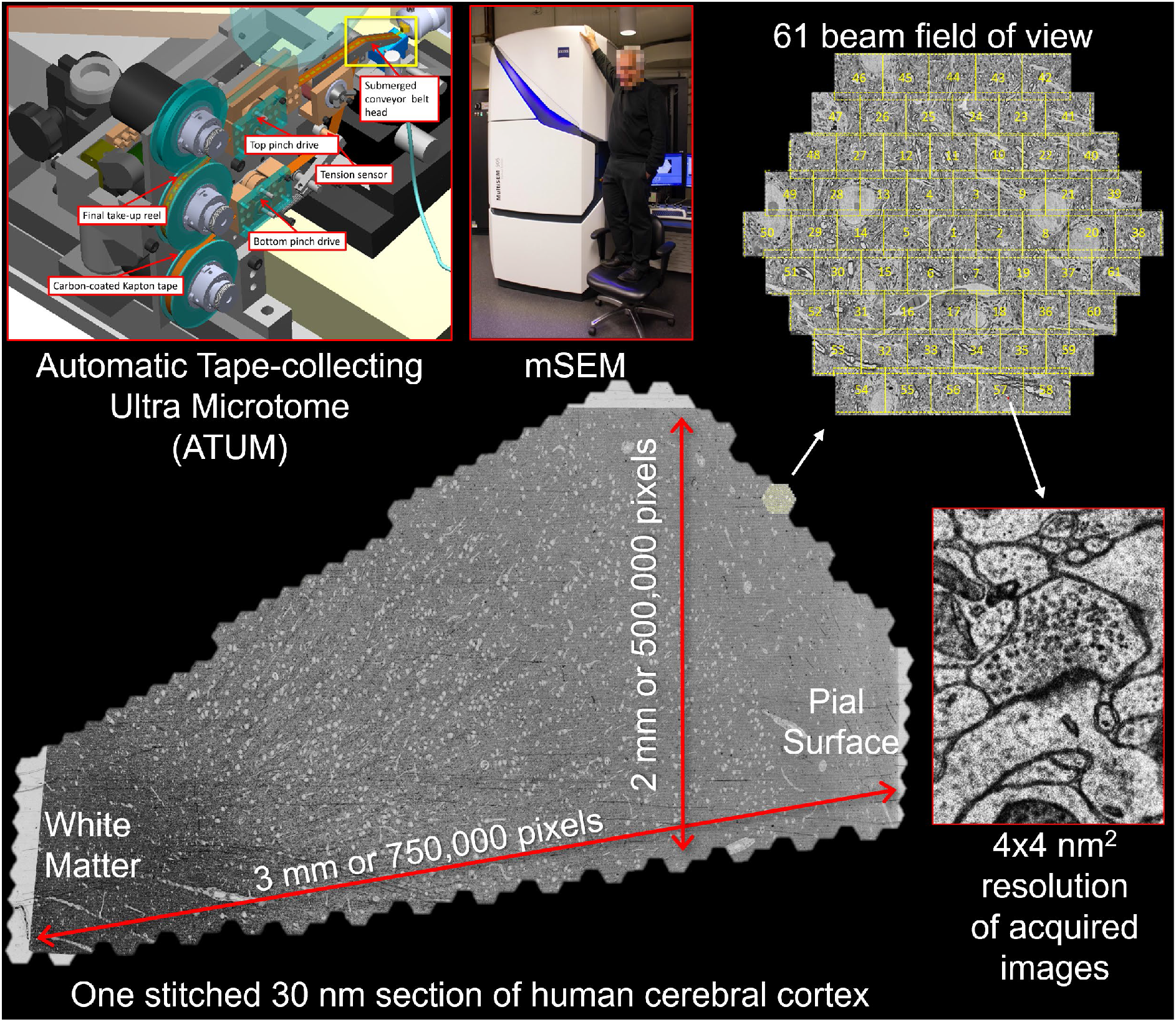
Image acquisition for the human brain sample. A fresh surgical cerebral cortex sample was rapidly preserved, then stained, embedded in resin, and sectioned. More than 5000 sequential ∼30 nm sections were collected on tape using an ATUM (upper left panel). Yellow box shows the site where the brain sample is cut with the diamond knife and thin sections are collected onto the tape. The tape was then cut into strips and imaged in a multibeam scanning electron microscope (mSEM). This large machine (see middle panel with person on chair as reference) uses 61 beams that image a hexagonal area of about ∼10,000 μm^2^ simultaneously (see upper right). For each thin section, all the resulting tiles are then stitched together. One such stitched section is shown (bottom). This section is about 4 mm^2^ in area and was imaged with 4 x 4 nm^2^ pixels (see image of synapse at lower right). Given the necessity of some overlap between the stitched tiles, this single section required the collection of more than 300 GB of data.

The acquired image data was then re-composed into three-dimensional cellular objects from which all the neuronal and glial elements could be itemized. The neurons in the volume and the processes of neurons passing through the volume were then rendered and their synapses identified to give rise to the connectomic reconstruction. This entire compute-intensive process consisted of a series of largely automated workflows (**Fig. 2,** and see Methods). The raw acquired image tiles were first stitched together and coarsely aligned by using microscope stage coordinates, semi-automated feature correspondences, and image patch cross-correlations to relax an elastic triangular mesh of each tile and each section (**Fig. 2A**, left) ^8^. A fine-scale refining alignment based on optical flow between neighboring sections removed remaining drift and jitter from the volume (**Fig. 2A**, center and right). From a total of 247 M tiles (2.1 PB), 196 M tiles containing cortex (1.7 PB) were stitched, aligned, and ultimately segmented, creating a unified ∼1.4 PB human cerebral cortex image volume.

**Figure 2:**
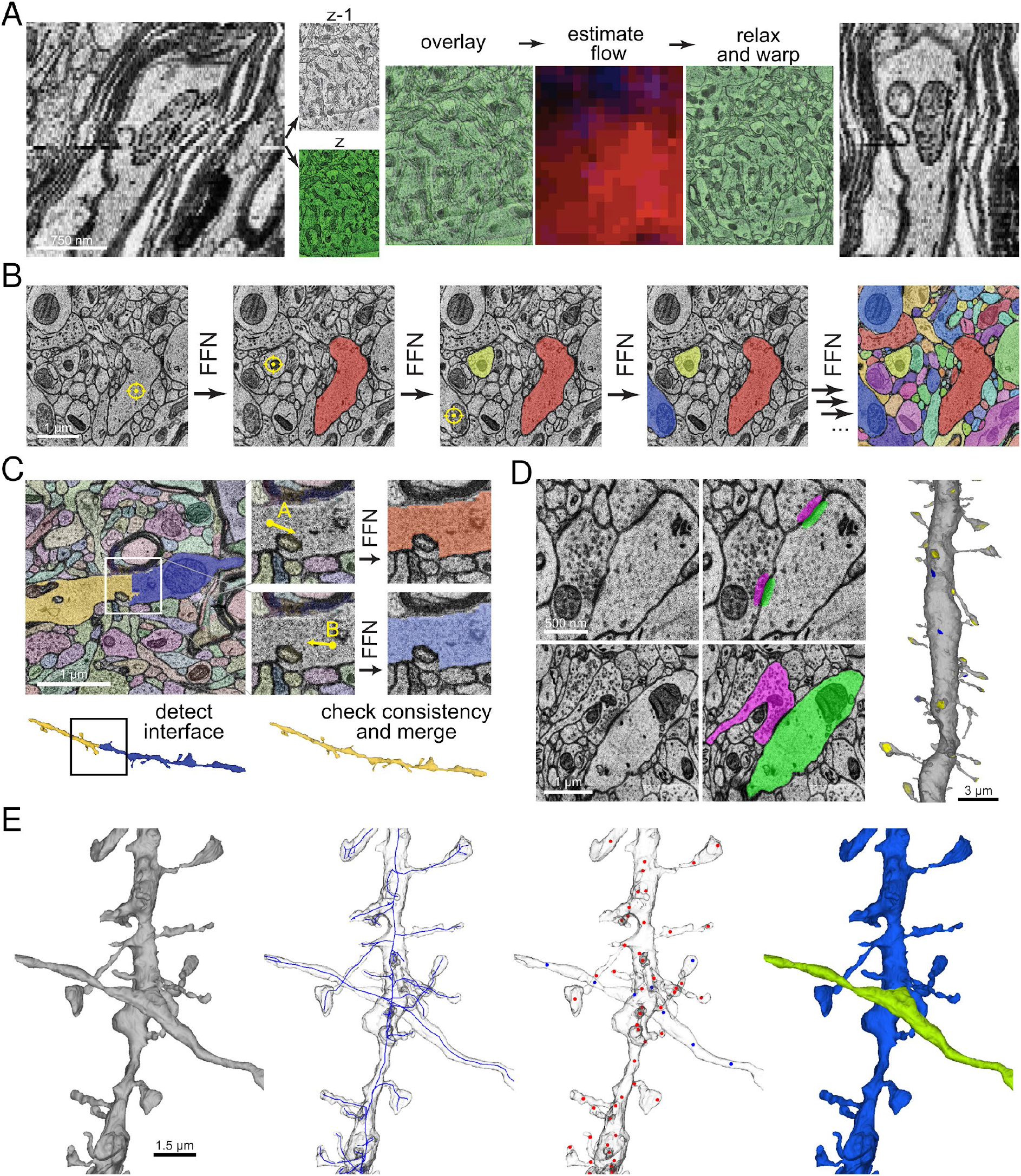
Reconstruction pipeline overview. **A:** Fine-scale alignment with optical flow. Left: An XZ cross-section of the initial coarsely aligned subvolume exhibits drift and jitter. Center:Two adjacent XY sections z (green) and z-1 are overlaid to illustrate their misalignment. Image patch-based cross-correlation computes an XY flow field between them. Red and blue intensities, which indicate the respective horizontal and vertical flow components are used to warp one of the sections, improving their alignment (relax and warp overlay). Right: XZ view of the same subvolume with flow realignment applied. **B:** Example of sequential segmentation with an FFN. XY cross-sections illustrate the 3D segmentation process. Each yellow crosshair indicates the seed location for the next segment. **C:** FFN agglomeration. Left: Site between two adjacent base segments (white box in 2D, black box in 3D below) is a candidate agglomeration location. Center: FFN segmentation is seeded from points A and B independently. Right: If the resulting A and B segmentations are mutually consistent, the object pair is merged (below). **D:** Synapse detection and classification. Top: XY cross-section of EM image input to synapse detection model (left), and the resulting presynaptic (magenta) and postsynaptic (green) prediction masks (right). Bottom: cross-section of EM image (left) and presynaptic (magenta) and postsynaptic (green) object segmentation (right) inputs to excitatory versus inhibitory classification model. Right: <u>3D render</u> of a dendrite with predicted incoming excitatory (yellow) and inhibitory (blue) synaptic sites. **E:** Subcompartment prediction and merge error correction (<u>link</u>). Leftmost: a single reconstructed object with a merge error where axon and dendrite cross near each other. Left center: the object is converted to a reduced skeleton representation (blue). Right center: fields of view around a subset of skeleton nodes are input to a subcompartment classification model. Red nodes: predicted dendrite; blue nodes: predicted axon. The inconsistency in subcompartment predictions is detected, and the agglomeration graph is cut at the location that maximally improves subcompartment consistency. Rightmost: the separated axon and dendrite after applying the suggested cut.

Reconstruction of the structure of every cell and process in the aligned volume proceeded via Flood-Filling Network (FFN) segmentation and agglomeration (**Fig. 2B and C**) ^9^. Multi-resolution FFN segmentation and oversegmentation consensus produced base segments (also known as supervoxels, **Fig. 2B**) that were then agglomerated via FFN resegmentation followed by mutual consistency criteria to produce more complete reconstructed cells (**Fig. 2C**). Synaptic connections were added by a pre- and postsynaptic masking model applied to EM image blocks (**Fig. 2D,** top), while the polarity of synapses (excitatory versus inhibitory) was predicted by a classification model that considered the EM imagery centered around each putative synapse, as well as the local pre- and postsynaptic neuron segment masks and whether or not the postsynaptic site was a dendritic spine (**Fig. 2D**, bottom and right). A skeleton (“ball and stick”) representation was also automatically generated from the volumetric segmentation, and higher-dimensional embeddings were computed for skeleton nodes with the help of a self-supervised neural network model. Skeletons were then used for automated subcompartment classification, and embeddings for glial type identification.

The agglomerated segmentation was further refined using subcompartment predictions (**Fig. 2E**). A classification model predicted axon, dendrite, astrocyte, soma, cilium, and axon initial segment classes at skeleton node locations distributed throughout each cell^10^. Occasional agglomeration errors produced merges between nearby objects, such as a passing axon and dendrite (**Fig. 2E**, leftmost). We used subcompartment predictions to distinguish the two merged objects (**Fig. 2E**, center right) and allow an automated cut to separate them (**Fig. 2E**, rightmost). The final data set is provided with two different agglomerations: c2 agglomeration favors fewer breaks (and hence longer processes) but with a higher number of incorrect mergers and c3 agglomeration that is more coservative and has shorter fragments but fewer merge errors (both are available <u>here</u>).

For analysis, the various forms of data have been made available in SQL databases that enable specific queries about the neuronal and synaptic circuit data. We wrote software to aid in these queries including modifications to <u>Neuroglancer</u> and a new program, <u>CREST</u>.

### Cellular and Macroscopic Organization

We first analyzed the cellular and macroscopic organization of the brain sample (**Fig. 3**).

**Figure 3:**
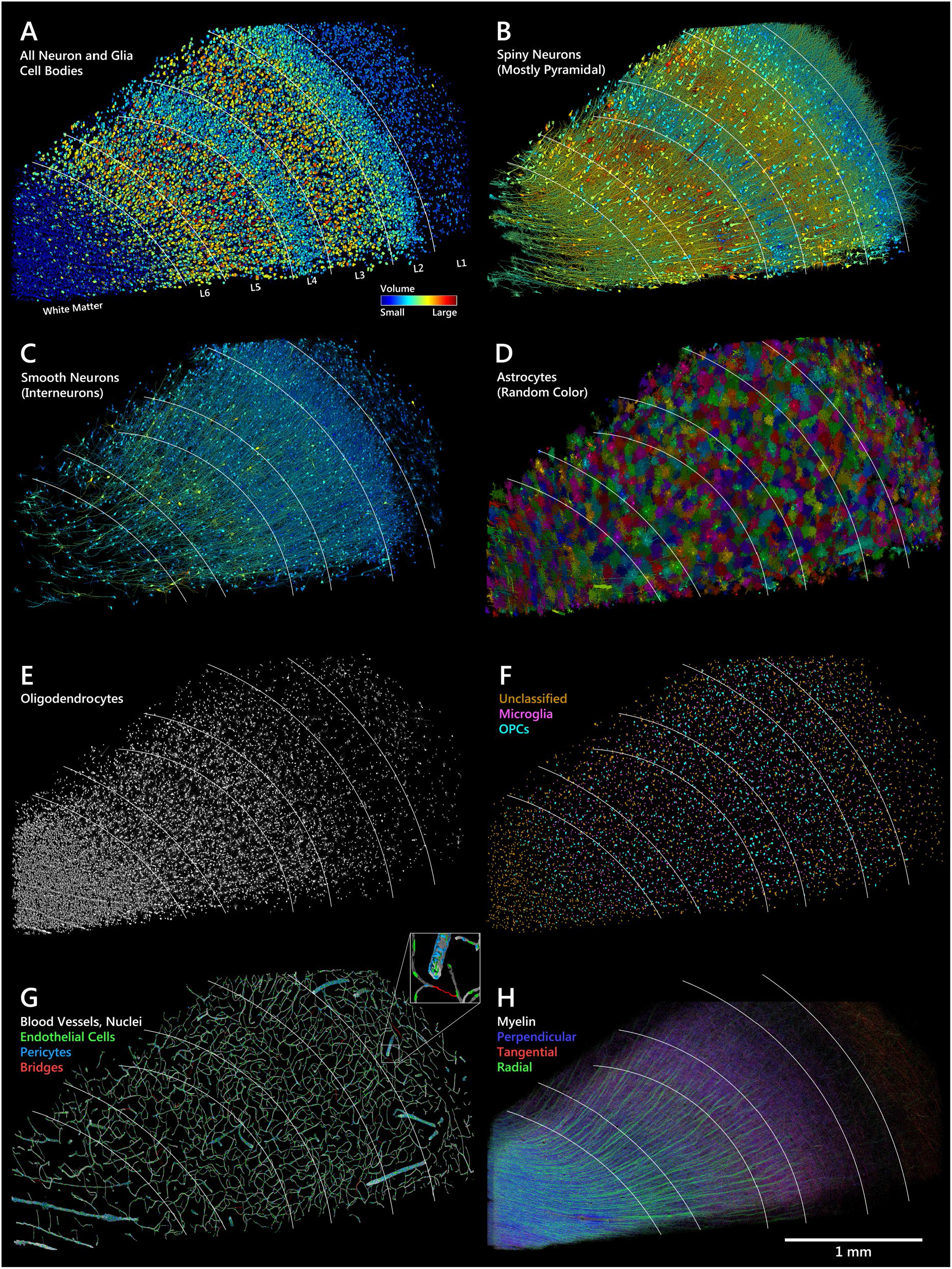
Distribution of cells, blood vessels and myelin in the sample. White lines indicate layer boundaries based on cell clustering. **A**: All 49,080 cell bodies of neurons and glia in the sample, colored by soma volume. **B**: Spiny neurons (putatively excitatory), colored by soma volume. **C**: Interneurons (few spines, putatively inhibitory), colored by soma volume. **D**: Astrocytes mostly tile but in some cases, arbors of nearby astrocytes interdigitate. **E**: Most of the oligodendrocytes in the volume. Note clustering along large blood vessels, especially in white matter. **F**: Cell bodies of microglia and oligodendrocyte precursor cells (OPCs). **G**: Blood vessels and the nuclei of the 8,136 associated cells (<u>link</u>). Inset shows a magnified view of the location of the individual cell types. **H**: Myelinated axons in the volume, color-coded by topological orientation. Most axons in white matter run in the perpendicular direction. Thick axon bundles run between white matter and cortex in the radial direction. In layer 1, a set of large-caliber myelinated axons runs tangentially through our slice, parallel to the pia. In layers 3-6 many myelinated axons also run in diagonal (tangential/perpendicular) directions.

We manually identified all of the cell bodies in the data set (see **Supplementary Table 3**). The cell census included 49,080 neurons and glia (**Fig. 3A**), and 8.1k blood vessel cells (**Fig. 3G**). The number of neurons in this human sample is ∼16k/1 mm^3^, many fold lower than the density of neurons in mouse cortex ^11, 12^. Simply by measuring the neuron and glial cell body cross-sectional areas colorized by size, the cortical layering was obvious (**Fig. 3A**). Two regions that had a paucity of neurons (white matter at the left edge of the images in **Fig. 3**) and the most superficial layer (at the right edge of **Fig. 3A**) were populated mainly by glial cells whose cell body sizes were smaller than neurons (**Fig. 3A**, blue). The largest cells (red) were mostly in a broad deep infragranular band (i.e., closer to the white matter) and a more superficial (supragranular) one. Because the cells were segmented into 3-dimensional objects, their appearance plus the ultrastructural features of somata allowed classification of the cells into types across layers. The largest cell somata belonged to spiny pyramidal neurons. These cells typically possess one large apical dendrite projecting to the most superficial layer (**Fig. 3B**). However, even their somata were distinctly different in size across cortical layers. Non-pyramidal neurons were much less spiny, had smaller cell body sizes and had a less obvious layer arrangement (**Fig. 3C**). We sought to find an objective layering criterion and used cell soma size and clustering density to group cells into three non-adjacent layers and assigned the cells adjacent to these layers to four additional layers (see Methods). This approach generated a 6-layered cortex and white matter. The fiducial lines in each panel of **Fig. 3** are based on these layers (named in **Fig. 3A**).

The glial cells also showed differences between layers (**Fig. 3D, E and F**). The compact and complicated arbors of protoplasmic astrocytes are densely tiled in all layers (**Fig. 3D**), but the fibrous astrocytes in the white matter are more elongated than the protoplasmic astrocytes that occupy the cortical layers. Astrocytes in layer 1 were somewhat smaller in expanse, in line with astrocytes characterized in mouse cortex ^13^. Layer 1 astrocytes also stand out because of higher densities of these cells (see **Supplementary Fig. 1A and 5A**). Interestingly, the astrocyte density divides layer 1 into a superficial and deep region with the upper part of layer 1 being most densely populated with astrocytes, while the deeper part of layer 1 and layer 2 contain fewer astrocytes compared to upper layer 1 or layers 3-6, and even the white matter. Layer 1 is also unusual because it includes small aggregates of astrocytes (**Supplementary Fig. 1B**) where the arbors of neighboring astrocytes extensively intermingle. Transcriptomic characterization of astrocytes in the cerebral cortex has previously separated layer 1 and 2 from the other layers ^14^ or separated astrocytes into three cortical layers ^15^. Layer 1 includes the primate-specific interlaminar astrocytes ^16^, however, the aggregates of highly overlapping astrocytes we see in layer 1 are not interlaminar astrocytes, as they are lacking projections and are further away from the pial surface. We also noted extensive overlap between protoplasmic astrocytes in many layers (**Supplementary Fig. 1C**). This intermingling was unexpected since protoplasmic astrocytes (in rodents) are mainly described to have non-overlapping territories ^17–19^. Oligodendrocytes had a different distribution. They were most plentiful in the white matter, as expected, given their role in myelin formation. There were also long lines of oligodendrocytes that surrounded the larger radially directed blood vessels in the white matter. The ultrastructure of this association is shown in **Supplementary Fig. 2A-C**. We noted perivascular oligodendrocytes ^20^ throughout the volume, with similar distance and interaction to blood vessels as microglia and OPCs (further described in **Supplementary Fig. 2D and E**). Oligodendrocyte density decreased in more superficial cortical layers, reaching a minimum in layer 2 (**Fig. 3E and Supplementary Figure 5A**). In layer 1 they were slightly higher in density, probably related to the horizontally running myelinated axons in this layer (see below). Oligodendrocyte precursor cells which have different morphology and function than oligodendrocytes were difficult to differentiate by appearance from microglia (see **Supplementary Fig. 3**). Therefore, we used the skeleton node embeddings and a small set of manually labeled examples to train a model to classify candidate cells as microglia or OPCs (see Embeddings in Methods). The model predicted 2,049 cells to be microglia and 1,395 cells to be OPCs, whereas 2,836 cells were deemed ambiguous, often because of merge or split errors in the segmentation (see Methods and **Supplementary Fig. 25**). **Fig. 3F** shows the spatial distribution of the three classes. Unclassified cells tend to be at the sides of the sample owing to poorer alignment there and in the white matter, where the appearance of these glial cells is somewhat different than in the cortex. The reconstructed blood vessels (22.6 cms in total length) also did not show much evidence of layer-specific behavior (**Fig. 3G**). There was clearly a lower density in the white matter, presumably because of the lower energy requirements of myelinated axons, as compared to unmyelinated processes ^21^. Panel **Fig. 3G** also shows the location of 4,604 endothelial cells (∼20 per mm of vasculature) lining the vessel lumen (green) and a more heterogeneous group of 3,549 pericytes (blue, ∼15 per mm of vasculature) within the basement membrane but displaced slightly further from the lumen. Identification of smooth muscle cells as a type distinct from pericytes was possible in arteries and arterioles, however this distinction was not as clear in veins and venules as predicted by the expression profiles of these cells ^22^. We estimated the number of vascular smooth muscle cells in the sample to be 574 and that there were 78 fibroblast-like pericytes ^22^ surrounding the smooth muscle cells. In the capillaries, where smooth muscle cells are missing, pericytes (total 2339 in the volume) help regulate blood flow ^23, 24^. Perivascular macrophages ^25^ (total 396) were identified based on location and large intracellular granules, as well as 128 perivascular lymphocytes, also located within the basement membrane. The total number of pericytes included a group of 25 cells we could not categorize further (see Methods for more details about cell type criteria). In addition, 46 circulating white cells (mainly neutrophils) were observed in the blood vasculature of the 1 mm^3^ volume which was not perfused to remove blood.

Although astrocytic end feet touched blood vessels in many locations, their cell bodies were usually not abutting the blood vessels, but microglia, OPCs and oligodendrocytes were adjacent to the blood vessels. We also noted the capillary-free “Pfeifer spaces” surrounding the larger blood vessels ^26^. Based on the number and positioning/orientation of contractile cells, we could separate the larger vessels into arterioles and venules. However, the narrowness of the sample prevented us from reconstructing a full blood circuit from artery to vein. In the vasculature we found 73 thin bloodless bridges connecting different capillaries ^27 28 29^ that were composed of a basement membrane and pericytes but lacking endothelial cells (marked red in **Fig. 3G**, and shown in more detail in **Supplementary Fig. 4**).

From all of this cellular data, we could describe the complete cellular census of this brain sample (**Supplementary Fig. 5** and **Supplementary Table 3**). The oligodendrocyte was the most common cell type. 20,139 oligodendrocytes were found in this sample. Glia outnumber neurons by 2:1 (32,315 versus 16,087) though this varies by cortical layer (see **Supplementary Fig. 5A, B**). The 8,096 vasculature cells were already described above. The most common neurons were spiny cells (10,531 spiny neurons, of which 8,803 had a clear pyramidal shape. The spiny cells accounted for 69% of the neurons in the sample). We describe one new subclass below. The 31% (n=4,688) non-spiny neurons were classified by us as interneurons. There was a subset of neurons that did not easily fit into this binary categorization because either their somata were not fully in the volume, or more rarely, they appeared anomalous in other ways (868 cells). Other unusual neurons are shown in **Supplementary Fig. 6.**

Myelin was found throughout the volume (**Fig. 3H**). Its density was highest in the white matter as expected. There were layer-specific differences in myelin density. The layer with the least myelin was layer 2. As already described, layer 2 was also the layer with the lowest density of oligodendrocytes (see **Fig. 3E** and **Supplementary Fig. 5A**). Because we annotated the myelin and skeletonized the axons it wrapped, it was possible to render not only the density of the myelin, but also its direction. Myelin direction is the basis of the diffusion tensor measurements made in the human brain *in vivo* ^30^. The myelin in the white matter is running primarily orthogonal to the plane of the section and hence appears blue in this direction-color plot. Bundles of myelinated axons project radially (green color) between the white matter and the cortical layers. In layer 3 the myelin is also running mostly orthogonal to the plane of sectioning (blue). In layer 1 myelin is running horizontally within this layer and hence orthogonal to the white matter and the radial myelin and colorized red (see also **Supplementary Fig. 23B**).

Because all the objects in the cortical tissue were annotated by types we could assess the volumetric contribution of different types of cells and cell parts to the cortical parenchyma. By volume, this brain sample was 40.6% unmyelinated axons, 26.1% dendrites, 16.0% astrocytes and other glial cells, 9.6% somata, 7.6% myelinated axons, 0.07% axon initial segments, and 0.03% cilia (**Supplementary Fig. 22**). The ratio of dendritic processes to axonal processes (4.9:1) is tipped more in favor of unmyelinated axons than the volumetric measurements above because individual axons on average occupy less volume than dendrites. In addition to these well known categories, we found in the volume a number of UCOs (unidentified cortical objects) that accounted for very little volume. These are shown in **Supplementary Fig. 7.**

### Synaptic Connectivity

The most functionally significant aspect of cortical tissue is the synaptic connectivity that allows neurons to send and receive signals to and from other neurons. This wiring diagram is likely central to the way human brains store memory and give rise to behavior. The segmentation of neurons described above was insufficient to generate a wiring diagram because these segmentations did not identify synapses. As described in **Fig. 2**, we therefore used machine learning tools to train automated synapse classifiers to identify the pre- and postsynaptic component of each synapse and whether the presynaptic terminal was putatively excitatory or inhibitory. Proofreading showed that the number of missed synapses (i.e., false negatives) was 12%. Because we found no convincing cases of dendrites, glia, or cell somata establishing synapses (i.e., presynaptically) we could eliminate many of the false positives. After these corrections, the final false positive rate was 1.5%. Automated classification of each synapse as excitatory or inhibitory was based on the appearance of each individual synapse and whether its postsynaptic structure was a dendritic spine or not, as well as the whether the presynaptic neuron was an excitatory or inhibitory type, for those neurons where the soma was in the volume. This approach classified excitatory and inhibitory synapses correctly 99.42% and 92.00% of the time, respectively. In total, we found 149.8 M synapses in the volume between axons and either: dendrites (99.4%), AIS (0.197%), or somata (0.393%). There were: 40.6 M (27.13%) inhibitory and 109.2 M excitatory (72.87%). The density of the automatically annotated excitatory synapses was highest in layers 1 and 3 (**Fig. 4A**). The distribution of the inhibitory synapses was different and peaked in layer 1 (**Fig. 4B**). The percentage of excitatory synapses of total E/(E+I) is highest in layer 3 and lowest in layer 1 (**Fig. 4C**). A relatively higher density of inhibitory synapses in human cortex layer 1 has been reported before ^31^. Synapse density estimates for different layers are shown in **Supplementary Fig. 26**.

**Figure 4:**
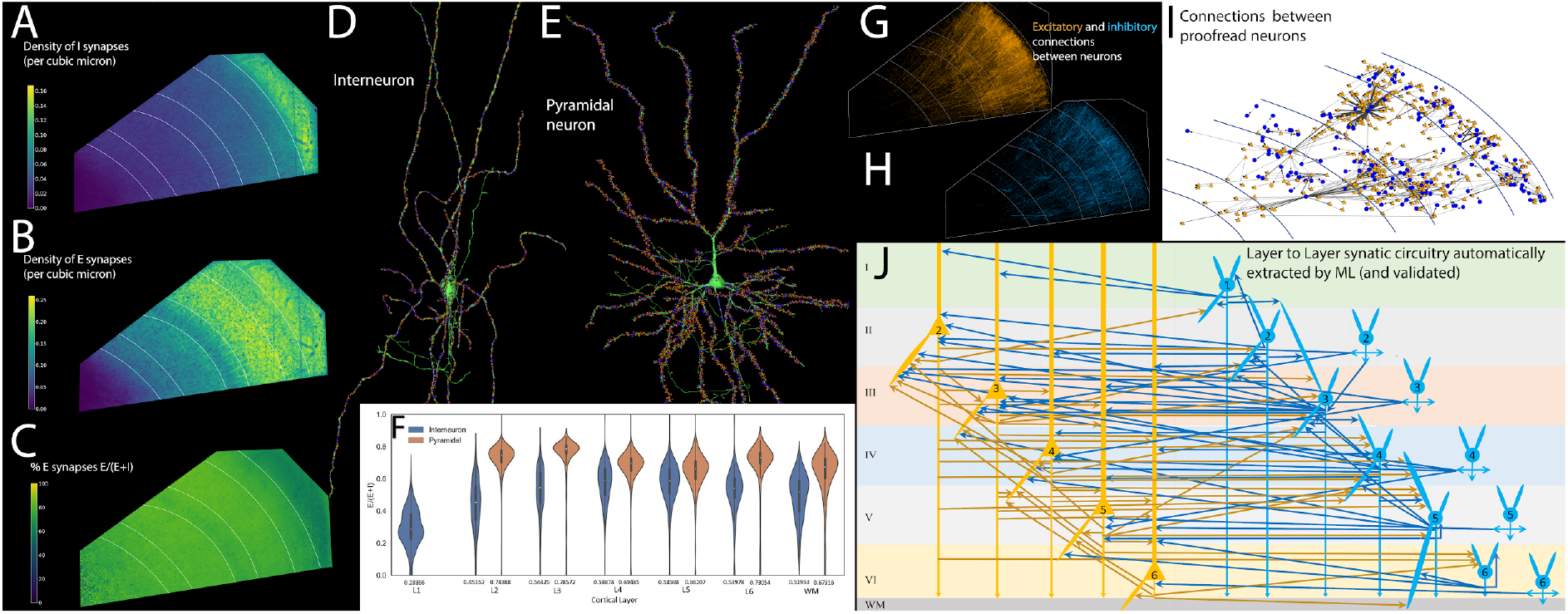
Synapses and circuit reconstruction. **A**: Volumetric density of inhibitory synapses. **B**: Volumetric distribution of excitatory synapses. **C**: Excitatory-inhibitory balance (E/E+I). Lowest values are purple, highest values are yellow. **D**: Interneuron and **E**: Pyramidal neuron with synapse locations shown in orange for synapses classified as excitatory and blue for synapses classified as inhibitory. Note large numbers of I synapses on the pyramidal neuron’s axon initial segment. **F**: Balance of inhibitory and excitatory synapses (E/E+I) onto interneurons (blue) and pyramidal neurons (orange) for neurons with cell bodies in different layers. **G,H**: Excitatory (orange) and inhibitory (blue) synaptic connections between neurons with cell bodies in the volume, based on non-proofread, automatically extracted cells and synapses; high-resolution online versions are available for both <u>inhibitory</u> and <u>excitatory</u> synaptic connections. **I**: Synaptic network between a set of 104 manually proofread neurons and their postsynaptic targets in the volume. Pyramidal cells are shown as orange triangles, interneurons as blue circles. Blue circles indicate spiny stellate neurons. **J**: From the automatically segmented connections we identified 96 categories of synapses (all subsequently validated by proofreading) and diagrammed the layer location of the synapses (arrowheads) and the layer location of the cell somata of the pre- and postsynaptic partners of each category (blue neurons are non-spiny and orange are spiny). Chandelier neurons whose axons innervate initial segments are the rightmost cells.

Because each identified synapse was associated with a postsynaptic target, it was possible to analyze the inhibitory and excitatory drive to every neuron in the volume (see an example interneuron and pyramidal neuron in **Fig. 4D and E**). Inhibitory neurons on average received roughly equal numbers of excitatory and inhibitory synapses (**Fig. 4F**) and they were distributed uniformly along dendrites and the cell somata. Excitatory neurons had relatively few synapses on the cell body and proximal (non-spiny) dendritic branches. For excitatory neurons, the synapses on the cell body and proximal dendrites were largely inhibitory. In contrast, in the spiny dendritic regions there were more excitatory synapses than inhibitory and because so much of the dendritic tree was spiny, the total excitatory drive as percent of total synapses on pyramidal cells was shifted to more excitation than the input to inhibitory neurons. There was also a large number of inhibitory synapses from chandelier interneurons on the axonal initial segment to the vast majority of spiny neurons (e.g., **Fig. 4E**) but not on interneurons. One exception, these chandelier axons also innervated the axon initial segment of other chandelier interneurons (see **Supplementary Fig. 8**). In layers 2-6 and the white matter, the tendency was the same: Pyramidal cells received a greater proportion of excitatory synapses than non-spiny interneurons (**Fig. 4F**).

In this volume there were many hundreds of millions of axons originating from sites that were outside the volume. More rarely were the presynaptic terminals from axons of neurons in the volume. This asymmetry is accounted for by the fact that the majority of axonal input to cells originate from neurons outside the volume and most axons of neurons in the volume project outside the volume where they establish the majority of their synapses. Nonetheless, among the sparse axonal synapses of neurons with somata in the volume, 11,470 (71.3%) were synaptically connected to one or more other neurons with somata in the volume. We have graphed all of these connections (excitatory input **Fig. 4G**; inhibitory input **Fig. 4H**). These synapses gave rise to 29,498 in-volume neuron to neuron connections, and comprised 38,191 synapses. These synapses were mostly on dendrites (36,158, 94.68%), axonal initial segments (1065, 2.79%) and axonal input to neuronal somata (968, 2.53%). This neuron connectivity circuit data is available at full resolution for both <u>inhibitory</u> and <u>excitatory</u> connections.

To better understand potential modes of information flow in the network, we algorithmically traversed the connectome according to its connectivity. Specifically, we chose L4 pyramidal neurons as a starting point because this layer is often the target of feed-forward connections from other brain regions^32^ and input to other cortical layers^33^ and then traversed their postsynaptic partners if a threshold criteria was satisfied (see Analysis of information flow through network in Methods). We found that the layer 4 pyramidal cells (n = 908) primarily connect to cells in layer 3, which in turn connect to cells in layer 2. As this progression iterates toward the upper surface of the cortex, connections to cells in deeper layers 6 and 5 also become visible (see <u>video online</u>). Because the number of connections being traversed is extremely sparse, owing to axonal breaks in the automatic segmentation, and the fact that only connections between neurons with somata in the volume are being considered, this information flow should not be confused with a neural activity simulation of this cortical slice.

Our large-scale automated segmentation combined with the development of efficient segmentation proofreading tools such as <u>CREST</u> also make possible the reconstruction of verified neural networks composed of the outputs of hundreds of neurons to hundreds of postsynaptic neurons. To obtain such a network, we proofread 104 neurons, which together with their postsynaptic target neurons form a network of 585 neurons, shown in **Fig. 4I**. In this network axonal branches that were broken by split errors and merge errors (more rare) were corrected to give the entire arbor of neurons with few branches missing in order to provide the connectivity of these neurons to other cells within the volume. Within this network, a small number of neurons form a disproportionately large number of the connections, with the most connected neuron, a chandelier cell, synapsing on 70 other neurons in the network. The number of postsynaptic neurons within the volume receiving synapses from a single proofread neuron ranged from 0 to 70, with a mean of 5. Given the number of neuronal elements in petascale volumes it is not conceivable to proofread more than a small fraction of the data segmented by machine learning. We expect that with improvements both in image data quality and advances in AI, future datasets will substantially improve the automatic segmentation, however we wondered if we could leverage the large number of connected pairs discovered by machine learning to find a substantial fraction of the canonical intracortical circuit. For this analysis we used the more conservative c3 agglomeration which minimized merge errors, albeit with more axon breaks. This approach meant that we were certainly missing a substantial number of axonal connections between neurons in the volume. Nonetheless, we analyzed nearly 30,000 connections between neurons with somata in the volume, all identified by machine learning. We categorized neurons as excitatory and inhibitory based on the presynaptic neuron’s dendrites (smooth or spiny). We identified the layer in which the pre- and postsynaptic somata were located and the layer where the synapse(s) that connected them were located. Because there were no spiny neurons found in Layer 1, we had 6 layers possessing inhibitory cell bodies and 5 layers (layer 2-6) with excitatory cell bodies. We further divided the set of inhibitory cells into interneurons that innervated cell somata and dendrites and the set of interneurons that innervated axon initial segments, which we presume are Chandelier cells ^34, 35^. We drew a circuit diagram based on these connections in which we connected neurons based on both the layer of origin of the pre- and postsynaptic cell and the layer where a synaptic interconnection was found (**Fig. 4J**). All told, we found 96 verified categories of connections based on spot proofreading. The spot proofreading revealed that at least 2/3 of the machine learning (ML)-identified connections were real, although false positive rates varied by connection type (**Supp. Table 5**). Because most of the categorical connections were based on multiple examples, even removing the false positive connections had a small effect on the final number of connection categories. Moreover the connections based on manual proofreading to correct all the axon breaks from 104 randomly chosen neurons (see above) revealed only 2 categories of connections that were missed in the machine learning data (**Supp. Table 6)**. These results suggest that the automatic segmentation even with a conservatve agglomeration that minimizes merge errors at the expense of fewer long axonal segments, still could reveal the vast majority of cell-to-cell connectivity categories in this cortical slab. The connectivity that was revealed connected the majority of cells within and between layers and presents a complex picture of what a full canonical circuit would look like. It is important to emphasize that the approach we took does not provide insight into which types of connections are most or least common. The most numerous connection found was between layer 2 excitatory to layer 2 inhibitory cells (>3000 such connections). This may reflect the short distance needed for axons to connect two cells of the same layer.

### New morphological subcategories of layer 6 triangular neurons

The deepest layer of the cerebral cortex has remained poorly studied compared to more superficial layers for a number of reasons ^36^. Among these is the fact that there is a greater diversity of cell types in this layer, especially in primates ^37^. When cell types are classified by Golgi stains or dye fills, as is often the case in the cerebral cortex, the data may be too sparse to categorize types clearly. This human cortical sample however provides a very large set of neurons that reside in each layer and therefore could potentially be used to reveal categories that were previously unnoticed. As a test of this idea we looked at the so-called “triangular” or “compass” cells that have long been known to reside in layer 6 but are not well understood ^36, 38–40^. These neurons have an apical-going dendrite and are spiny like the more common pyramidal neurons. However these cells each have a second large basal dendrite that projects in a more tangential direction, unlike typical pyramidal cells that have a skirt of several basal dendrites. In layers 5 and 6 we located all the triangular neurons (n = 864, roughly one third of the spiny neurons in these layers in our sample; **Fig. 5A**). The cells we studied all did have a large apical-going dendrite, but unlike typical pyramidal cells, they all also had one large basal dendrite that was projecting in another direction. For some of these cells the large basal dendrite projected directly toward the white matter (green cells, **Fig. 5B**). The majority however, had one large basal dendrite that projected roughly horizontally within the layer. Interestingly, the direction these basal dendrites projected was highly constrained. Almost equal numbers of these triangular neurons fell into two subcategories: those whose large basal dendrite projected roughly orthogonal to the cutting plane, but towards section #1 and those whose large basal dendrite also projected orthogonal to the cutting plane, but in the opposite direction (towards section #5292) (**Fig. 5B**; yellow neurons with basal dendrites projecting towards section #1 or “reverse-going direction”; and pink neurons whose large basal dendrites projected in the opposite or “forward-going direction’’; see <u>video online</u>). Two individual neurons from these two subgroups are shown in **Fig. 5C, D and E**. We measured the angle of each of the basal dendrites of all the triangular neurons. The angles formed a bimodal distribution with peaks for basal dendrites that point along the z-axis in the forward or reverse directions (i.e., where 90 degrees means the basal dendrite is pointing in the z-axis towards slice 0, 270 degrees means the basal dendrite is pointing in the z-axis towards slice 5292, and 0 or 180 degrees means the basal dendrite points within the cutting plane; see **Fig. 5F, G** and **Supplementary Fig. 21**). The fact that the bimodal modes peaked at 60 degrees away from the white matter (i.e., away from the inverted radial direction) (**Fig. 5G** and **Supplementary Fig. 21F**) meant that these two sets of basal dendrites were not parallel to each other. Rather, each subgroup had basal dendrites that projected at a more or less constant angle downward, toward the white matter. For this reason the forward and reverse going basal dendrites formed two sets of parallel basal dendrites that were easily distinguished, one headed forward in the z-axis but tipped slightly towards the white matter and one headed in the opposite direction, and also tipped slightly toward the white matter but at the mirror symmetrical angle (see, **Fig. 5C, D and E**). Notably, there were almost no triangular cells with a large basal dendrite that projected tangentially, that is, in directions that were roughly aligned to the plane of section (gray cells in **Fig. 5G**, **Supplementary Fig. 21 D,E**).

**Figure 5:**
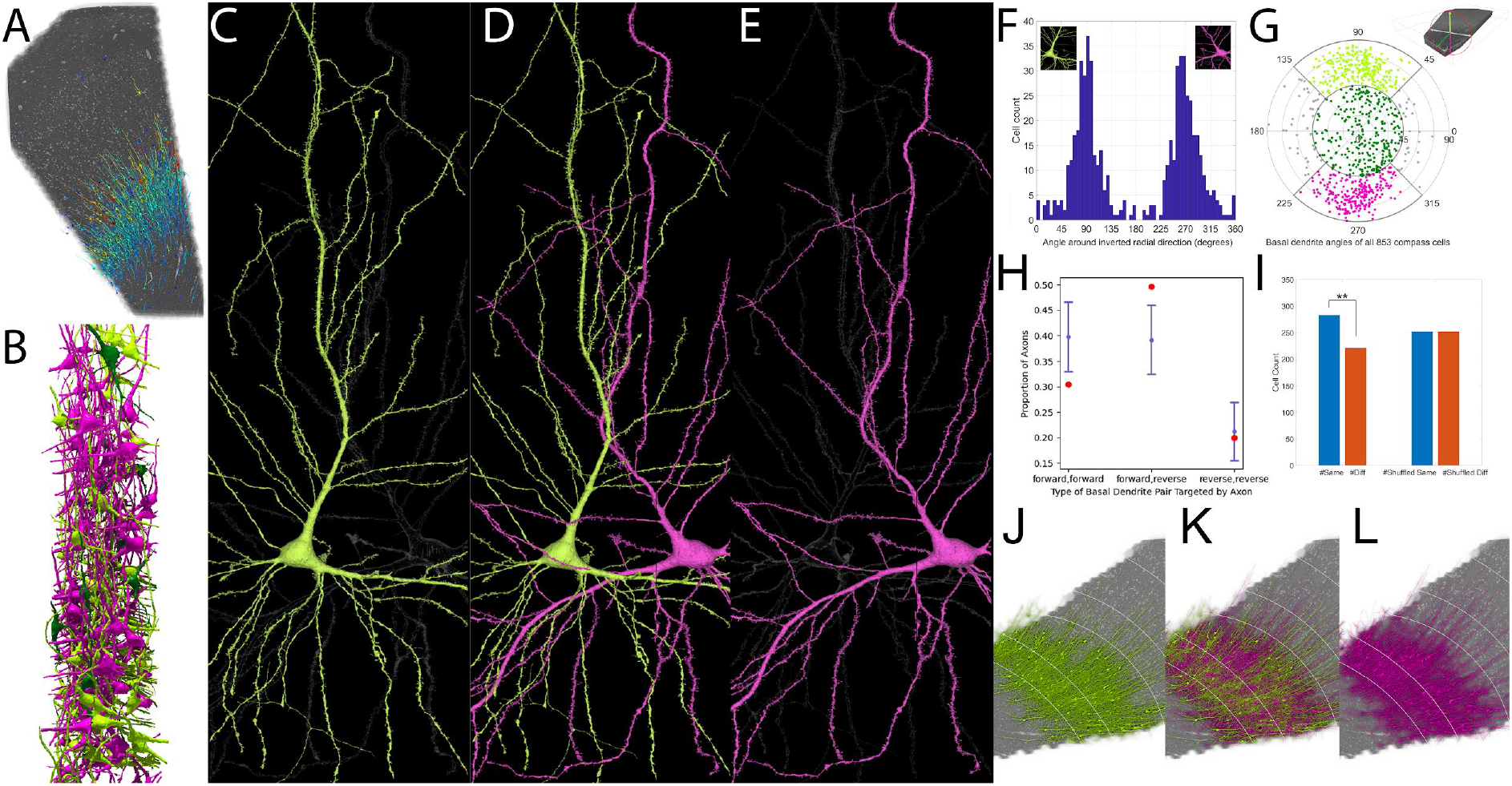
Two mirror symmetrical subgroups of deep layer triangular neurons. **A**: Location of neurons with both one large apical and one large basal dendrite; most are found in layers 5 and 6. **B**: Side view of a selection of these neurons, with apical dendrites pointing either forward in the z-stack (magenta) or in the reverse direction (light green) or straight toward the white matter (dark green). Notice that many of the magenta or light green neurons project their large basal dendrite at very similar angles. **C, D, and E**: Example of two bipolar pyramidal neurons with basal dendrites pointing in opposite directions showing the mirror symmetry in these two subgroups (<u>link</u>). **F:** The histogram of basal dendrite angles (572 cells; light green, magenta and gray cells from G) shows a clear bimodal distribution with peaks around 90 and 270 degrees representing the light green and magenta groups respectively. **G:** The polar plot of the data in F shows peaks of the basal dendrite directions at ∼ 60 degrees away from the inverse radial direction. Light green: 257 triangular cells with basal dendrite pointing towards section 0; magenta: 247 triangular cells with basal dendrite pointing towards section 5292; gray: 68 triangular cells with basal dendrite pointing sideways in the cutting plane; dark green: 281 compass cells with basal dendrite pointing towards the white matter. 11 compass cells excluded (basal dendrite pointing away from the white matter). **H**: For axons innervating two bipolar pyramidal neurons, connections to neurons of the same polarity are overrepresented and connections to neurons of opposite polarity are underrepresented (blue dots and 95% confidence intervals; red dots show expected values). **I**: Statistically significant likelihood that neurons whose large basal dendrite points in the same direction are nearer to each other than expected by chance. **J, K, L**: Anatomical clustering among members of the two subgroups.

Looking at the full cell segmentations of each of these 257 reverse-going + 247 forward-going triangular neurons with tipped basal dendrites (n=504), several other consistent features were notable. First, the apical dendrites were also sometimes tipped so that neurons whose basal dendrites projected in opposite directions sometimes looked roughly mirror symmetrical to each other (see **Fig. 5D**). Second, there was an asymmetry to the location of dendritic side branches originating from both the proximal parts of the large basal dendrite and the apical dendrite. Side branches were far more prevalent on the outer edges of these bent bipolar cells than the parts facing inward. For an example, see in **Fig. 5C**, the dendritic side branches of the proximal apical dendrite facing to the left, and the dendritic side branches of the large basal dendrite projecting downward. The same is true in mirror symmetry for the cell shown in **Fig. 5E**. This was a general feature of these cells.

The different shapes, albeit mirrored, of these two categories of neurons raised the possibility that they are functionally distinct as well. Because we had the complete set of axonal inputs that made synapses to all of these cells, it was possible to analyze if their synaptic input was distinct. We thus analyzed the axonal innervation to 504 neurons with forward- and reverse-going basal dendrites to see if individual axons that innervated more than one triangular neuron showed a tendency to innervate neurons whose basal dendrites projected in the same direction. If there were equal numbers of axonal inputs to the forward- and reverse-going neurons, then one would expect twice the likelihood of individual axons to innervate one forward- and one reverse-going dendrite than 2 forward-going or 2 reverse-going (as is the case in flipping a coin twice yielding twice as often one head one tail (HT 50% of the time) versus two heads (HH 25% of the time) or two tails (TT 25% of the time). Because the total number of axonal inputs to forward- and reverse-going neurons was not equal, the expected outcome, assuming a random process, was finding axons that innervated two neurons with forward-going basal dendrites (FF) to be 32% (32% of the axons that innervated two neurons with large basal dendrites would innervate both forward-going neurons), 49% FR and 19% RR (**Fig. 5H**, red dots). However, the connectivity was significantly skewed for axons to choose pairs of neurons with basal dendrites that projected in the same directions: FF (40% versus 32% expected) or RR (21% versus 19% expected) compared to dendrites pointed in opposite directions, FR (39% versus 49% expected) (**Fig. 5H**, blue dots, p = 3.34 · 10^-14^). In an attempt to understand the origin of this specificity, we divided the axons into excitatory, inhibitory, and inhibitory onto the axon initial segment of the triangular cells. The results were equally significant for the subset of excitatory axons that innervated the basal dendrite neurons as it was for the inhibitory neurons that innervated the dendrites and somata of these cells (**Supplementary Fig. 24**). However, the chandelier interneurons that innervated the axon initial segments of more than one of these cells, did not show a significant preference for co-innervating FF or RR pairs, and in fact, as would be expected in a random model, made most of their co-innervation on RF pairs (**Supplementary Fig. 24**). These triangular cells’ dendritic arbors and axon initial segments overlapped in the volume (see **Fig. 5B and D**). Nonetheless, when we rendered the two subgroups of neurons (forward-going or reverse-going) we noticed that they were not uniformly distributed in layer 6. Neurons whose large basal dendrites pointed in the same direction are nearer to each other than expected by chance (**Fig. 5I**) and the two sets, despite some overlap, appeared clumped (**Fig. 5J, K and L**). Each clump formed a rough radial column approximately 250 μm from the next column of the same subclass. Patchiness of axonal projections to layer 6 have been seen with anterograde labeling experiments in humans ^41^.

### Axonal Targeting

Because we could identify all the synapses on a neuron and the identity of the presynaptic partner of each (**Fig. 6A**), we noted that only a small percentage of the axonal inputs to a neuron established more than one synapse with that postsynaptic cell (**Fig. 6B**). For nearly all cells, the histogram of the number of synapses per axonal input showed a rapid fall off from the prevalent motif where an axon established one synapse with a target cell (> 90%). Two synapse contacts occurred uncommonly, however all neurons showed some two synapse contacts. Even fewer three synapse contacts were observed and typically four synapse connections occurred only 0.001% of the time. One common exception to weak connectivity is the chandelier axon input to axon initial segments of pyramidal cells. Their cartridge inputs sometimes give rise to ten or more synapses from a single presynaptic neuron. On dendrites however, weak connections are the rule. Nonetheless, we noted exceptional outlier connections on dendrites, and more rarely somata where a single axon established eight or even twenty synapses with a single target cell (**Fig. 6C and D**). In the example shown in **Fig. 6C**, an axon establishes nineteen excitatory synapses distributed to several dendrites of an inhibitory neuron. Another example in which an excitatory neuron in the volume establishes nine synapses on an inhibitory postsynaptic neuron is shown in **Supplementary Fig. 19**. This example is notable because of the way in which the synapses are established on both sides of an en passant synapse between an axon and a dendrite (see below).

**Figure 6:**
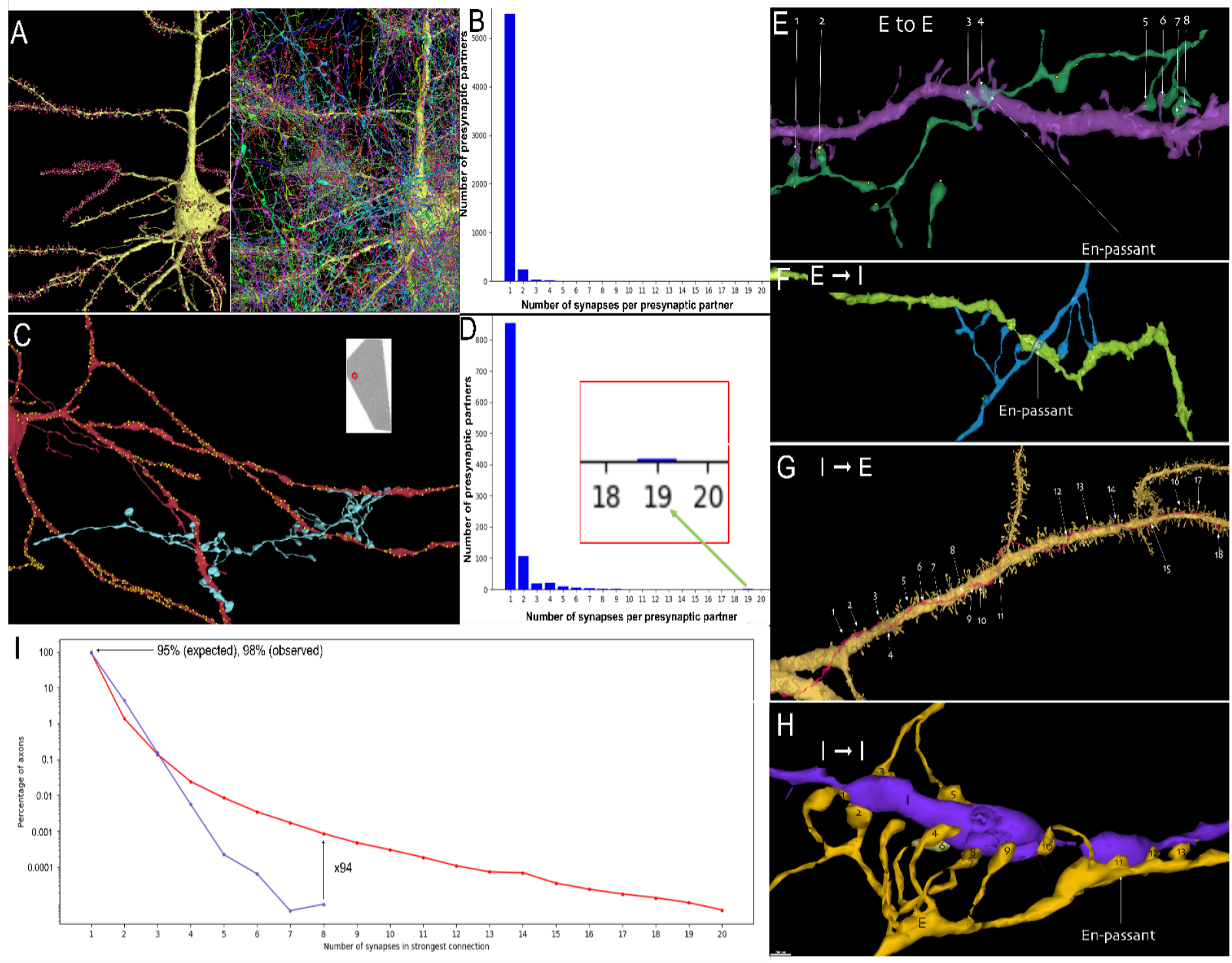
Unusually powerful synaptic connections. **A**: We assayed all innervation to each neuron in the volume. Left panel, a pyramidal cell showing the sites of its incoming synapses (red balls). Right panel shows all the axons that give rise to the incoming synapses. **B**: The vast majority of the 5600 axons innervating this pyramidal cell establish one synapse with it, with a small number of axons that provide two to four. **C:** Occasionally a very large number of synapses exist between an axon and a dendrite, as here where an axon (cyan) provides 19 synapses to three dendrites of an inhibitory neuron (red) in layer 2- see inset. Yellow balls show sites of all incoming synapses. **D**: Histogram shows that most inputs to this cell are weak and the powerful axon is an outlier. **E-H:** Despite their rarity, in such big data there are many strong connections between excitatory and inhibitory neurons. **E:** For example, in this case an excitatory axon (green) forms 8 synapses onto a spiny dendrite of an excitatory neuron (purple). As often was the case, one of these connections is en passant and the rest of the synapses appeared to require directed growth of the axon to contact this same dendrite. **F** An excitatory axon (blue) forms 8 synapses onto a smooth dendrite of an inhibitory neuron (green) again with one en passant connection and the rest apparently requiring directed growth. **G**: An inhibitory axon (red) forming 18 synapses on the apical dendrite of a spiny pyramidal excitatory neuron (yellow). **H**: An inhibitory axon (yellow) forming 13 synapses onto the smooth dendrite (purple) of another inhibitory neuron, including one en passant contact. **I**: A log normal plot showing incidence of axons establishing between 1 and 20 synapses to individual postsynaptic target cells (red line). This incidence far exceeds a model where these same axons establish the same number of synapses but are slightly displaced in space (blue line, see Methods). For all connection strengths greater than 3 synapses, axons show more multiple synapses than expected by chance. For example there were 94-fold more axonal connections with 8 synapses to one target cell than expected if axons establish the same number of synapses on nearby dendrites in a random way.

Such outlier connections were not restricted to excitatory input to inhibitory dendrites (see also **Fig. 6F**). Excitatory axons also occasionally formed strong (multisynaptic) connections onto spiny (excitatory) postsynaptic neurons (**Fig. 6E**). In addition, inhibitory axons made such outlier powerful connections onto the shafts of excitatory neurons (**Fig. 6G**) and onto the dendrites of inhibitory neurons (**Fig. 6H**). Thus both excitatory and inhibitory neurons establish unusually strong connections with both excitatory and inhibitory postsynaptic cells, albeit rarely. Importantly, these strong connections appeared to be a property of the particular pre- and postsynaptic pair: most of the synapses established by the axons that formed outlier connections were typical one-synapse connections (on other neurons) and most of the input to the postsynaptic cells that received these powerful inputs were typical one-synapse weak connections (from other axons). Despite the overall rarity of strong connections, we found that 28% of the 2659 neurons that were well innervated in the volume (i.e., that had at least 3000 axonal inputs onto their dendrites) had at least one input that had 7 or more synapses, raising the possibility that rare powerful axonal inputs are a general characteristic of neuronal innervation in the human cerebral cortex.

Many of these powerful connections shared a common morphological configuration. In some cases the axon co-fasciculated with the dendrite to remain in close contact for tens of microns, allowing it to establish many en passant synapses with the same target cell (e.g., **Fig. 6G**). More commonly however, the axon did not appear to have a special affinity for growing along the dendrite and approached the dendrite, as was typical of axons that made one synapse connections, by forming a synapse at the site of intersection without deviating its trajectory before or after the synapse. Remarkably, in addition to a synapse at the closest point of intersection, these axons sent terminal branches to the same target cell, usually both before and after the intersection. Because many of these axon fragments left the volume without reaching a soma, it was not possible in most cases to decide which side of the intersection was actually “before” or “after”. Nonetheless, the behavior of the axon was the same on both sides: the axon potentially sprouted “up” to establish synapses on the dendrite, and on the other side it potentially sprouted terminal branches “down” to establish additional synapses with the same target cell (see **Fig. 6E, F and H** and **Supplementary Fig. 19**). All of these cases, and many of the thousands strong connections not shown, were suggestive of intentionality, meaning that some pre-postsynaptic pairs had a reason to be far more strongly connected than was typical. The alternative possibility is that, given the thousands of axonal inputs to each of thousands of target cells, outlier results are simply part of the long tail of a distribution. These powerful connections may also be more common than we found given that axon breaks in the automatic segmentation likely had a disproportionately greater likelihood of reducing the number of connections as the length and number of synapses produced by an axon increased. We thus sought a conservative model to simulate the number of axons with powerful connections, based on the actual trajectories of axons and dendrites and the observed properties of axonal branches. The model allowed every simulated axon to form the same number of synapses as it did in the actual data, but now on any of the dendritic branches that came within its vicinity, to match the reach of the actual axons. We were interested to see how often an axon would establish one or more synapses with a single target cell. The results shown in **Fig. 6I**, indicate that this random model of synaptic partnering is inconsistent with the incidence of strongly paired neurons that we found in H01 (p < 10^-10^). This tendency for axons to establish more synapses with certain target cells than expected by chance was found to about the same degree when we analyzed just inhibitory or just excitatory axons. Thus, amongst a large number of exceedingly weak connections, human cerebral cortex neurons receive a small subset of inputs with approximately an order of magnitude more power.

## Discussion

The central tenet of connectomics is capturing both big and small scales: reconstructing individual synaptic connections in volumes large enough to encompass neural circuits ^42^. Our aim in this work was to study the structure of the human cerebral cortex at nanometer-scale resolution within a ∼millimeter-scale volume that permitted seeing all of the cortical layers and some white matter. To observe intracortical connections between six layers of cortex, it was necessary to image a volume encompassing its full thickness, which in humans (including this sample) averages 2.5 mm from the top of layer 1 to the bottom of layer 6 ^43^. Given that the axons of neurons may travel over one hundred microns before making their first synapses ^44^, the other two dimensions of the imaged volume must be wide enough and deep enough to trace intracortical network connections. We found that a full cortical thickness volume equivalent to a cubic millimeter provided us with sufficient tissue. The nanometer scale is required to identify individual synapses and distinguish tightly-packed axonal and dendritic processes from one another ^45^. This big and small requirement necessitated the acquisition of trillions of voxels, and hence more than a petabyte of digital image data. This petascale dataset offers the opportunity to look at the same volume of brain tissue at supracellular, cellular, and subcellular levels and to study the relationships between and among large numbers of neurons, glia, and vasculature. Most importantly perhaps it gives a glimpse into the enormous complexity of the synaptic relationships between many neurons in a slab of human association cerebral cortex.

This “digital tissue” ^46^ is a ∼660,000-fold scale up of an earlier saturated reconstruction from a small region of mouse cortex, published in 2015 ^47^. Although this scaleup was difficult, it was not hundreds of thousands of times more difficult and took about the same amount of time as the previous data set (∼4 years). This means that many of the technical hurdles with imaging and computer-based analysis have improved dramatically over the past few years. This improvement was in large part due to two noteworthy advances: fast imaging owing to multibeam scanning electron microscopy ^48^ and the profound effect of AI on image processing and analysis ^9^. The rapid improvements over the past few years ^49–58^ argues that analyzing volumes that are even three orders of magnitude larger, such as an exascale whole mouse brain connectome, will likely be in reach within a decade ^59^.

Studying human brain samples has special challenges. Fortunately, the quality of the H01 brain sample was comparable to cardiac perfused rodent samples used in the past. This strongly suggests that rapid immersion of fresh tissue in fixative is a viable alternative to perfusion and should be especially useful in human connectomic studies going forward. More problematic is that fresh samples from completely normal individuals are unlikely to ever be available via this neurosurgical route. Although this patient’s temporal lobe did not show obvious signs of abnormality, as stated by the neuropathologist, it is possible that long term epilepsy had some more subtle effects on the connectivity or structure of the cortical tissue. Moreover, it is likely that an epilepsy patient such as the one who supplied this sample was treated (albeit with limited success, hence the surgery) with pharmacological agents that could affect brain structure. Importantly however, the neurosurgical specimen that we obtained was not part of the pathological process per se, it was removed only because it was “in the way” – that is, the surgical procedure could not be done without its removal. Given that patients with successful outcomes after surgery have normal brain function, it is assumed that the procedure pinpointed the abnormality, and the remaining brain is functioning normally. The normal functioning brain would presumably include the incidentally resected tissue. However, only by comparing samples obtained from patients with different underlying disorders may we eventually learn whether this sample is normal. There were some oddities in this tissue that we found, but at present, we cannot decide if they are pathological or just unusual. These include a number of extremely large spines and axon varicosities filled with unusual material (see **Supplementary Fig. 7**).

Another challenge with human brain tissue from the association cortex is that it is probably the location of circuits that were established as a consequence of experience. If memories are stored in this part of the human brain, then it is unlikely that another brain will be similar, in the way that, for example, multiple *C. elegans* nematode brains are stereotyped.^60^ But even in isogenic worms, 40% of the neuron to neuron connectivity was different between specimens. Given the far greater variability in human experience, behavior, memory and genetics, and the fact that humans and other vertebrates have pools of identified neurons classes rather than individual identified neuron types, it will no doubt be challenging to compare neural circuits between brains. This challenge also presents an opportunity: to uncover the physical instantiation of learned information. Even if the circuits differ in their particulars, it is possible that a metalogic for memory can be uncovered by looking at enough data, maybe in the future field of “engramics” ^61, 62^. To be sure, approaches to the profound questions of uncovering the meaning in neural circuit connectivity data are in their infancy, but it would seem to us that perhaps the best stimulus for making progress will be an abundance of actual data -- this petascale dataset is a start.

However, we have come to realize that big data raises big problems that do not have ready answers. Although segmentation is improving rapidly, the automatically reconstructed circuits are far from perfect. The tension between split errors and merge errors means that depending on the aim, one may prefer segmentations that provide fewer split errors or fewer merge errors (we share online two agglomerations that are both these alternatives, c2 and c3). In this sample we used higher level semantic annotations to remove some merge errors, such as trimming astrocytes or axons that were merged to dendritic trees. These higher-level machine learning approaches are likely to be profoundly useful as they become perfected in the years to come. At the current level of automated accuracy, it is impractical to manually correct all segmentation errors in a volume as large as H01. However, when we scrutinized the segmentation errors in this data, they were almost always secondary to flaws in the image acquisition, such as staining artifacts at a critical site, alignment difficulties due to variations in the thickness, or distortions of the sections that were picked up onto tape. We suspect that improved methods of tissue sectioning and staining may help to reduce the number of errors in the future.

One power of connectomic study of large data is that rare events, which might be too uncommon to find in a small sample, are manifest in the larger data size. It has long been recognized that dendritic trees allow neurons to collect information from, as shown in this study, many thousands of different neurons. Because the overwhelming majority of these connections are weak (∼ 99.9% with 1, or at most 4 synapses), it has been assumed that neural processing must occur by integration via spatial and temporal summation of a number of weak inputs that happen to be active at roughly the same time. We found however that amongst the very large population of weak axonal connections, there are a few inputs that are outliers forming ten or even twenty synapses on the dendrites of a postsynaptic neuron. By virtue of their common origin, these multiple release sites assure that one input will synchronously activate a target cell, perhaps strongly enough to be suprathreshold (if excitatory) or able to block activity (if inhibitory). In a much smaller volume of proofread mouse somatosensory cortex we previously saw axons that made as many as up to five synapses on the same dendrite that also did not appear to be a chance occurrence^47^, perhaps implying that sparse powerful inputs may be a general feature of mammalian brains.

A number of years ago, neuropsychologists ^63^ made the case that motor learning occurs in three stages: cognitive, associative, and autonomous. The point being that early on a motor task is hard to do, and takes much cognitive effort, however eventually it occurs with little thought (e.g., learning to use the brake pedal when seeing a red light as a beginner driver, versus the automatic and almost unconscious braking by an experienced driver). Clearly the human brain can reach a decision much more quickly once a task is completely learned. How might that occur? One possibility is to strengthen a pathway so it is less reliant on summation from many sources, as might occur if one input becomes capable of activating a target cell on its own. Interestingly, this is the maturational strategy in the final limb of the motor system ^64, 65^. In young animals (probably including humans) neuromuscular junctions are innervated by many weak motor axons, at which point it is likely that multiple axons need to be activated synchronously to cause reliable muscle fiber twitching. Over time, many of these inputs are eliminated and the remaining axon compensates by adding more synaptic release sites ^66^. The remaining input is strong enough to dependably activate the muscle fiber given the high quantal content of twenty or more release sites. Such strengthening also occurs in the CNS, such as the strong excitatory input of one climbing fiber on an inhibitory postsynaptic neuron, the Purkinje cell. This multisynaptic release also emerges after a set of weaker climbing fiber inputs are eliminated and the remaining axon adds many synapses ^67^. Is it possible that the association cortex permits some inputs to become autonomous drivers of activity by the addition of synapses? If so, such connectivity should be even less common in younger human brains.

The presumption has long been that human neuroscience *must* utilize approaches that are both less direct, and more poorly resolved than those that can be brought to bear on questions in animal models. In particular, it has been assumed that while it might be possible to get complete wiring diagrams and structural cell type analysis of all cells in the cerebral cortex of a rodent, or in the CNS of a fruit fly or a worm, a similar feat would be impossible in a human. However, should the technical barriers with human specimens be overcome, would it not be more relevant to do these same kinds of analyses in human specimens? Would it not be especially significant to do this in specimens from humans afflicted with psychiatric or developmental disorders? The petascale data presented here supports the idea that fine-scale connectomics is a viable path for learning about the human brain.

## Acknowledgments

We thank Matthew Frosch, MD, PhD, Massachusetts General Hospital for kindly providing the human tissue. We are very grateful for all the advice from the multiSEM team at Carl Zeiss that helped us get their device into our workflow. We would like to thank Kathleen Rockland for helping us with relevant literature. We are also appreciative for the careful reading of the manuscript by Suzanne Montgomery. Several students provided help in various aspects of this project; they include: Allen Judd, Elisa Paravino, Ray Jiang, Rachael Han, Peng Miao, Tianxin Lu and Jana Afeeli. We gratefully recognize the generous support from the NIMH for the Conte Center Award: P50 MH094271, The Stanley center at the Broad Institute, and the BRAIN Initiative of the NIH for support from U19 NS104653, U24 NS109102 and UO1 EB026996.

## Data availability

All data is available via the dedicated webpage for this project: http://h01-release.storage.googleapis.com/landing.html.

## Code availability

All custom code used is cited in the text, and is freely available at https://github.com/ashapsoncoe/h01.

## Statistics

For the analysis of spatial clustering of cells in Figure 5, Fisher’s Exact Test was used to establish whether equal-color neighbours were occurring significantly more often than by chance (p = 0.00745, n = 504). For the analysis of forward and reverse facing connectivity in Figure 5, the Chi-Squared Test was used to compare observed and expected frequencies of forward-forward, forward-reverse and reverse-reverse linked pairs of forward and reverse going neurons (p = 3.34·10-14, n = 1180). For the analysis of connectivity strengths in Figure 6, the Chi-Squared Test was used to compare observed and expected frequencies of axons’ strongest connections (p < 10^-10^, n = 79,816,870), but plotted as percentages for clarity.

## Materials & Correspondence

Correspondence and requests for materials should be addressed to J.W.L. and V.J.

## Author contributions

Tissue sample was prepared for EM and embedded for sectioning by DL. Block trimming and initial screening was done by DRB. Sample block was sectioned by RLS. Sections were mounted onto wafers and post-stained by ASC. Raw electron microscopy data was acquired by ASC, RLS and YW. Image quality assessment tools were developed by YW and used to assess images by ASC. Image alignment was performed by A Pope, YW, SW and A Peleg, with fine realignment by MJ. Bridging of re-entry sections was performed by SW, AP, ASC, AF and RK. Ground truth for segmentation was produced by ASC, DRB, AF and RK. Ground truth for semantic segmentation was produced by ASC, NK and DRB. Segmentation and agglomeration was performed by MJ, with further correction of agglomeration by PHL. Ground truth for synapse prediction was produced by ASC, AF, RK, JWC, DA, JL, DW, ZL and HF. Synapse prediction was performed by TB. Ground truth for synapse excitatory vs inhibitory classification was produced by ASC, TB and NK. Synapse excitatory vs inhibitory classification performed by TB. Ground truth for subcompartment classification produced by PL, ASC, AF and RK. Skeletonization and subcompartment classification was performed by PHL. Embeddings produced by SD. Proofreading of neurons performed by ASC, BF, HW, JWC, DA, JL, and LB. Identification of cortical layers performed by ASC and LB. Neuroglancer software developed by JMS. API development and support performed by LL. CREST proofreading program developed by ASC. Extension of VAST to access Google-hosted datasets developed by DRB. Manual identification of cell nuclei performed by ASC, BF and HW. Manual cell soma painting and classification of cells performed by DRB. Analysis of powerful synaptic connections performed by ASC. Analysis of triangular neurons performed by DRB and ASC. Astrocyte distance analysis, type-specific blood vessel distance analysis, and cell density analysis was performed by DRB. Analysis of synaptic network was performed by ASC, JWL and YW. Analysis of inputs to interneurons and pyramidal neurons by NK, YW and ASC. Manual annotation of blood vessels cells and their skeletonization was performed by ES and DRB. Manual reconstruction of whorled axons and identification of anomalous objects and chandelier-chandelier connections was performed by NK. Analysis of distribution of myelin was performed by YW. Manuscript written by JWL, ASC, MJ, AP, DRB, TB, PHL, VJ.

## Methods

### Sample acquisition and preparation

A 45 year old woman with a history of simple and complex partial seizures with occasional generalization, refractory to medical management, underwent surgical ablation of an epileptic focus in her left medial temporal lobe. During the procedure medial temporal lobe tissue containing the epileptic focus, as well as unaffected cortical tissue from the left anterior temporal lobe was removed. A pathological assessment of the excised medial temporal lobe sample showed hippocampal sclerosis, and marked neuronal loss from CA1 and less severe neuronal loss from DG, CA2 and CA3, but no significant pathologic changes were noted in the anterior temporal lobe sample that we reconstructed. The anterior temporal lobe sample when excised was 2.5 cm by 0.8 cm in its longest and perpendicular axes, forming an irregular oval shape that included the full thickness of the human cerebral cortex (see **Supplementary Fig. 11**). Immediately after excision, Matthew P. Frosch, M.D., Ph.D., a neuropathologist, fixed the sample by immersion in cold 2.5% paraformaldehyde / 2.5% glutaraldehyde in 0.1 M Sodium Cacodylate Buffer, pH 7.4 (Electron Microscopy Sciences, #15949) ^68^) and maintained the sample in fixative overnight. The sample was then washed in 0.1 M Sodium Cacodylate and 2 mM CaCl_2_ buffer, trimmed manually and divided into 300 micrometer-thick sub-samples by Vibratome (Leica VT1000S), each then stained with reduced osmium tetroxide-thiocarbohydrazide (TCH)-osmium ^6^, washed in ddH_2_O, stained with en-bloc 2% uranyl acetate overnight at 4°C, dehydrated with 20%, 50%, 70%, 80%, and then 100% ethanol, washed twice in propylene oxide (PO), immersed in a 50:50 mixture of PO and 812 Epon resin (EMbed-812, Electron Microscopy Sciences, #14121) overnight, followed by a 30:70 PO Epon mixture immersion overnight and then 100% Epon immersion overnight. The infiltrated tissue sample was cured at 60°C for 48 hours.

### Sample sectioning

The second Vibratome section down from the original sample surface was selected for electron microscopy imaging. The resin block was trimmed using a 3 mm UltraTrim diamond knife (Diatome, USA) and ultramicrotome (UC6, Leica, Germany) to a rectangular area of 4584 x 1975 micrometers, oriented so that it included all cortical layers from layer 1 down to superficial white matter. 30-40 nm thick serial sections were cut with several 4 mm wide Ultra 45 or Ultra 35 diamond knives using the automated tape collection ultramicrotome (ATUM) system as described in ^47^ at a cutting speed of 0.3 mm/s and collected onto carbon-coated Kapton tape. After 1639 serial sections had been cut, the cutting process became unstable, with sections intermittently breaking into two or more pieces (see **Supplementary Table 1** for per-section details), resulting in cutting being paused after 1695 sections so that the knife could be replaced, as blunting of the knife during the cutting process has been observed to result in unstable cutting ^47^. Upon resuming the cutting, due to the replaced knife being significantly non-parallel to the face of the block, sections were initially composed of one part of the face of the block (termed ‘reentry’ sections), and gradually became ‘full’ sections after thirty further sections had been cut. We estimate that we lost no more than the equivalent of three 30 nm sections in this reentry region. The subsequent alignment process in the reentry region is described below. After a further 66 sections had been cut, cutting remained unstable, as indicated sections which alternated between being thicker and thinner than the desired 30 nm thickness. Cutting was paused and the block rotated 180 degrees, with cutting resumed at 33 nm per section. This resulted in a second series of ‘reentry’ sections of 62 sections duration, after which cutting remained stable for a further 3193 sections, giving a total of 5053 consecutive sections cut and collected onto tape. The tape was manually inspected for evidence of failure to cut a section followed by a ‘double-thickness’ section of 60 or 66 nm thickness and found that 5.8% of sections were ‘double-thickness’, as listed in **Supplementary Table 1**. These sections are deliberately duplicated in the digital aligned stack to maintain a realistic size of the digital dataset, resulting in 5292 layers in the digital stack.

### Estimation of tissue compression introduced during ultrathin sectioning

Comparing electron microscopic images of full ultrathin sections with known pixel resolution to photographs of the trimmed tissue sample in the resin block before sectioning allowed us to estimate the compression factor of ultrathin sections with respect to the embedded block which occurred during sectioning and tape collection. Compression in the direction parallel to the knife edge was negligible (0.997) whereas there was a large compression perpendicular to the knife edge (0.72). This means that sections in the electron microscopic stack are compressed by 27.96% with respect to the tissue in the resin block, but only along the longer axis of the section. The corrected true pixel size in the published aligned image stack is thus estimated as 11.1 nm x 8 nm or 5.55 nm x 4 nm in the full-resolution images. This analysis only estimates size changes between embedded tissue and the EM image stack. It excludes any size changes that may have occurred between the tissue *in vivo* and the processed, resin-embedded sample.

### Wafer fabrication and mapping

The tape holding the sections was cut into strips containing between nine and fifteen sections each and attached via strips of 25.4 mm double-sided carbon tape (Ted Pella, USA) onto either round or square silicon wafers (University Wafers, USA) of either a 4 inch diameter or a 90 mm x 80 mm area, respectively. Each wafer held nine strips of tape, and between 110 (round wafer) and 135 (square wafer) sections. To enhance the signal from cell membranes, each wafer was first plasma-treated for 30 seconds (operating pressure of 6 x 10^-1^ mb, plasma current of 10 mA) to increase its hydrophilicity and then immediately stained with 4% uranyl acetate for three minutes, rinsed with ddH_2_O for thirty seconds three times, air-dried, stained with 3% lead citrate for three minutes, rinsed and air-dried as before, stored overnight under vacuum and mounted on a metal wafer holder with fiduciaries to target high-resolution imaging by the multibeam scanning electron microscope (mSEM). As previously described ^7^, we mapped the position of each section on the wafer, relative to fiducial marks on the stage by using a reflected light microscope to produce a low-resolution (3.57 μm / pixel) optical image of each wafer mounted on the wafer holder, which identified the position of each section relative to the wafer holder fiduciaries. These positions were sent to the electron microscope to guide the stage automatically to each section. Several whole sections were imaged by low-resolution SEM (220 nm / pixel, secondary emission, FEI Magellan microscope), within which cortical layers and a perpendicular axis of apical dendrites and axonal bundles were observed (see **Supplementary Fig. 9** for a representative example). A six-sided polygonal region of interest (ROI) was defined to follow this axis, using distances from the corners of the section as indicated in **Supplementary Fig. 9**. This ROI was superimposed onto each section in the optical image of each wafer using the Zen software package (Zeiss), to target high resolution imaging in the mSEM.

### EM image acquisition

To speed imaging, a wafer attached to the stage was mounted in the multibeam scanning electron microscope (mSEM; Zeiss), the stage was automatically sent to each section, and the scope’s software calculated the local stage movements to image each section. Each section was imaged by scanning 61 overlapping rectangular regions with 61 electron beams simultaneously ^48^, producing 61 image tiles, each 3128 x 2724 pixels, comprising one multibeam field of view (mFOV) that had a length of 108 μm ^48^ (**Fig. 1**). Once one mFoV had been acquired, the mSEM stage moved to an adjacent site within the section to acquire another mFoV. Each image ROI required approximately 700 overlapping mFoVs to image the entire ROI at 4 nm per pixel resolution, with a tile overlap (within mFoV) of 10% and a between-mFoV overlap of between 3% and 10% (see **Supplementary Table 1** for all per-section imaging metrics). Each ROI was imaged with a landing energy of 1.5 kV with each scanning beam at 576 pA, and a dwell time of either 200, 400 or 800 ns per pixel. The data was imaged using secondary electron emission. Typically sections were imaged with a 200 ns dwell time, but if small structures such as the outline of synaptic vesicles, or thin processes could not be seen clearly, then the section would be reimaged at a higher dwell time. At a dwell time of 200 ns and stage settling-time of 0.6 s, acquisition of one ROI takes approximately thirty minutes. Brightness and contrast for each ROI were set to maximize the dynamic range of the images acquired, by maximizing the spread of the histogram of image grey levels without clipping its tails. Prior to imaging each ROI, the mSEM was programmed to determine the optimal focus distance and stigmation settings at 12 or more ‘focus support points’ (FSPs) within the section. If this procedure failed at more than 25% of the FSPs, then the ROI was not acquired, and the procedure was restarted with new FSPs added and failed FSPs removed or moved to other locations. Once this procedure had succeeded at 75% or more FSPs, Delaunay triangulation was used to interpolate a topological map of the ROI, to guide the autofocus of the mSEM during the imaging of the ROI.

### EM image quality checks

After each ROI had been acquired, a suite of custom MATLAB scripts (https://github.com/lichtman-lab/mSEM_workflow_manager) was used for further quality checks. A previously described measurement of image quality ^7^ was plotted for each mFOV in the section, highlighting areas within the section of relatively lower image quality, which enabled the identification of out of focus areas, so that affected ROIs could be reacquired with additional FSPs. A correlation measure between tiles that should overlap with one another, both within-mFOV and between-mFOV, allowed the identification of any ‘gaps’ in the imaged ROI. The tops of tiles were automatically checked for evidence of either insufficient mSEM stage settling time (manifesting as a ‘sawtooth’ appearance), or charging (manifesting as compression of the image). Degrees of mFoV rotation and completeness of image tiles or metadata files were also recorded. If any of these tests indicated errors in image acquisition, that would impact on subsequent image stitching, alignment and segmentation, then the section in question was reacquired. Per section quality check results are listed in **Supplementary Table 1**. In total, 5053 consecutive ROIs and 1830 TB of raw imaging data was acquired.

### Image Alignment, Stitching and Rendering

Stitching and alignment presented particular challenges due to the large section area; the fragmentation of some sequences of consecutive sections at coincident fracture boundaries; and reentry periods where, due to sample tilt during sectioning, some sections interfaced with several adjacent sections rather than single ones. These issues were addressed by the techniques described below. The large size of the dataset, also a challenge, was addressed by parallelizing each step over image tiles or regions of the volume, as appropriate. The overall approach was similar to Saalfeld et al. ^8^.

#### Stitching

To stitch imagery within sections, we obtained point correspondences between overlapping tiles, then optimized tile position estimates to minimize the correspondences’ sum of squared errors. We found correspondences by first matching whole tiles around positions suggested by stage position data (or by preliminary, partial stitch solutions in the case of some sections with poor stage position data and low cross-tile overlap), then matching patches around locations suggested by these whole tile matches. We used CLAHE-enhanced imagery subsampled to 8 nm resolution, patches of 400 x 400 nm, the normalized correlation coefficient as a match measure, and subpixel estimation of correlation peaks. To obtain well distributed correspondences, we attempted matches at successive locations generated by Halton sequences, stopping when finding a desired number of patches per pair, or an excessive rate of match failure. We found that the relative position of tiles within the 61-beam field of view varied during acquisition (perhaps due to variations in section surface height) so we solved translation of each field’s tiles individually, interpolating from adjacent fields in cases of insufficient correspondences (e.g., in low texture regions such as blood vessels). When stitching together whole fields, we found that these could not always be adequately aligned by translations or other affine transforms, perhaps because of deformation of the tissue during image acquisition. After solving for an optimal translation of each field, we modeled each with a triangular elastic mesh and adjusted mesh vertices to further reduce correspondence errors, formulating the problem as a sparse nonlinear system and solving it with the conjugate gradient method.

#### Alignment correspondences

To align sections, we first matched coarse and fine structural features between different sections, then used those to guide searches for patch matches. We sought matches between all pairs of sections up to three sections apart or, within reentry periods where more widely separated sections could still be physically proximate due to cutting angle, up to 200 nm apart. Where consecutive sections were fragmented at coincident boundaries, we also obtained matches to widely separated sections in order to adequately constrain those fragments. Features and matches were obtained from section renderings generated from the previous steps stitching solutions, with CLAHE enhancement. We detected coarse features using the OpenCV SimpleBlobDetector^69^ on imagery subsampled to 2-micron resolution, typically obtaining features at blood vessels and some cell nuclei. We detected fine features using a Google proprietary scale-invariant keypoint detector comparable to SIFT ^70^ on 64 nm imagery. In both cases, we used a SIFT-like feature descriptor to characterize the neighborhood around each feature, and compared those descriptors to identify putative feature pairs. With coarse features, we used RANSAC ^71^ to find a global affine aligning each pair of sections, constraining affines to ones with plausible rotation, scale and skew. However, with fine features, the density of features together with the size and deformation of sections made RANSAC insufficiently selective, so in that case we used JLinkage ^72^, a multi-model variant of RANSAC, to perform a similar function, finding a series of affines each valid in some local region. Next, we performed patch matching at both coarse and fine resolutions. At 8 micron intervals, and in 64 nm resolution renderings, we matched 6 x 6 micron patches in 64 x 64 micron neighborhoods to obtain coarse patch matches, using a distance-weighted average of neighboring fine feature matches to locate each search neighborhood. We then repeated this at 1.2 micron intervals in 32 nm renderings, matching 2.4 x 2.4 micron patches in 16 x 16 micron neighborhoods to obtain fine patch matches, using coarse patch matches to locate neighborhoods. In regions where feature or patch matches could not be found between a pair of sections, but were found between those and other nearby sections, we cascaded transforms to get the needed neighborhood location estimates. Finally, we filtered each set of patch matches for spatial coherence, discarding any that differed significantly from transforms fit to neighboring matches.

#### Alignment transforms

Fixing the position of one central section, we used a random subset of patch match correspondences to estimate similarity transforms for all other sections, optimizing both correspondence errors and a regularization term maintaining scale near unity. Then, modeling each section as transformed by its similarity transform with an elastic mesh, we adjusted mesh vertices to reduce correspondence errors. We performed this step first with coarse meshes and sparsely sampled correspondences, followed by progressively finer ones and larger fractions of available correspondences, to a finest mesh resolution of 8 microns vertex spacing. In later steps of this process, we subdivided the volume into progressively smaller, overlapping segments of contiguous sections and solved these independently, while keeping their boundary sections fixed, reducing alignment problems to a tractable size. The final step repeated the penultimate one, while using staggered segment boundaries to improve continuity.

#### Rendering

We classified each whole tile as to whether it primarily depicted tissue as opposed to resin or tape. For most sections, this classification was done by a random forest classifier trained on manually labeled tiles characterized by statistics of intensity and spatial frequency. For some sections within reentry periods, tile classification was done by measuring the entropy of the spatial frequency spectrum of ORB keypoints ^73^, selecting tiles where that measure exceeded a threshold, and classifying as tissue the morphological closure of the largest connected component of such tiles. Finally, for selected sections throughout the volume, tissue tiles were identified manually by drawing section-bounding polygons. Classification results were used to exclude non-tissue tiles from the volume.

We rendered a 3D volume of the aligned tissue tiles with each triangle of elastic mesh determining an affine transform of pixels within that triangle. We used CLAHE enhancement and bicubic interpolation. Where multiple tiles had content for one pixel, we selected the tile with the closest center. In general, each tissue section provided content for one or two layers of the volume, depending on the estimated thickness of that section. However, within reentry periods, where the change in cutting angle resulted in some sections filling their volume layers incompletely, we completed those layers by copying in voxel values from adjacent layers. This replication was limited to copying voxels identified as tissue, to ones identified as not tissue (i.e., resin or tape).

#### Fine-scale alignment with optical flow

In order to address remaining misalignment (such as systematic drift and section jitter), we used the section-to-section optical flow regularized with an elastic mesh to realign the complete dataset, with the results of elastic alignment as its initial state. We computed an optical flow field between every section and its two predecessors using cross-correlation. The flow vector was estimated by identifying the peak of the correlation image computed between 160 x 160 pixel patches extracted from the same XY position in the two sections. The patch centers were distributed over a regular grid with 40 pixel spacing. In addition to the flow vector, we recorded the ratio of the peak height, and the minimum correlation image intensity within a 5 pixel radius region around it (“sharpness”). If more than one peak was detected in the correlation image, we also recorded the “height ratio” of the two largest peaks. The flow field estimation procedure was performed over data at 8 x 8, 16 x 16, 32 x 32, 64 x 64, 128 x 128, and 256 x 256 nm^2^ pixel size, computed with area-averaging.

At every downsampling level, we filtered the flow field for “local consistency” by invalidating entries with: height ratio lower than 1.6, sharpness lower than 1.6, absolute flow magnitude larger than 40 pixels, or an absolute deviation from the 3 x 3 window median of more than 10 pixels. We then upsampled all flows to the highest resolution, and attempted to replace invalid flow entries with values of a lower resolution flow field at the same XY location, with preference given to estimates obtained at higher resolutions. The reconciled flow field was then filtered once more for local consistency as described above. The reconciled flow field might still contain invalid entries, either due to consistency filtering, or failure to find a valid value in the lower resolution flow fields.

We modeled every section as a spring-mass system of Hookean springs ^74^, with nodes of the springs positioned at the grid used for flow estimation, and nearest neighbor and next nearest neighbor nodes within every section connected together. Sections were optimized sequentially. Valid flow vectors were used to connect the optimized section with two previous sections with 0-length springs with a spring constant 10x lower than the in-plane springs ^8^. We treated this system as a set of damped harmonic oscillators at critical damping, and integrated it in time with the velocity Verlet scheme ^75^ until the maximum velocity magnitude for all nodes fell below a threshold value of 0.01. Optimization proceeded simultaneously towards lower and higher *z* coordinates, starting with *z* = 1900 as the anchor section.

In this procedure, significant imaging artifacts or missing sections can cause mesh distortions which propagate through the stack. To avoid this problem, we manually reviewed downsampled versions of all sections in the dataset, and decided to mark 287 single sections as invalid (i.e., both preceding and following sections were valid), as well as 96 sections within contiguously invalid blocks of two or more (up to nine) consecutive sections. The invalid sections were not included in the optimization, and instead the flow field was computed between the valid sections directly preceding and following the invalid block. Invalid sections were then optimized separately in a second pass, at which point all valid sections were held fixed. A representative example of a corrected region of misalignment is shown in **Supplementary Fig. 12.**

### Semantic segmentation

#### Valid tissue detection

We used a simple heuristic filter to detect “out of bounds” (OOB) regions of the dataset which did not contain neural tissue. First, we computed a high-pass filtered version of the imagery as *I*_hp_ = *I* + min_3×3_(255 - *I*), where *I* is the pixel intensity of the CLAHE-normalized data downsampled to a pixel size of 64 x 64 nm^2^ and min_3×3_ is the minimum function, applied convolutionally with a kernel size of 3 x 3 pixels. We then downsampled *I*_hp_ to 320 x 320 nm^2^ pixel size, filtered the results with a 10 x 10 pixel uniform filter, and binarized the results at a threshold of 230. To form the final OOB mask, we computed the in-plane connected components of the binary mask, and removed any components smaller than 10,000 pixels.

#### Voxel-wise classification

We trained four convolutional networks to perform four separate semantic segmentation (i.e., voxel-wise multi-class prediction) tasks: 1) tissue classification, 2) neurite type classification, 3) myeloid body detection, and 4) data irregularity detection. The tissue classification model operated on the aligned data at 64 x 64 x 66 nm^3^ voxel size, but all the other models used the aligned data at 32 x 32 x 33 nm^3^ voxel size. The architecture of the networks, training hyperparameters, and class balancing were the same as in Januszewski et al. ^9^, with all networks having a 65 x 65 x 65 voxel field of view (FOV) at their operating resolution. The tissue classification model was trained with random 3D rotation augmentation in addition to the reflection and axis permutation augmentations used for the other models.

The tissue classification model classified every voxel as neuropil, nucleus, myelin, blood vessel, or fissure - see **Supplementary Fig. 13** for a representative example. “Fissures” denoted unusual cylindrical-shaped regions of the dataset in which it was impossible to distinguish individual neurites. We hypothesize that these regions originated from physical damage during tissue resection. The dataset contains five distinct fissure regions, two of which penetrate the whole stack, while the remaining three are more superficial. To speed up annotation, the myelin class included both the myelin sheath and the axon it wrapped, unlike prior work ^9^. Ground truth annotations were collected by manual mask painting in a web-based tool (“Armitage”). 17,361,918 voxels were annotated as neuropil, 2,731,823 as nucleus, 2,702,242 as blood vessels, 1,529,997 as myelin, and 25,678,685 as fissure.

The neurite classification model classified every voxel as axon, dendrite or glia. Ground truth data was created by selecting neurite fragments in a partially agglomerated neuron (instance) segmentation of the dataset. The neurites were then skeletonized, and the nodes of the skeletons served as the FOV centers for the training examples of the network. In total, we manually identified 887 axon fragments (298,250 points), 71 dendrite fragments (307,339 points), and 75 glia fragments (810,480 points).

The myeloid body classification model performed voxel-wise binary classification. Ground truth data was created by manually annotating segments in the base neuron (instance) segmentation of the complete dataset as myeloid (2,001) and not myeloid (2,662). All voxels belonging to the manually selected segments were used as FOV centers for training the classifier (myeloid: 6,696,491, not myeloid: 5,743,067,192).

The data irregularity model classified every voxel as regular or irregular. Irregular voxels were deemed to be those corresponding to acquisition artifacts, such as dust, knife marks, or partially missing sections. Ground truth for this model was collected through manual annotation, using a procedure similar to that used for the tissue classification model. 15,541,762 voxels were manually annotated as regular, and 13,396,821 as irregular.

### Neuron and glial segmentation

The dataset was segmented with Flood-Filling Networks (FFNs), with scale- and seed-order over segmentation consensus ^9^. We trained three separate FFN models, operating on 32 x 32 x 33 nm^3^, 16x 16 x 33 nm^3^ and 8 x 8 x 33 nm^3^ data. The 16 x 16 x 33 nm^3^ and 8 x 8 x 33 nm^3^ models used an FFN model architecture identical to that in Januszewski et al. ^9^: Eight residual modules, an FOV size of 33 x 33 x 17 voxels and an 8 pixel step size in-plane and 4 pixel step size in the axial direction. The lowest resolution 32 x 32 x 33 nm^3^ model used a slightly modified architecture: twelve residual modules, an FOV size of 33 x 33 x 33 voxels, and an 8 pixel step size in both in-plane and axial directions. The models were trained with 375 Mvx (32 nm), 1,285 Mvx (16 nm) and 6,142 Mvx (8 nm) of ground truth data created by human annotators through de novo voxel-wise painting or correction of automatically generated candidate segmentations ^76^.

Segmentation proceeded in two stages. First, we created a “base segmentation” that was optimized to minimize the frequency of merge errors. Then, we applied a multi-step agglomeration procedure to reduce split errors, while keeping the rate of false mergers at an acceptable level.

#### Base segmentation

The base segmentation was created by first segmenting the EM imagery downsampled to 32 x 32 x 33 nm^3^ and 16 x 16 x 33 nm^3^ voxel size with FFN models trained for these respective resolutions. Segmentation results for both forward and reverse seed ordering were combined with an over segmentation consensus procedure that produced a new segmentation with smaller segments formed so that all voxels contained in a consensus segment belonged to exactly one segment in both the forward and reverse input segmentations ^9^. We then removed any objects smaller than 100,000 voxels from the 32 nm segmentation and upsampled it to 8 x 8 x 33 nm^3^, and segmented the empty areas with the 8 nm FFN model. Finally, we computed the over segmentation consensus between the segmentations from all three resolutions to build the overall base segmentation.

During FFN inference, we used FOV “movement restriction” in areas where the magnitude of optical flow (see ‘Fine-scale alignment with optical flow’) exceeded 8 pixels (at 32 nm), 6 pixels (at 16 nm), or 4 pixels (at 8 nm). The FFNs were also prevented from creating seeds in voxels where the mean intensity of the EM images within a 7 x 7 in-plane window centered at the voxel was lower than: 100 for the 32 nm model, 80 for the 16 nm model, and 130 within a 14 x 14 window for the 8 nm model. This procedure prevented segments from being initiated within myelin, lipofuscin, or other electron-dense structures.

We also excluded (“masked”) various regions in the dataset and prevented the FFN from performing inference in FOVs centered in these areas. Masked areas included: 1) 135 largest segments created by binarizing the “blood vessel” and “fissure” predictions of the tissue classifier and computing their connected components, 2) regions in which optical flow alignment mesh nodes connected to springs whose relative horizontal or vertical compression or extension exceeded 20% after optimization, and 3) regions in which more than two sections (not necessarily in consecutive order) were missing (as determined by the OOB mask) within a 17-section window centered at the FFN FOV.

### Agglomeration

#### Pairwise resegmentation

We used FFN resegmentation to agglomerate objects from the base segmentation as described in Januszewski et al. ^9^. Briefly, we evaluated segment pairs (A, B) selected due to their spatial proximity, and performed FFN inference within a small subvolume centered on the pair of original segments, creating temporary segments A* and B* in the process. These were then used to compute: the recovered voxel fractions (f_AA_, f_AB_, f_BA_, and f_BB_, where f_AB_ is the fraction of B found in A*, and so on), the Jaccard index J_AB_ between A* and B*, and the number of voxels contained in A* or B* that had been ‘deleted’ (i.e., during inference their value in the predicted object mask fell from > 0.8 to < 0.5) during one of the runs (d_A_, d_B_). Criteria based on these scores were used to include some segment pairs as edges in the agglomeration graph, as outlined in **Supplementary Table 2**. A subset of segment pairs for which an artifact was detected by the data irregularity detection model within five sections of the pair center point were reevaluated with the voxels classified as irregular replaced by those from the preceding section (type “artifact” in **Supplementary Table 2**).

#### Endpoint resegmentation

Since pairwise resegmentation can fail to connect segments which are heavily fragmented or separated by more than a few unsegmented (background) voxels, we also skeletonized the base segmentation with TEASAR ^77^ and identified all skeleton nodes of degree 1 (“endpoints”). For each base segment A, and for all its endpoints we then ran FFN inference within a subvolume centered at each endpoint, seeding from locations originally covered by A, and generating a new segment Q. When Q covered at least 60% of the voxels of segments A and B within the corresponding subvolume, we considered (A, B) as a candidate agglomeration graph edge. The candidate edge was accepted if there was another candidate edge (B, A), i.e. generated from an endpoint of segment B.

#### Ensembling

In addition to evaluating agglomerations for selected segment pairs and endpoints, we also computed agglomeration candidates based on ensembling multiple base segmentations. Specifically, we segmented the complete dataset with twenty snapshots of the FFN network weights saved during training. This resulted in twenty segmentations, which we compared to the base segmentation by computing the number of voxels covered by both a base segment A and a segment A’ from an alternative segmentation. A’ was considered to “match” A if the number of overlapping voxels exceeded either 10,000, or 1,000 voxels when they corresponded to at least 50% of all voxels of A. When an alternative segment A’ matched two segments A and B from the base segmentation which were also spatially proximal (i.e. we could compute a decision point for it, as used in FFN agglomeration described above), we generated a candidate edge (A, B) for the agglomeration graph. An edge was accepted if it was generated by a sufficiently large fraction of the alternative segmentations (see **Supplementary Table 2).**

We used the tissue classification model to determine the dominant type for every base segment (supervoxel). Segments for which the fissure class was dominant, as well as those created by the 8 nm FFN model for which myelin was dominant, were excluded from all stages of agglomeration. Segments with a dominant myelin classification were excluded from agglomeration stages involving the 8 nm FFN model, regardless of the origin of the segment. We also converted the predictions of the myeloid classification model into individual instances by computing the connected components of the binary predictions. We excluded from agglomeration all base segments that: overlapped with at least 1,000 voxels of a myeloid segment, were no larger than 1,000,000 voxels, and 90% of their voxels overlapped with a myeloid segment.

Wherever further processing required establishing a total order within the agglomeration graph, edges were sorted in descending order of max (f_AB_, f_BA_) within the stage from which they originated, and stages were ordered as listed in **Supplementary Table 2**. The agglomeration graph resulting from the preceding steps, as well as merging segments completely contained within other segments, had 1.7B edges. We denoted the agglomerated segmentation 20201123b. An earlier version of the agglomeration graph which did not include stages of type “ensemble” and “artifact” as listed in **Supplementary Table 2**, or myeloid segment filtering as described above, was denoted “20200916” and used for some of the analysis requiring lower false merge rates.

### Soma and fragment type separation

In order to create a database of cell bodies within the volume, we first binarized the “nuclei” predictions of the tissue type classifier, and computed the connected components of the resulting binary mask, forming an instance segmentation of the putative nuclei. These instances were then reviewed by human annotators in order of descending volume in a web-based visualization tool showing 3D meshes and 2D cross sections of the segmentation and EM data (“Neuroglancer”). Annotators classified nuclei segments as correct, incorrect, or merged. In the “merged” case, point markers were placed to indicate the approximate centers of the individual nuclei. In addition, we manually painted masks for all somas on every 128th section at a downsampled (512 x 512 nm^2^) resolution. These annotations were then converted to points, and reconciled to build a set of 48,682 cell body center locations. Finally, we identified large segments near the z-boundaries of the volume that were initial regions of axons or dendrites whose cell bodies were not contained within the volume itself. Specifically, we reviewed all segments not previously annotated as a cell body and having a cross section of at least 1,000 pixels at 32 x 32 nm^2^ pixel size at z = 250 and z = 5100 (close to the boundary, but also far enough from the boundary such that at least some large fragments were likely to continue deeper within the volume). Neurites which could be identified as likely having their soma outside of the EM volume (based on spatial orientation and branching pattern) were converted to a point annotation located at the centroid of its cross-section at z = 250 or 5100. These points were added to the cell body center annotations, bringing the total number of annotations to 53,395.

We then computationally filtered the agglomeration graph to enforce that segments corresponding to distinct annotated cell body points remain in separate components. Specifically, when computing connected components we scanned the list of edges sorted in descending order of the agglomeration scores, adding them sequentially to a disjoint-set data structure, and discarding any edges which would cause the annotated objects to be merged together. 0.6 M edges were removed as a result of this enforced soma separation procedure.

We also associated every segment with a per-class (axon, dendrite, glia) voxel count according to the predictions of the neurite type classification model, excluding any voxels identified as nuclei by the tissue type model. The voxel counts were updated to reflect the merge decisions between the segments as the agglomeration graph was processed. Wherever an edge would merge two objects containing at least 100,000 voxels each, and 80% of one were classified as glia and 80% of the other as non-glia, the edge was removed from the agglomeration graph. During this process we maintained two sets of connected components and their voxelwise classifications -- one containing all the edges in the agglomeration graph, and the other containing all edges with the exception of those involving a segment previously identified as containing a cell body. The second set of connected components was necessary to track the connected components of neurites specifically, as the segments containing the soma often also contained parts of its dendritic branches and the axon, and as such could not be associated with an axon/dendrite type. We applied the same splitting rule (100,000 voxels, 80% of the same type) between axons and dendrites using the neurite-specific set of connected components. 3.7 M edges were removed due to this neurite type separation procedure.

We acknowledge that the resulting set of cell somata still contains some merge errors. In comparison to the manually annotated cell somata, which were completed afterwards, we found that in the C3 version of the automatic segmentation there are 1345 segments which contain more than one cell body, with a total of 2936 manually annotated cell bodies in those segments (504 of which are neurons). Frequently this includes small glial satellite cell bodies merged with a larger neuronal cell body, and sets of smaller glial cells next to blood vessels. 1100 manually annotated cell bodies could not be assigned to a C3 segment because they did not overlap with any C3 segment.

### Skeletonization of segments

We preprocessed the voxel segmentation data to remove any small holes or “island” segments that were completely contained within a surrounding segment. We computed these relabelings at 32 x 32 x 33 nm resolution on blocks of 256 x 256 x 256 voxels with overlap of 128 voxels in each dimension. First, we computed connected components on all background labeled voxels within the block and assigned each background component a new unique object ID. Next, we determined the set 𝔼 of all “exterior” objects, i.e. those that touched a face of the block. Finally, we built a 6-connected region adjacency graph (RAG) between all object labels, and found all object pairs A, B for which all paths from A to 𝔼 in the RAG passed through a single “surrounding” object B. The surrounding objects B correspond to the articulation points of the RAG. Relabeling the voxels of each A with the ID of the corresponding B fills all holes and islands that are completely surrounded within the block, including in cases where there are multiple A’s per B.

We then skeletonized the voxel segmentation via automated block-wise TEASAR ^77^, based on the Kimimaro implementation ^78^. Kimimaro supports running with non-overlapping blocks via special handling of the block faces, but we found that running with overlapping blocks yielded smoother block transitions. We skeletonized at 32 x 32 x 33 nm resolution on blocks of 512 x 512 x 512 voxels with overlap of 128 voxels in each dimension and invalidation scale 3.0, and then clipped the skeletons back in overlapping regions. This yielded a set of unconnected skeleton fragments for each segment ID.

To reconnect fragments, we then iterated through all fragment endpoints and connected each endpoint to the nearest skeleton node within the set of other fragments for the same segment ID, provided the distance was within 1.5 μm. To avoid excess / spurious reconnections, the candidate connections were ordered from shortest distance to longest, and the set of still-unconnected fragments for the same segment ID was updated at each iteration. Skeletons were then eroded back from endpoints by 100 nm, and sparsified to approximately 300 nm node spacing while retaining all branch points and endpoints.

Finally, we made two types of skeleton corrections in order to bias skeleton connectivity to properly reflect the segment agglomeration graph. First, we cut skeleton edges that jumped between widely separated parts of the base segment agglomeration graph (> 3 hops away). This can occur when two branches of the same cell happen to pass close to each other, or when two dendritic spines from the same shaft touch, resulting in a skeleton edge that erroneously connects parts that are spatially adjacent, but are cytoplasmically discontiguous (a “self-merge error”). One weakness in the > 3 hops criterion occurs in areas where image irregularities result in many small base segments; to address this we also kept skeleton edges that connected parts of the agglomeration graph > 3 hops away if the agglomeration graph was confirmed to have a path connecting them via the set of base segment IDs that impinge on a field of view surrounding the skeleton edge.

Second, we aggressively reconnected pairs of skeleton fragments (maximum reconnect distance 10 μm) if their nodes overlapped the same base segment. This largely fixed disconnections remaining after the standard reconnection procedure (see above), particularly in large objects such as somas where the block overlap was insufficient to ensure smooth block transitions. It also fixed most disconnections introduced by the self-merge error cutting procedure. We considered base segments in descending order by the number of distinct skeleton fragments that each base segment overlapped. As above, we sorted candidate connections from the shortest distance to longest and recomputed the still disconnected set of fragments at each iteration. In this case, only nodes that overlapped the same base segment were considered for reconnection, but we did not require that reconnection initiate from endpoints as above. For the skeletonization of volume 20201123c2 (see below), self-merge cutting removed 9,813,220 skeleton edges, while aggressive base segment reconnection added 4,519,438 edges in the final skeletons. For volume 20200916c3 (see below), self-merge cutting removed 7,500,074 edges and reconnection added 3,291,662.

### Cellular subcompartment classification and merge error correction

#### Subcompartment classification

We trained deep networks to classify the cellular subcompartment or cell type for a subset of skeleton nodes (20% uniformly sampled) following the approach of Li et al. ^10^. The model architecture was a ResNet-18 ^79^ with all convolutions extended to 3D and inputs of 129 x 129 x 129 voxels at 32 x 32 x 33 nm resolution centered on each skeleton node. The three input channels comprised 1) CLAHE normalized EM masked by the segmentation, 2) a pre-synapse mask, and 3) a post-synapse mask (see ‘Synaptic site detection’). The model outputs were probabilities for four classes: axon, dendrite, astrocyte, or soma, and models were trained via stochastic gradient descent with batch size 64 for up to 1.5 M steps. The number of unique training examples were 262,395 axon; 1,151,175 dendrite; 1,392,702 astrocyte; and 39,444 soma. For training we class-balanced the examples by upsampling axon, astrocyte, and soma classes to match the number of dendrite examples. In later experiments, two additional classes were added: cilium and axon initial segment (AIS), and the number of training examples were 257,713 axon; 1,151,175 dendrite; 1,392,702 astrocyte; 40,072 soma; 577 cilium; and 4,682 AIS. We upsampled axon, dendrite, and soma to match the number of astrocyte examples, and upsampled cilium and AIS to 10% of the other class counts.

#### Merge error correction

We built on the approach of Li et al. ^10^ to use subcompartment predictions to fix segmentation errors based on the observation that while FFN base segments are largely free of merge errors, occasionally in FFN agglomeration two base segments with inconsistent classes (e.g. axon versus dendrite) are erroneously merged. In brief, merge error correction consists of 1) soma removal, 2) branch consistency calculation, and 3) cut candidate scoring and thresholding. Prior to processing, the agglomeration graph for each object was modified to remove cycles by breaking an arbitrary edge of each cycle. We also simplified on Li et al. ^10^ by considering only the highest probability predicted class label for each node, rather than working with predicted class probabilities per node.

First, for the subset of objects that included somas, we detected the soma and removed it so that the remaining detached branches could be considered independently. For each agglomerated object, we identified the base segment with the largest number of soma predicted nodes and considered this a true soma if the number of soma nodes exceeded four. We then iteratively considered adjacent base segments in the agglomeration graph, and added them to the soma cluster if they also had more than four soma predictions. The resulting soma cluster generally contained the majority of the soma, as well as most proximal dendritic and axonal branches. These soma base segments were then removed from the agglomeration graph, and each remaining disconnected branch subgraph was considered independently.

For each remaining object we took the predominant predicted node class as the proposed object class, and assigned a weighted consistency score as the total number of nodes with that class prediction. Finally, we considered every edge in the agglomeration graph as a cut candidate, and computed the consistency scores of the two new objects that would result from the proposed cut. For a cut to be accepted, the sum of the two new scores had to exceed the original object score by at least five. We found it helpful to increase this threshold to 15 in cases where the predominant class of the leaving object was dendrite or soma, due to an occasional tendency of the prediction model to cluster these predictions at the ends of branches. In this case, the leaving object was defined as the side of the cut further from the soma, or else the side further from an arbitrary base segment if no soma was detected. We also found it helpful to prevent the removal of large axon components if the cut score improvement was only marginal, so we added a criterion that when the leaving object class was axon the number of leaving axon classified nodes divided by the leaving cut score could not exceed 6.0.

We combined 2.9 M suggested consistency cuts computed from the 4-class (axon, dendrite, astrocyte, soma) predictions on agglomeration 20201123b, along with 3.7 M partially redundant suggested cuts from the “Soma and fragment type separation” procedure above, for a total of 6.0 M distinct cuts. Applying these cuts to 20201123b resulted in segmentation 20201123c2. Applying the same set of cuts to 20200916 resulted in segmentation 20200916c3. Because we removed somas first to avoid spurious suggested cuts at the soma / branch interface, we did not fix any agglomeration errors involving the soma cluster of base segments, although methods to address this were proposed for songbird data ^10^. We did not distinguish between myelinated and unmyelinated axons in the subcompartment predictions, however that information can be useful in biological analyses and is available from the myelin mask described in “Semantic segmentation” above. Therefore we post-processed the subcompartment predictions for any skeleton component that entered the myelin sheath for more than approximately 3 μm consecutively by incrementing their predicted node class labels by 1000.

#### Subcompartment rendering

To produce a volumetric rendering of the subcompartment classification, we first ran topology-preserving erosion on all segments at 64 x 64 x 66 nm resolution. We then computed marker watersheds within each segment, with seed positions at each subcompartment classified node, and assigned voxels within each watershed to the corresponding subcompartment. Because the subcompartment axon class label conflicted with the conventional volumetric background label (0), we incremented the rendered subcompartment classes by 100.

#### Spine detection

We trained a separate network to detect dendritic spine subcompartments, using the same configuration as described for “Subcompartment classification” above. For spine detection, the training data consisted of 33,753 positive examples from pyramidal cell dendritic spines, and a total of 2,628,949 negative examples, comprising 1,976,109 examples from axon, soma, or astrocyte; 578,439 from pyramidal dendrite shafts; and 74,401 from non-spiny dendrites. To detect spines, we ran this network on all skeleton endpoint nodes (leaf nodes with exactly 1 impinging edge) for segments with at least 5 dendrite labeled nodes in the 20200916c3 subcompartment classification.

### Embeddings

We trained a ResNet-18 ^79^ with all convolutions extended to 3D an input field of view of 129 x 129 x 129 voxels at 32 x 32 x 33 nm resolution to produce embeddings for volume locations centered at skeleton nodes. The only input channel was the CLAHE-normalized EM masked by the segmentation. Specifically, the input channel only contained the EM values where the segmentation matched the neuron id, all other values were set to 0. The model output was a 32 dimensional vector. The network was trained following the SimCLR framework ^80^ using contrast and brightness augmentations as well as shifts of the field of view. We used a batch size of 256 pairs and trained the network for 300k steps.

After prediction, we sampled ∼130k skeleton nodes from segments with more than 150 skeleton nodes for dimensionality reduction. From each segment we sampled embeddings from 2% of the skeleton nodes. We next fit a UMAP ^81^ on the sampled embeddings and applied it to all embeddings.

We used these embeddings to distinguish microglia and oligodendrocyte precursor cells (OPCs) in the dataset. First, we identified a number of microglia and OPCs manually by their shape and ultrastructural features (36 microglia, 21 OPCs). Then, we trained a linear classifier to label local embeddings as either belonging to microglia (N=29,536) or OPC (N=25,035). We tested the model’s performance by training on randomly selected 75% of the cells and testing on the remaining ones. The model achieved a mean accuracy of 0.89 (100 sampling rounds). We inferred cell labels for 6,280 cells identified as either microglia or OPC by averaging the probabilities of the local classifications and thresholding at 0.6 for each class. This left 1,058 cells unclassified. Additionally, we excluded 1,778 cells due to large false mergers as identified in the embedding space. In total, our model identified 2,049 microglia, 1,395 OPCs and left out 2,836 cells.

### Manual cell body labeling and classification

To enable efficient manual labeling of all cell bodies (and nuclei of blood vessel cells), VAST ^76^ was extended so that image data could be streamed in on demand from the Google Brainmaps server, and functionality was added to let users quickly jump between specific sections at large intervals while showing intermediate sections during the transitions. Inspection of the EM images at different resolutions showed that the lowest resolution at which cell bodies could be reliably identified in XY sections was mip level 4 (128 nm per pixel). Since the smallest cells in the dataset have a cell body around 6 micrometers in diameter, we chose a Z-stepping of 128 sections (4.2 micrometers) to minimize the number of sections that needed to be painted and at the same time to minimize the chances of missing small cell bodies.

The manual labeling was done in VAST using a screen pen tablet (Wacom Cintiq 13HD, Wacom Inc.). The initial labeling was done cell-by-cell in the following way: Starting from specific XY sections, any unlabeled cell body cross-section which was spotted (nucleus visible) was given a new label ID. It was then painted in that section and also in all other multiple-of-128 sections in which its nucleus was visible, and beyond to capture the whole cell body, scrolling forward and backward through the stack in 128-section steps. This ensured that each cell body painted received exactly one ID and was not erroneously recounted in a different section.

Once the whole dataset was traversed in this way and no unpainted cell bodies were spotted easily, the tissue sample was visually scanned for any unlabeled cell bodies in a systematic way. A XY grid was overlaid over the complete dataset and each grid cell was systematically visually traversed in Z (**Supplementary Fig. 10 A, B**). Any unpainted cell body was added to the list. Each verified grid cell was marked. This should ensure that almost all cell bodies were found. If there are any missing cells, they are likely small glia which only show up in a single painted section or may in rare cases fall between painted sections.

Once all cell bodies were identified and listed in VAST, we classified them manually as neurons or glia of different types based on their appearance in the EM sections and their 3D shape. Criteria used from EM images included cell body and nucleus size, cell body outline, cytoplasm shading, and appearance of the endoplasmic reticulum. For glial cells, the appearance in the EM section is sufficient to identify astrocytes and oligodendrocytes reliably. We did not attempt to manually separate microglia and oligodendrocyte precursor cells, which have only subtle differences in their arborization and ultrastructural features. For neurons, it was necessary to verify/correct their classification by displaying their 3D models in Neuroglancer (**Supplementary Fig. 10 C**).

### Manual labeling of blood vessels and their nuclei

As for cell bodies, XY cross-sections of blood vessels were initially manually painted in VAST at mip 4 (128 nm per pixel), every 128^th^ section. Different label IDs were used for different parts of the blood vessel network in such a way that individual labels (segments) do not contain branching (linear pieces of the network; new color introduced at branch points). To ensure that no branches were missed, each segment was initially tagged as ‘unfinished’. The tag was only removed once inspection of the segment showed that all branches coming off of it were also labeled. Additional inspection of regions devoid of capillary labels and comparison to automatic blood vessel identification revealed some missing blood vessels, in particular large arteries, which were disconnected from the local capillary network. These were added to the segmentation. In a second step the blood vessel labels were augmented by adding intermediate sections semi-automatically to get to every 64^th^ section in Z, followed by manual correction and specific labeling of thin blood vessel branches at even higher resolution.

Based on the linear blood vessel segments, nuclei of blood vessel cells were identified and painted in VAST at mip 3, every 32^nd^ section. Blood vessel cell bodies have more complex shapes than their nuclei, making nucleus labeling a good alternative to enumerate and locate these cells. Blood vessel cells were first classified as endothelial cells and pericytes, followed by sub-classification of pericytes into further cell types based on location with respect to the blood vessel lumen and basement membrane, as well as their appearance in EM. Circulating immune cells were located inside the blood vessel lumen. Perivascular lymphocytes appear similar to circulating lymphocytes but reside within the basement membrane. Perivascular macrophages were defined by their location in the outer layer of basement membrane and abundance of large granules. The difficulty in distinguishing fibroblast-like pericytes from perivascular macrophages resulted in a small group of cells being marked as undecided. Fibroblast-like pericytes were defined by the location surrounding smooth muscle cells and lack of large granules. Smooth muscle cells were separated from pericytes based on their location and slightly larger cytoplasm.

### 3D rendering of cells

Projection images (**Fig. 3**) were rendered directly from volumetric (voxel) data in Matlab (The Mathworks Inc.) using VAST and the Matlab script VastTools. The method produces parallel projection images and uses a simple simulation of light occlusion to generate shading. Other images of cells were either generated from Neuroglancer directly, or were rendered in 3dsMax (Autodesk Inc.) using 3D meshes generated from voxel data through VAST and Matlab (The Mathworks Inc.).

### Synapse identification

#### Synaptic site detection

To identify synaptic sites, we trained a classifier based on the UNet architecture described in Çiçek et al ^82^ to label three classes: background, pre-synaptic, and postsynaptic (see **Supplementary Fig. 14** for representative example). The model used the same three-down/three-up architecture as described in the paper, with initial dimensionality of 32, and a multiplicative factor of 2 for each down and up stage.

Ground truth labels were generated by having trained annotators volumetrically label each pre- and postsynaptic site on both sides of the both sides of the synaptic cleft, for each Z slice where the synaptic density was visible. Six regions of interest were selected (one from each cortical layer) of size [600, 600, 3300] voxels at a resolution of 8 x 8 x 33 nm. The ROI shape was chosen to both allow annotators to observe the entire XY FoV on a single screen to minimize missed annotations, while capturing as much variation in neuropil as possible in Z. Only chemical synapses were labeled, with particular care taken to ensure that any synaptic sites that were nearly parallel with the cutting plane were also labeled. In total 6434 synaptic sites were annotated from 3217 unique synapses across the six ROIs.

Training examples were balanced between positive examples centered on the synapse, and negative examples corresponding to locations where there were no pre- or post-annotations within the FoV. During training, we performed data augmentation, including permutations of X and Y, and reflections along X, Y, and/or Z. The input FoV was [403,403,42] voxels centered around the example points, corresponding to an output size of [191,191,14]. Due to the network’s large memory requirement, a batch size of 2 was used, with batch normalization applied during training. Due to the relative sparsity of synaptic annotations compared to the total number of voxels in the volume, positive pre-/post-annotations were weighted 2x compared to background classifications in the loss calculation. The network was trained to 51 M steps at a learning rate of 10^-5^ using asynchronous stochastic gradient descent, using 32 NVIDIA V100 workers.

#### Prediction filtering and connectome assembly

After generating predictions from the synapse detection model across the entire h01 sample, a unique identifier was assigned to each pre- and post-synaptic site by running connected components, resulting in 1.9 B unique synaptic sites. While this site count is much larger than the number of expected sites from a sample of this size, we had intentionally biased the network towards capturing more true positive sites at the expense of potential false positives by weighting the positive examples 2x during training. To correct the false positives as a result of the bias, a multi-stage filtering pipeline was applied to ensure site predictions were potentially valid synapses.

First, we discarded synaptic sites that contained less than 30 voxels. We further discarded sites that were in non-neuropil “OOB” regions (as classified by the mask described in “Valid tissue detection”).

Because the synaptic predictions were volumetric in nature and did not take the neurite segmentation into account, it was possible for a synaptic site prediction to span more than one neuropil segment, causing multiple true synaptic sites to be merged into one predicted site after applying connected components. This failure case was particularly relevant to cases where a single pre or postsynaptic partner made synapses with multiple pre or postsynaptic partners, such as a single spine revealing multiple synaptic inputs. To correct for this, we first masked the volumetric labels associated with each site by the segmentation to remove any synaptic labels that exceeded neuropil boundaries, then applied a one voxel 2D erosion to correct for anisotropy. Connected components were then applied to the result. In the case of two or more components, each component was considered a new, unique synaptic site if its volume was greater than 20% of the original prediction’s volume. Any components that did not meet this criteria had their voxels assigned to the closest valid component. Finally, any voxels spanning cell walls that were masked out by the segmentation were joined and assigned to the closest resulting split site.

After splitting, two additional filtering steps were applied. Each post-split site was required to have a minimum voxel size of 100. Next, sites were required to be in areas classified as neuropil, filtering out any sites that were within blood vessels, tears, myelin, or missing data; a site was considered valid if 20% or more of its voxels resided in areas classified as neuropil. Finally each synaptic site was assigned an associated neurite supervoxel based on the neurite segmentation volume (the supervoxel with the largest overlap with the synaptic site mask was selected). If the site mask did not overlap with any supervoxels (potential out of bounds or cell-wall-only), the site was dropped.

After all synaptic sites were identified, split, and filtered, individual sites were paired into synapses. For each site, a local search was performed for any adjacent site within 800 nm. If one or more complimentary sites were identified (e.g. one or more post-sites for a given pre-) and were associated with two unique supervoxels, the closest site was selected as the partner and considered a true synapse. Note that because the process is applied to both pre- and post-sites individually, this pairing process can identify polyadic synapses where one site is partnered to two or more complementary sites.

Because of errors in the original volumetric predictions, the model may correctly identify a synapse, but incorrectly swap pre- and postsynaptic sites. To correct these instances, a final conservative reorientation step was applied using the subcompartment fragment classifications produced for all neurites. For each site associated with a single synapse, the nearest skeleton node was identified based on shortest-path distance through the associated neurite and the node’s class is identified (see section ‘Cellular subcompartment classification and merge error correction’ for fragment categories). If no nearby skeleton node was identified within a radius of 5.3 μm from the synaptic site, the closest skeleton node was assigned based on Euclidean distance (ignoring the segmentation). The identified class was then used to generate a vote to correct (“flip”) the orientation of the synapse. For each site, a set of votes were collected:

1. If an outgoing synapse originates on an axon, the outgoing vote is “no flip”.
2. If an outgoing synapse does not originate on an axon, the outgoing vote is “flip”.
3. If an incoming synapse is not onto an axon, the outgoing vote is “no flip”.
4. If an incoming synapse is onto an axon, the incoming vote is “flip”.
5. If either an outgoing or incoming synapse is onto an astrocyte, no vote occurs.
6. If the nearest skeleton node has no compartment class, no vote occurs.
7. If there is no nearby skeleton node, no vote occurs.
8. If there is no skeleton for the neurite, no vote occurs.

Once all votes were generated for a site, votes were applied based on a consensus measure:

1. If both votes agree to flip or to keep the original orientation, the corresponding correction was applied.
2. If only one vote is present (flip/no flip) the corresponding action is applied.
3. If two votes were generated but they conflict, no correction was applied.
4. If no votes occur, no correction was applied.

#### Correction of synapse over-splitting

Following the processes described above, 190,576,758 individual synapses were identified, although many duplicate synaptic predictions existed at the same individual synaptic site, artificially inflating the number of identified synapses. To correct this, a procedure was applied to consider each pair of synapses sharing the same pre- and postsynaptic associated agglomeration ID and decide whether that pair represented one synapse (should be merged together) or two distinct synapses (should be kept apart), as follows. For each pair, the distance between the centroids of each member of the synaptic pair was calculated. If this distance was less than 750 nm, the pair was merged together. If the distance was greater than 1050 nm, the pair was kept separate. If the distance was in between, then on the pre- and postsynaptic sides of the pair the distance along the skeleton of the pre- or postsynaptic partner between the skeleton nodes closest to each of the two synapses was calculated, and divided by the distance between the two synapses. The greater of these two (pre- and post-) skeleton distances was then used in a logistic regression model (<u>train_synapse_merger_upper_and_lower_thresholds.py</u>) to predict whether the pair should be kept separate or merged together. The overall accuracy of this approach was AUC=0.875. Training data (<u>001_pr_axons.zip</u>) was generated by randomly selecting 117 axons making 10 or more synapses and assessing for each pair of synapses being made onto the same postsynaptic partner, whether they should be merged together or not, using our tool, CREST (see ‘Measurement of synapse prediction errors’, below). After application of the model (<u>get_syn_pairs_with_skel_dists_and_apply_merge_model_parallel.py</u>), the number of predicted synapses fell to 166,216,068. Filtering these synapses to only include those where the presynaptic site was an axon and the postsynaptic site was either a dendrite, soma or axon initial segment, resulted in 133,704,943 synapses.

### Excitatory versus inhibitory synapse classification

To determine whether a synapse was excitatory or inhibitory, we trained a two-class ResNet-50 classifier with an input FoV of [100,100,24], whose input was two-channels: the image data normalized to be centered around zero ((uint8 data - 128) / 33), and a weighted mask of the segments associated with the synapse (pre_weight = 0.95, post_weight = −0.95, background = 0). The center position of each training example was the geometric mean of the pre- and post-masks and included an augmentation that was randomly offset up to +/- [17, 17, 4] voxels in each direction (in order to encourage translation invariance). Additional augmentations of random XY permutations, XY reflections, and 2D rotations around the Z axis were also applied.

Ground truth used to train the classifier was generated by manually proofreading 47 cells (22 excitatory and 22 inhibitory, classified by a single human expert based on morphology) for all outgoing axonal synapses, under the assumption that all outgoing synapses from a single cell were of a common type. Any outgoing synapses that were not originating from axons were discarded, along with any outgoing synapses that were originating from axon fragments containing a merger. This resulted in a total of 2143 excitatory and 2080 inhibitory synapses for training. Since the locations of synapses were determined by the connectome assembly pipeline, no negative examples were generated or used. The model was trained for 733k steps using a batch size of 16 and a learning rate of 2.5·10^-4^.

Given that each dendritic spine is thought to generally correspond to one excitatory synapse ^83^, a subsequent correction step was applied to synapses occurring on spines. We assigned synapses as excitatory if their post-synaptic segment contained a spine-positive skeleton node within 1 μm (see “Spine detection” above), potentially overriding the type determined by the classifier. In addition, any outgoing synapses that originated from a known excitatory or inhibitory cell type - pyramidal and interneuron, respectively - was corrected to the corresponding type.

To assess the performance of the classifier, 2,129 verified synaptic connections identified as arising from excitatory or inhibitory neurons (see section ‘Analysis of manual and ML-generated neuronal networks’, below) were compared to their predicted E or I status (<u>measure_e_vs_i_accuracy.py</u>). Excitatory and inhibitory synapses were correctly classified 99.25% and 89.68% of the time, respectively, with an AUC score of 0.948. Performance was broadly similar across the six cortical layers (**Supp Table 7**). Counts of excitatory and inhibitory synaptic inputs to all neurons were then generated (<u>get_neuron_e_to_i_ratios.py</u>).

### Measurement of synapse prediction errors

To assess our ability to identify synapses arising from axons, where the postsynaptic target structure is either a dendrite, soma or axon initial segment, we made a random selection of 50 synapse-forming axons from across the volume (see <u>50_random_pure_and_majority_axons_FP_and_FN.csv</u>), and verified each predicted synapse. We also searched for synapses that the model failed to predict by examining each bouton or swelling along the axon. The overall false negative rate was 12% and the overall false discovery rate was 1.5%.

### Correction of segmentation errors by computer-assisted proofreading

Since the automatic agglomeration produces neurons and glia that are prone to having split- and merge-errors, we sought a method of correcting these errors efficiently, to facilitate certain analyses that would require 100% correct connectomic data. To this end, we created a simple Python-based tool with a GUI, which makes use of the <u>Neuroglancer Python API</u> and other Python packages to allow the correction of segmentation data, verification and classification of synapses, and structured exploration of the synaptic network of a dataset, called CREST (Connectome Reconstruction and Exploration Simple Tool; <u>CREST_v1.0.py</u>).

For this purpose, a biological object is considered ‘correct’ once all of the base segments that are part of it in the dataset are identified, and no base segments that are not part of it are considered to be part of it. CREST starts with the base segments that have been automatically agglomerated together as part of a single biological object and presents them to the user. The user can then correct split errors by adding on agglomerated segments (which results in all of the base segments that make up that agglomerated segment being added on), or removing base segments that do not belong to that biological object (correcting mergers).

Because CREST treats the single biological object as a graph of connected base segments, correction of mergers is very efficient, as when the user removes a single base segment, all base segments connected to the main part (the part which contains the cell body) via that base segment will be removed at the same time. In the cases where the user wishes to correct structures without cell bodies (such as axonal and dendritic fragments), they may specify which part of that structure is to be considered as the ‘cell body part’ for the purposes of merger correction.

The structures corrected (e.g. neurons) are often highly-branching and complex objects and it can therefore be challenging for the user to keep track of which branches they have corrected. To address this, CREST introduces two features to aid the proofreader. Firstly, the user may specify any number of custom types of Neuroglancer ‘point annotations’, via the GUI, which can be placed at the end of a completed branch, where the ‘type’ of point can be used to record the reason that the branch could not be extended further. The second feature, which builds more on the Neuroglancer Python API, allows the user to mark a base segment and all base segments on the other side of that base segment with respect to the cell body in a certain color. This serves two purposes - firstly to act as a visual aid to the proofreader that a part of a branch is complete (as it is in color - incomplete parts are grey by default). Secondly, the specific color indicates a specific ‘cell structure type’ (e.g. axon, dendrite, cilia), which is saved in a simple JSON state that also includes the complete set of base segments, point annotations and underlying base segment graph, and can be reloaded in CREST for further proofreading.

An example of a complex cell, which contacts 70 postsynaptic partners which also have cell bodies in the volume, completed by CREST in Neuroglancer, is shown in **Supplementary Fig. 15**. Following training in use of CREST and neurobiology, four proofreaders were able to complete 104 randomly-selected neurons (where the center-point of the cell body of each neuron fell between 23 and 30 micrometers in the z axis from the start of the block), which, along with their un-proofread postsynaptic target neurons, form the basis of the network displayed in **Fig. 4I.**

### Analysis of manual and ML-generated neuronal networks

Base segments of 104 proofread neurons whose ‘cell structure type’ had been marked as axonal during the proofreading process with CREST (see ‘Correction of segmentation errors by computer-assisted proofreading’) were identified, and synapses whose presynaptic associated base segments were present within this group were identified as the outgoing synapses of the proofread neurons. These synapses were used to identify the postsynaptic partner neurons with cell bodies in the volume, whether proofread or not (<u>get_edge_list_for_pr_cells_and_unproofread_targets.py</u>), forming the network plotted in **Fig. 4I** (<u>edge_list_to_graph.py</u>).

For the ML-generated network, all connections between segments where both the pre- and post-synaptic segments had been identified as neurons, the presynaptic structure had been identified as an axon and the postsynaptic structure had been identified as either dendrite, soma or axon initial segment, were identified and form the ML-generated network (<u>edge_list_to_graph.py</u>). Because individual connections can erroneously be automatically identified between neurons due to merge errors in the agglomeration or, less commonly, false positive synapse predictions, we sought to measure the false positive rate of each connection type, where one connection type is a combination of a pre and postsynaptic neuron type, and a neuron’s type, for this purpose, is determined by the combination of its cortical layer membership and whether it is excitatory or inhibitory. For example, a layer 2 excitatory neuron forming synapses on a layer 3 inhibitory neuron would be considered one connection type. We made a random selection of connections of each possible connection type (<u>get_subset_of_connections_to_check.py</u>) and created a simple neuroglancer-based python program to manually verify each synapse forming each connection (<u>proofread_connections.py</u>), where an individual synaptic connection between two neurons was considered to be verified if (a) the synapse itself was real and (b) the connection did not arise as a result of merge errors in the agglomeration on either the pre or postsynaptic side. An individual connection (which is defined as all of the synapses formed between two individual neurons), was considered to be verified if at least one of the individual synapses that formed it was verified. Summaries of ML-generated connection type counts and their false positive rates (**Supp. Table 5**), as well as manually-generated connection types (**Supp. Table 6**) were then generated (<u>get_connection_type_accuracy_summary_table.py</u>).

### Identification of cortical layers

For the purposes of analysis, we wished to divide the cortex into layers, each of which would run perpendicularly to the pial-white matter axis. However, we sought a method that would achieve this objectively, without any prior assumptions about the numbers of these layers, or their locations along the pial-white matter axis. We approached this problem in two parts - firstly the identification of clusters of neurons, and secondly, the fitting of boundaries to these clusters.

#### Identification of neuronal clusters

The clustering algorithm HDBSCAN ^84^ was used to identify clusters of neurons based on the distance of the neurons from one another in the x and y axes (we found that including the distance in z made no difference to the result), as well as the ‘distance’ (i.e. difference) in neuronal soma volume. HDBSCAN has two tunable parameters - the minimum number of points that a cluster must possess to be considered as a cluster (*min_cluster_size*), and *min_samples*, which determines how ‘conservative’ the clustering is. To set the value of *min_samples*, we first estimated a probability density function for the neurons, given their positions in x, y and soma size using a Gaussian kernel. For each neuron we then found the mutual reachability distance between it and each neighbor kth- closest or higher, where the mutual reachability distance between between it (a) and one of its neighbors (b) is the maximum of (1) the distance from a to its kth-nearest neighbor (its ‘core distance’), (2) the core distance of b and (3) the distance from a to b. We then calculated the mean mutual reachability distance for each neuron at that value of k, and calculated the correlation between the mean mutual reachability distance and estimated probability density for all neurons. The value of k that produced the greatest correlation between these two measures of neuronal density was selected as the value of *min_samples* for the HDBSCAN algorithm. HDBSCAN was then run, gradually increasing the value of *min_cluster_size* until multiple ‘acceptable’ clusters were identified. Because we sought large clusters running perpendicular to the pial-white matter axis, a cluster was considered ‘acceptable’ if it contained neurons at both the upper and lower edges of the dataset (perpendicular to the pial-white matter axis) and at least 50 neurons in total. This approach identified three clusters. Outlier points were removed from clusters by using a random sample consensus (RANSAC) regressor to fit a second order polynomial through the center of each cluster. For each cluster, the maximum residual threshold such that the resulting curve was still convex was chosen (<u>get_cluster_bounds.py</u>).

#### Fitting boundaries to neuronal clusters

After generating three clusters of neurons, it was possible to fit bounds to the upper and lower edges of each cluster. A concave hull was fit to each cluster, by fitting a Delaunay triangulation to the set of points and then only adding triangles of points to the resulting concave hull if the circumference of said triangle is below a given *ɑ* hyperparameter. Because of the varying density of points in the three clusters, the alpha hyperparameter was tuned to best generate a hull for each cluster of points. Additionally, where there still remained points that extended far out from the central mass of each cluster, by choosing an appropriate *ɑ* value these points were not included in the hull and thus classified as outliers.

With the concave hull of each cluster fitted, the edge points of each concave hull were isolated by only selecting endpoints of edges that were part of one triangle and not two (any edge within the concave hull would be part of two triangles). These edge points were then split into two sets separated by the original central second order polynomial that was fit to each cluster. This method generated two sets of points for each cluster, one corresponding to the upper edge of the cluster and one to the lower edge of the cluster.

Experiments were then conducted to see what curve could best be used to fit the boundary points of the clusters. Arcs of circles were found to be an excellent solution (<u>get_cluster_bounds.py</u>). For each cluster, a circular arc was fit through the center. Let the center of this circle be A. Circular arcs were then fit to the top and bottom edge points of the cluster with fixed center point equal to A. Thus each cluster was bounded by arcs of two concentric circles (see **Supplementary Fig. 16**). These six boundaries (<u>circle cortical layer bounds.json</u>), two for each of the three clusters, divided the cortex into seven regions, corresponding to classical descriptions of layers 1 through 6 and white matter. These boundaries were then used to classify all cells in the dataset into a cortical layer according to what bounds the cell was in between (<u>classify_layers_all_segments.py</u>).

### Measurement of synapse densities

To estimate the distribution of excitatory and inhibitory synapses across XY, the entire dataset was divided into cubes 10 micrometers on each side, and the numbers of each type of synapses counted in each cube. Only 133,440,987 synapses where the presynaptic structure had been identified as an axon and the postsynaptic structure had been identified as either dendrite, soma, or axon initial segment, were considered. The density of E synapses, density of I synapses, and percentage of synapses that are E, was then calculated for each cube, and the values in cubes were then averaged across z (excluding cubes where the total number of synapses within them was zero, to avoid large regions of no data impacting the results) and plotted as heatmaps (<u>plot_synapse_density_and_ei_ratio.py</u>).

### Measurement of layer 5 and 6 triangular cell basal dendrite angles

During manual classification of all neurons, spiny neurons with a distinct bipolar shape, one apical and one prominent basal dendrite, were tagged. Of these, we selected only cells in cortical layer 5 and 6 (estimated radial cortical depth > 1750 micrometers). This yielded a set of 845 neurons for this analysis. For each of these cells, further analysis was done at low resolution (mip 7, every 128th section) in Matlab (The Mathworks, Inc.). We used the manually labeled cell body to extract the Google segment (C3 agglomeration) which overlaps with it maximally (gseg), assuming that that segment corresponds to the analyzed cell. We then computed the bounding box of gseg and used connected components analysis to identify the main connected region of the object while discarding any disconnected parts. Next, we dilated the manual cell body mask by three pixels and subtracted it from gseg to disconnect all branches coming off of the cell body, followed by a second connected components analysis to individualize the branches. Next, we computed center-of-mass coordinates (centroids) of the manual cell body mask and of the two largest branches (by volume), assuming that one of them is the apical and the other the basal dendrite (the axon has a much smaller caliber and would have a much smaller volume that the two big dendrites of these neurons). Centroids which did not end up on the gseg segment were moved to the closest voxel which was part of gseg (minimal Euclidean distance).

Which of the two largest dendrites is the apical dendrite was decided by computing the inner product between the vector from the cell body center to the dendrite centroid and an estimated apical direction. The dendrite running closer to the apical direction was assumed to be the apical dendrite. Cases in which the two dendrites could not be extracted in this automatic way were removed from the list, leaving 816 cells for further analysis.

For the next step we computed a topological field to be able to define ‘radial’ and ‘tangential’ directions at different locations in the tissue. This was done in 2D (XY) only, assuming that the directional variability across sections is minimal. First, we manually fit a grid of tangential and radial lines to 2D projection images of different extracted properties of the sample (cell bodies, excitatory cells, and myelin) by using VAST’s skeleton functions (see **Supplementary Fig. 21A**). We then used Matlab’s function *scatteredInterpolant* to generate interpolated vector fields from these lines (**Supplementary Fig. 21B and C**). These topological fields were also used to define radial and tangential directions in the myelin image in **Fig. 3H**.

To compute the angles at which the basal dendrites run through the tissue with respect to radial, tangential and perpendicular directions, we used the vectors between the cell body centroid and the basal dendrite centroid (basal dendrite vector). Coordinates were scaled to nanometers for correct geometry. We computed polar coordinates of the basal dendrite vector relative to the inverted radial direction at the cell body location (taken from the vector field described above). We name the angle between the inverted radial vector and the basal dendrite vector the radial coordinate ‘rho’ (distance of point from center in Figure 5G), and the angle of the basal dendrite vector around the inverted radial vector the angular coordinate ‘theta’. Basal dendrites with rho below 45° point in a mostly radial direction towards the white matter (dark green dots in Figure 5G). Basal dendrites with rho between 45°-90° and theta between 0°-45°, 135°-225°, or 315°-360° point roughly along the sectioning plane and parallel to the pia (tangential direction; gray points in Figure 5G). Basal dendrites with rho between 45°-90° and theta between 45°-135° point in a mostly perpendicular direction towards section 0 in the stack (light green dots in Figure 5G) and basal dendrites with rho between 45°-90° and theta between 225°-315° point in a mostly perpendicular direction towards section 5292 in the stack (magenta dots in Figure 5G).

For statistical analysis of clustering of cells with basal dendrites pointing towards section 0 and towards section 5292 we selected cells with a basal dendrite pointing towards section 0 as described above (light green, 257 cells) and cells with a basal dendrite pointing towards section 5292 (magenta, 247 cells).

For each of these 504 cells, we found the nearest neighbor and compared their colors, yielding four groups (145 light green-light green, 109 light green-magenta, 112 magenta-light green and 138 magenta-magenta). We performed Fisher’s exact test on these numbers and found that equal-color neighbors were occurring significantly more often than chance (*p* = 0.00745) which we verified by shuffling the colors across cells followed by the same test (*p* = 0.476, n.s.).

### Analysis of axon partners of layer 5 and 6 triangular cell basal dendrites

Axonal inputs onto the largest basal dendrites of layer 5 and 6 triangular cells (see ‘Measurement of layer 6 pyramidal neuron basal dendrite angles’), were identified as follows. First, the skeletons with classified nodes (see ‘Cellular subcompartment classification and merge error correction’) of the layer 5 and 6 triangular cells were converted into graphs. The nodes corresponding to the cell’s soma were automatically identified and removed, creating several disconnected connected components corresponding to the axonal and dendritic branches of the neuron (<u>get_separate_components_of_neurons.py</u>). A point was placed within the thickest basal dendrite of each neuron and the disconnected skeleton branch with the skeleton node closest to this was identified as the thickest basal dendrite branch. The accuracy of this process was assessed by manually measuring, for 20 ‘forward basal dendrite’ and ‘reverse basal dendrite’ neurons, the number of dendritic branches wrongly identified as being part of the largest basal dendrite (90% of both ‘forward’ and ‘reverse’ neurons did not have any wrongly-identified branches), as well as the average number of sub-branches arising from the largest basal dendrite which were not identified as part of it, which was 1.9 for ‘forward’ neurons and 1.3 for ‘reverse’ neurons (Supplementary Table 4). All incoming synapses to that neuron were then downloaded, and those that were located within 3000 nm of at least one of the thickest basal dendrite branch skeleton were identified as synaptic inputs to the thickest basal dendrite branch, and of the the axons making these synapses, those where 100% of their classified skeleton nodes were of the type ‘axon’, were recorded as axonal inputs to the thickest basal dendrite branch (<u>get_partners_for_basal_dendrites_parallel.py</u>).

Having identified the axonal inputs to the thickest basal dendrite (and its subsequent branches), we decided to focus on those axons (n=1180) which made one synapse each on the thickest basal dendrite of two different layer 5/6 triangular cells, both of which would either be in the group of ‘forward-going’ triangular cells or ‘reverse-going’ triangular cells (see ‘New morphological subcategories of layer 6 triangular neurons’). We calculated the proportion of axons that fell into one of three groups; those that targeted two forward basal dendrites, those that targeted two reverse basal dendrites, and those that targeted one of each. For these three proportions we also calculated 95% confidence intervals,^85^ shown as blue bars in **Fig. 5G**. We observed that ‘forward’ basal dendrites receive more synapses than reverse basal dendrites, and have more skeleton nodes (**Supplementary Fig. 20**), suggesting that axons have a greater probability of randomly forming synapses onto ‘forward’ basal dendrites than ‘reverse’. Therefore, to take this into account, we decided to calculate expected proportions (or FF, RR and FR) under a null model whereby each axon selects two basal dendrite partners randomly, but with the probability of choosing either a forward or reverse basal dendrite determined by the total number of synapses being made onto these two types of basal dendrite, resulting in the point estimates shown as red dots in **Fig. 5G**. From these estimated proportions, we were able to calculate expected frequencies under the null model, and these were compared to the observed frequencies by a chi-squared test, which resulted in a p-value of 3.34·10^-14^, with the same approach used for excitatory (n=808, p = 1.60·10^-7^) and inhibitory (n=372, p = 9.89·10^-6^) subsets of these axons (**Supplementary Fig. 24**). This analysis excluded those axons making a synapse on any axon initial segment (AIS), whose interactions with layer 5/6 triangular cells were analysed separately in exactly the same way, except that axons targeting more than two triangular cells were included, and two triangular cell partners randomly selected from each axon’s set of postsynaptic targets (n=174, p = 0.319; <u>basal_dendrite_analysis.py</u>).

### Analysis of axonal selectivity

To simulate the expected frequencies of different connection strengths in the dataset for comparison to the observed frequencies (**Fig. 6**) we took the four-part approach described below; firstly, identifying for each synapse the point at which it attached to the shaft of its parent axon, secondly, measuring the distances between either ‘en passant’ or ‘terminal bouton’ synapses to the axonal shaft and estimating the distributions of these distances, thirdly, using these estimated distributions to simulate the random formation of synapses arising from the axonal shaft, and finally, using these simulated random synapses to estimate the expected frequencies of different connection strengths across the dataset as a whole.

#### Identification of distinct axonal skeleton components

To facilitate the analysis of the connectivity of individual axons, methods were developed to identify separate components of the skeletons of axons, in particular distinguishing the ‘shaft’ of the axon from its terminal bouton stalks, and based on this, distinguishing synapses that are intrinsic to the shaft (en passant synapses) from those that arise from the shaft on terminal bouton stalks (see **Supplementary Fig. 17** for a representative example). To achieve this we first attempted to identify the skeleton node where each synapse connected to the shaft (hereafter ‘root node’), whether or not that synapse was on a stalk or an en passant synapse. To do this, we first used CREST to manually mark the root nodes of several synapses. We then trained a simple model which for each synapse, (1) identifies the longest path from the skeleton node closest to the synapse to any ‘end node’ in the skeleton, (2) considers each node on this longest path in order, starting from the synapse-associated node, and decides whether it is the ‘root node’ of that synapse, based on whether the two longest paths arising from that candidate root node’s immediate neighbour nodes to end nodes in the skeleton (and not including current or previously-considered root node candidates, or the synapse-associated node) each exceed specified minimum lengths. Once a root node candidate is identified which meets these criteria, the search is terminated and that candidate is designated the root node of that synapse. If a specified number of candidate root nodes are rejected, then no root node is assigned to that synapse at that stage. All paths between ‘end’ nodes which pass through the root nodes, but not through synapse nodes or nodes on the paths between synapse nodes and their individual root nodes, are identified, and designated as ‘shaft’ nodes. For synapses without identified root nodes at this stage, their root node is set to the closest shaft node. Any branches shorter than a certain specified length that do not contain synapses are removed. The model therefore has four tunable parameters; the number of candidate root nodes that will be considered for each synapse, the lengths that the longest and second-longest paths arising from each candidate root note much exceed to be accepted as a root node, and the minimum length for a non-synapse containing side-branch to be kept ( <u>train_skeleton_pruner.py</u>, <u>prune_a_batch_of_skeletons_ig_parallel.py</u>). This process identifies root nodes with an accuracy of 94.4% and distinguishes shaft nodes from non-shaft nodes with an accuracy of 94.3%. Once this process is complete, the distance from each synapse-associated node to its root node is used in a simple logistic regression model to classify that synapse as either an ‘en-passant’ synapse or a ‘terminal bouton’ synapse (AUC = 0.968).

#### Estimation of distributions of distances from synapse to axonal shaft

For many connectomic analyses, it is desirable to compare observed results to the range of results that might have occurred given some null model. One approach to this for analyses concerning the targeting preferences of axons, is to simulate which postsynaptic partners any given axon might have been expected to target if it were making its outgoing synapses according to a process that is in some sense ‘random’. However, for any such random simulation / null model to be biologically plausible, it must be constrained in some way by real features observed in the data. To create a biologically plausible model under which an axon might make random outgoing synapses, we first examined the range of distances between synapses and the axonal shafts from which they arose, plotting the range of differences for ‘en passant’ synapses and ‘terminal bouton’ synapses separately. We observed that ‘en passant’ and ‘terminal bouton’ synapses are on average made 270 nm and 1500 nm from the axonal shaft respectively, with both distributions having long-right-hand tails. We were able to fit probability density functions (PDFs) to each of these datasets (see **Supplementary Fig.18** and <u>fit_distribution_to_synapse_distances.py</u>).

#### Simulation of random connectivity of an individual axon

We then devised a method of simulating random connectivity for a single axon as follows. First, the axon is moved (translated) by a user-specified distance in the x, y and z axes, or not at all. For each ‘en passant’ synapse, a random point along the shaft of the axon is chosen (the ‘root point’ of this simulated synapse), a distance for the simulated synapse to occur from the randomly-select root point is chosen according to the PDF of distances described above for ‘en-passant’ synapses, and then a random angle is chosen to determine the direction from the shaft that the simulated synapse will be made from, constrained to a plane that is perpendicular to the shaft skeleton at that root point. For each terminal bouton stalk synapse, the same process is used to generate a simulated synapse location around the shaft, except that the PDF of distances for terminal bouton stalk synapses is used instead, and the simulated synapse location may project from the root point in any direction in 3D space around it. Once a simulated synapse location has been selected, it the segment ID at that point lies on the surface of a segment which is included in a user-supplied list of ‘acceptable post-synaptic partners’ (typically dendrites), then it is accepted as a simulated synaptic connection. This process is repeated to simulate that axon’s connectivity an arbitrary number of times that is specified by the user. An alternative, more ‘constrained’ version (as opposed to the ‘unconstrained’ version described above) of this method of simulation has also been devised, whereby the simulated synapses arise from the actual root points of their real counterparts (<u>sample_points_around_neurites.py</u>). We verified for both ‘constrained’ and ‘unconstrained versions for a random subset of axons that the distribution of the sampled points for each individual axon followed the targeted distribution of synapse distances shown in **Supplementary Fig.18**. The distribution of distances for the simulated synapses was a close match to the empirically measured one.

#### Analysis of axonal selectivity

CREST was used to identify multisynaptic connections between neurons (i.e. strong connections). To establish whether the numbers of strong connections we observed in the data were likely have arisen by random chance, we first randomly selected 10,000 axons of each ‘strongest connection strength type’, where, for example, an axon whose strongest post-synaptic connection was formed of three synapses would have a ‘strongest connection strength type’ of 3. Where there were less than 10,000 axons in a certain connection strength type, we selected all axons of that type. Furthermore, we only selected axons which did not make any synapses on axon initial segments of neurons, and which only have ‘axon’ classified nodes in their skeletons (hereafter ‘pure axons’, see <u>get_sample_of_axodendritic_only_axons_organised_by_strength.py</u>). This resulted in a selection of 53,695 axons. We then used the ‘unconstrained’ version of the null model described above with a random 15 micron displacement in the x and y axes (‘Simulation of random connectivity of an individual axon’), for each axon simulating the same number of ‘en passant’ and ‘terminal bouton stalk’ synapses as are found on the axon, to simulate the postsynaptic targets of that axon, to estimate its strongest connection strength under the null model. To prevent axons being moved to within the somas of other cells, we rejected any randomly-selected 15 micron XY displacement location that fell within 25 microns of the centre of a cell body, resulting in 9,190 of the 53,695 axons having their partners randomly simulated. Following the simulation, we manually checked all axons making a high number (>=6) of simulated connections onto a single postsynaptic partner for any artifactual causes of such strong connections, and excluded one axon from the subsequent analysis. Each group of simulated axons (where one group is all axons with a certain combination of inhibitory/excitatory and a certain strongest partner strength) could then contribute simulated outcomes proportional to the overall occurrence of that group of axons in the dataset as a whole, allowing us to calculate expected frequencies of each type for all ‘pure axons’ and compare these to the observed frequencies with the chi-squared test (<u>connection_strengths_analysis.py</u>).

### Analysis of information flow through network

An <u>animation of information flow</u> in the automatically-generated network was generated in Python by stepping through discrete time points based on the following preset rules: 1) only pyramidal cells and interneurons with somata inside the volume were considered; 2) when a pyramidal neuron was activated, it added +1 per synapse to the inputs of all its postsynaptic partners; 3) when an interneuron was activated, it added −1 per synapse to the inputs of all its postsynaptic partners; 4) when the summation of all the input synapses to a neuron was larger than 0, it would be activated at the next time step and last for one time step. At the beginning of the process, all the pyramidal cells in cortical layer 4 were set to active. The activated neurons are shown as bright green (pyramidal) or bright red (interneurons) circles. Neurons receiving negative net inputs at the current time step are colored blue. Each time step is subdivided into multiple frames for visualization purposes, with the neurons receiving the largest absolute net inputs changing their color first followed by the ones receiving smaller net positive or negative inputs. The output edges of all the activated neurons were marked as colored straight lines (green for pyramidal and red for interneurons).

---

**Supplementary Figure 1:**
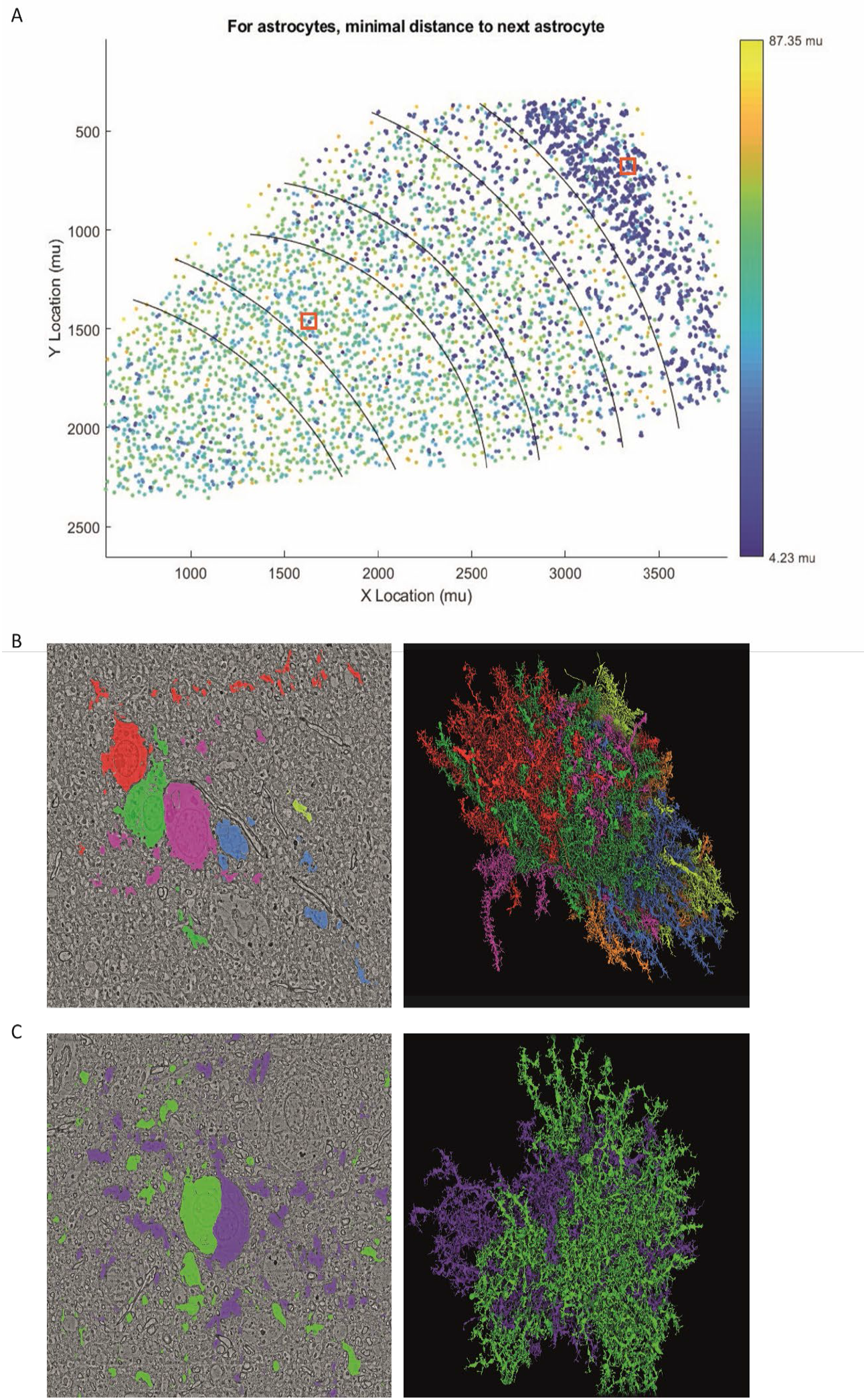
Minimal distance between astrocytes in cortical layers. **(A)** The Y and X coordinates for all 5474 astrocytes in the volume colored by the minimal distance to the next astrocyte. Red marking shows the approximate location of examples highlighted in B and C. **(B)** Example of six astrocytes in layer 1 (left panel shows four of the six cell bodies in the same EM section) resulting in cell aggregate with intermingling arbors, as shown in the 3D rendering (<u>link</u>). **(C)** Example of two astrocytes in layer 5 with closely connected cell bodies and overlapping territories (<u>link</u>).

**Supplementary Figure 2:**
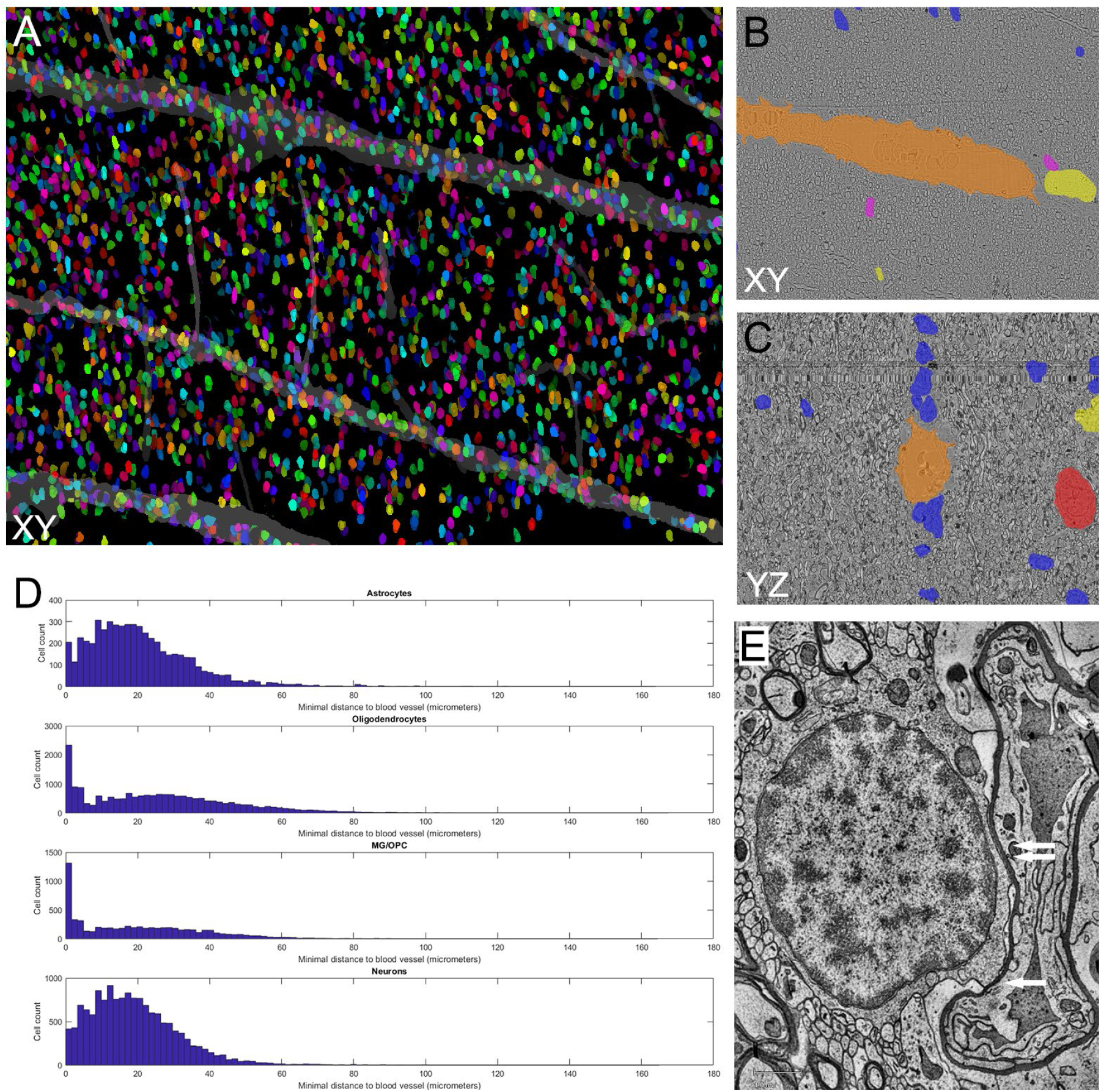
Affinity of oligodendrocytes for blood vessels. **(A)** Oligodendrocyte cell bodies (randomly colored) cluster at large blood vessels in white matter. In turn, around the blood vessel there is a region of lower oligodendrocyte density. EM image cross-sections in XY **(B)** and YZ directions **(C)** reveal that oligodendrocytes (blue) cluster above and below the large blood vessel (orange), but not at the sides. This suggests, at least in white matter, a mechanical reason for their location rather than a metabolic one (possibly related to tissue anisotropy; most myelinated axons in the area run along the Z axis). **(D)** Minimal distance between different cell types and blood vessels is shown in the x-axis (micrometers) and number of cells in the y-axis of the histogram. Astrocytes and neurons show similar distance to the blood vessels, while a large portion of oligodendrocytes and microglia/OPC’s are within a 5mu distance to blood vessels. **(E)** Cross-section of an example showing a perivascular oligodendrocyte. Single arrow shows a location where the oligodendrocyte membrane touches the basement membrane, double arrow shows a close-by location where the oligodendrocyte is separated from the basement membrane by a thin layer of astrocytic endfeet.

**Supplementary Figure 3:**
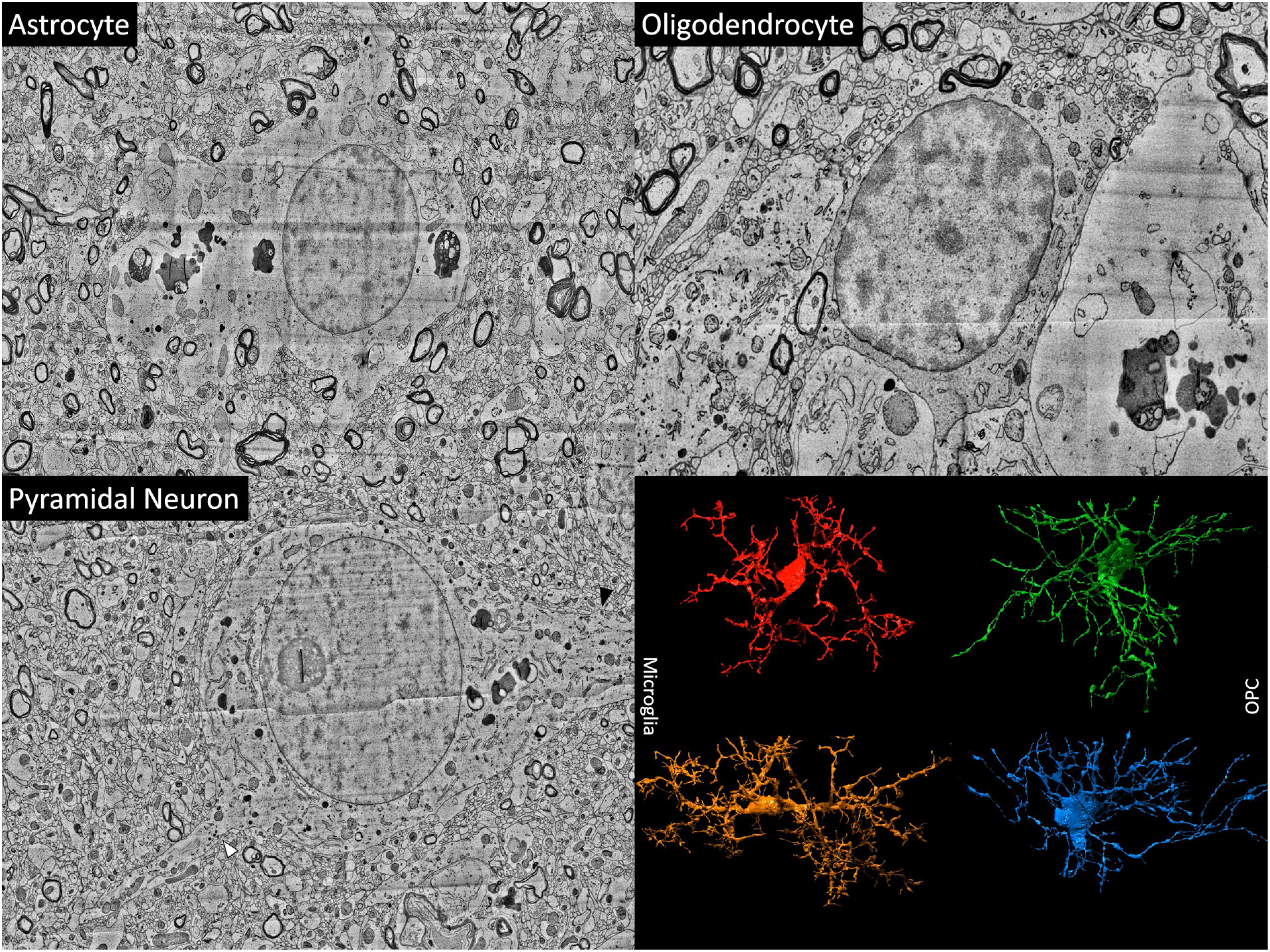
Cell type examples from our dataset. Example cross-sections of an astrocyte with empty-looking cytoplasm and typical feathered outline; an oligodendrocyte with gray cytoplasm, and a pyramidal neuron with visible apical dendrite (black arrow) and axon initial segment (white arrow). Lower right shows two examples each of microglia and oligodendrocyte precursor cells (OPCs) as 3D-rendered models.

**Supplementary Figure 4:**
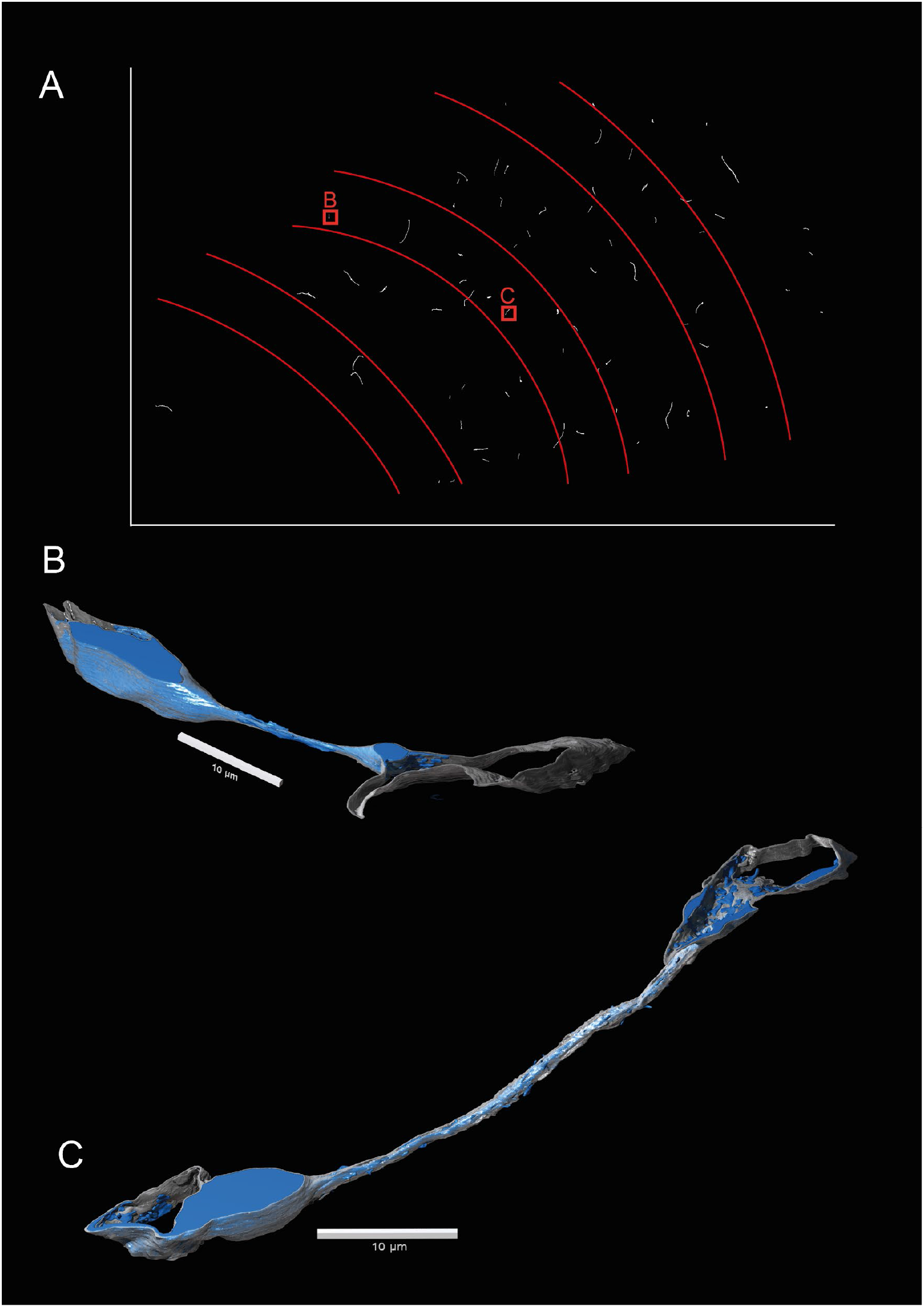
Thin bridges between blood vessels. **(A)** Among the segmented blood vessels we identified 73 thin bridges without blood vessel lumen, consisting solely of basement membrane and pericyte soma. Here only the thin bridges are shown, seen in red in **Fig. 3G**. These bridges were rare in white matter and layer 6. Two examples were investigated in detail, location shown in square boxes. **(B)** The thin bridge is approximately 10 μm and connects two closely located capillaries (<u>link</u>). Pericyte cell body is colored blue and the basement membrane is gray. The pericyte nucleus and large portion of soma is located in one of the capillaries. The pericyte projects through the bridge, which is only partly covered by basement membrane, and into the neighboring capillary interacting with the endothelial cell. **(C)** Shows a thin bridge approximately 40 μm long between two capillaries (<u>link</u>). The pericyte projection in the thin bridge is mostly covered by the basement membrane, however several pericyte protrusions and gaps in the basement membrane were observed.

**Supplementary Figure 5:**
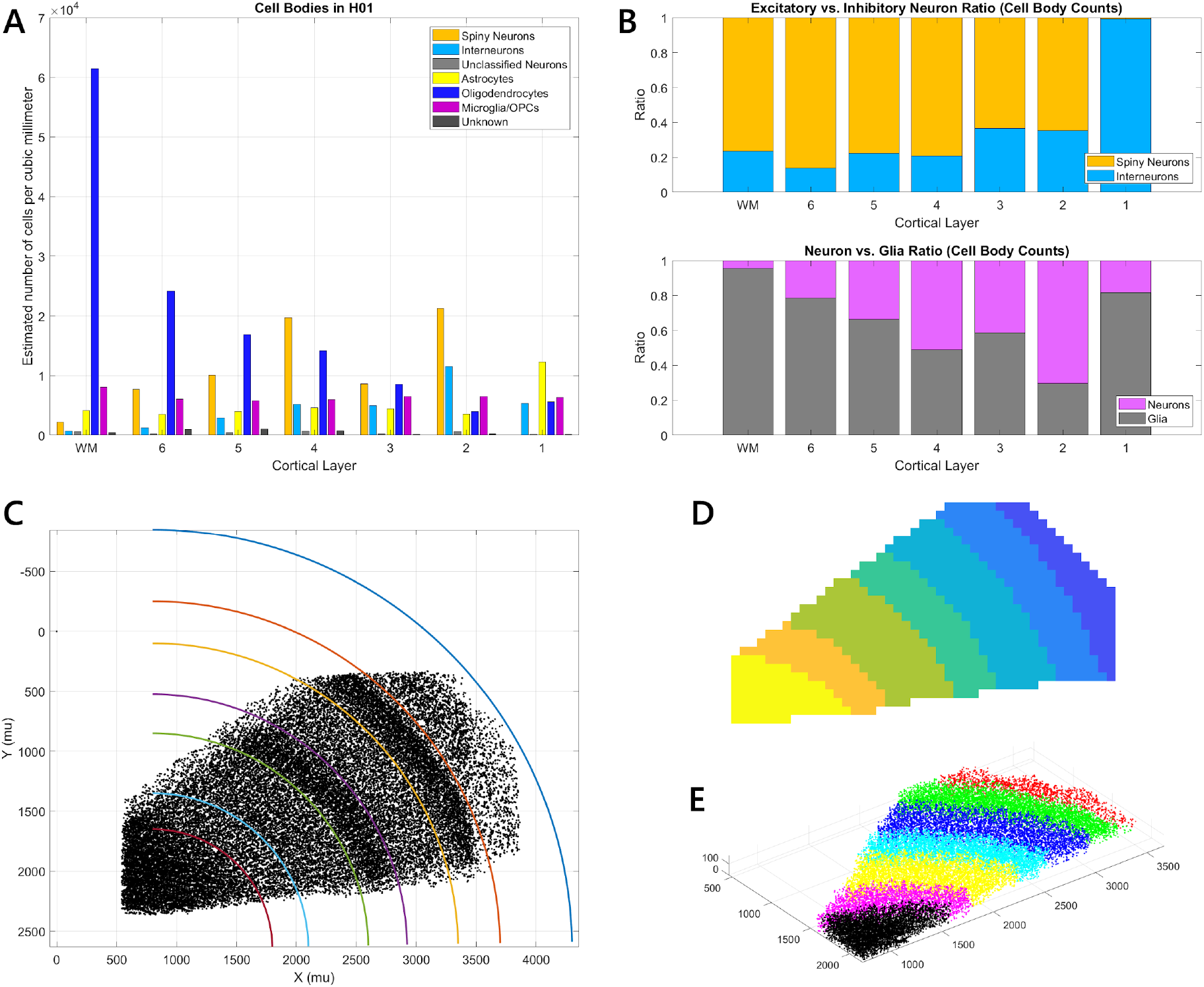
Cell densities per type and layer. **(A)** shows densities of cell bodies of different types per cubic millimeter in our sample, separate for each layer. These values are corrected for the estimated 28% mechanical compression in the cutting direction. The numbers for this plot are listed in **Supplementary Table 8**. **(B)** Shows the relative abundance of excitatory versus inhibitory neuron cell bodies (top) and neurons versus glia (bottom), for the layers in A. **(C)** For this analysis, layers were approximated by using a circular distance measure from a center point at (800, 2650) which approximates the curvature in the tissue. **(D)** To get cell body densities the number of cell bodies of each type has to be divided by the volume of tissue they are found in. Only the volume in which cell body labeling was fully saturated was used (see **Supplementary Fig. 10**), and individual tiles were associated with one layer each. **(E)** shows the XYZ locations of the cell bodies included in this analysis and their layer assignment in color. Note that towards layer 1 the upper layers are cut in deeper sections; this was taken into account for the volume estimation.

**Supplementary Figure 6:**
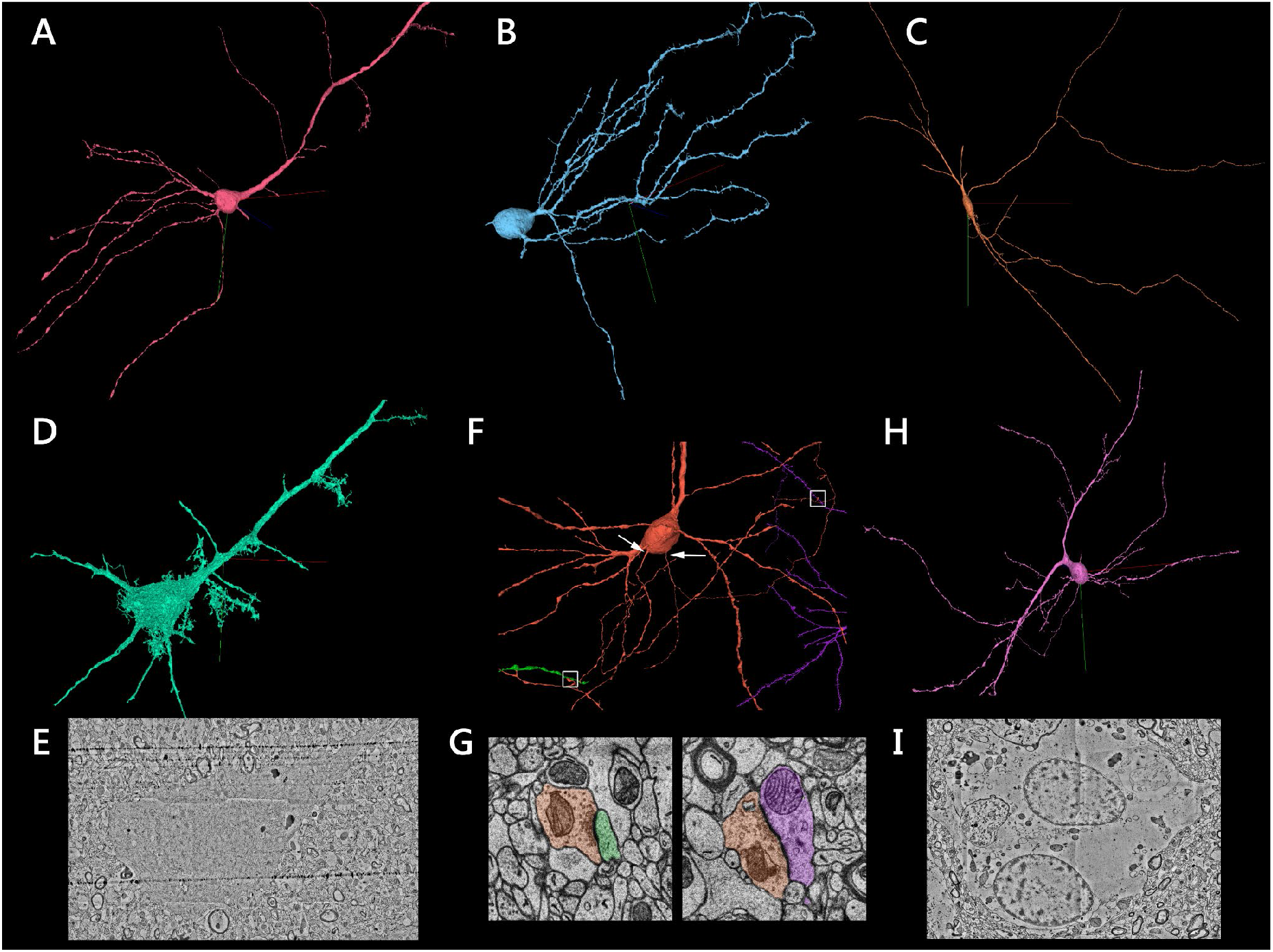
Examples of unusual cells. **(A)** Shows a <u>neuron with clear pyramidal shape which is almost devoid of spines</u>. **(B)** Shows a <u>neuron with interneuron morphology but many spine-like protrusions on its dendrites</u>. **(C)** Shows a <u>rare interneuron which extends horizontally</u> (tangentially) through our data set. This cell can be seen *in situ* in **Fig. 3C** in the middle of layer 3. **(D)** and **(E)** Show an example of a <u>‘dark’ cell</u> in the tissue which has pyramidal cell morphology. Most dark cells were found towards the upper edge of our sample, in layers 4-6. **(F)** This <u>neuron has two separate axons emerging from the soma</u> (white arrows). Both make outgoing synapses in the volume (white boxes), shown below in **(G)**. **(H)** Shows a <u>neuron with an unusual dendritic tree</u> (cell body displaced to the side). **(I)** Shows a cross-section of an <u>astrocyte cell body with two nuclei</u> (separate in 3D). We found 34 cells with two clearly separate nuclei in the tissue, most of which were astrocytes.

**Supplementary Figure 7:**
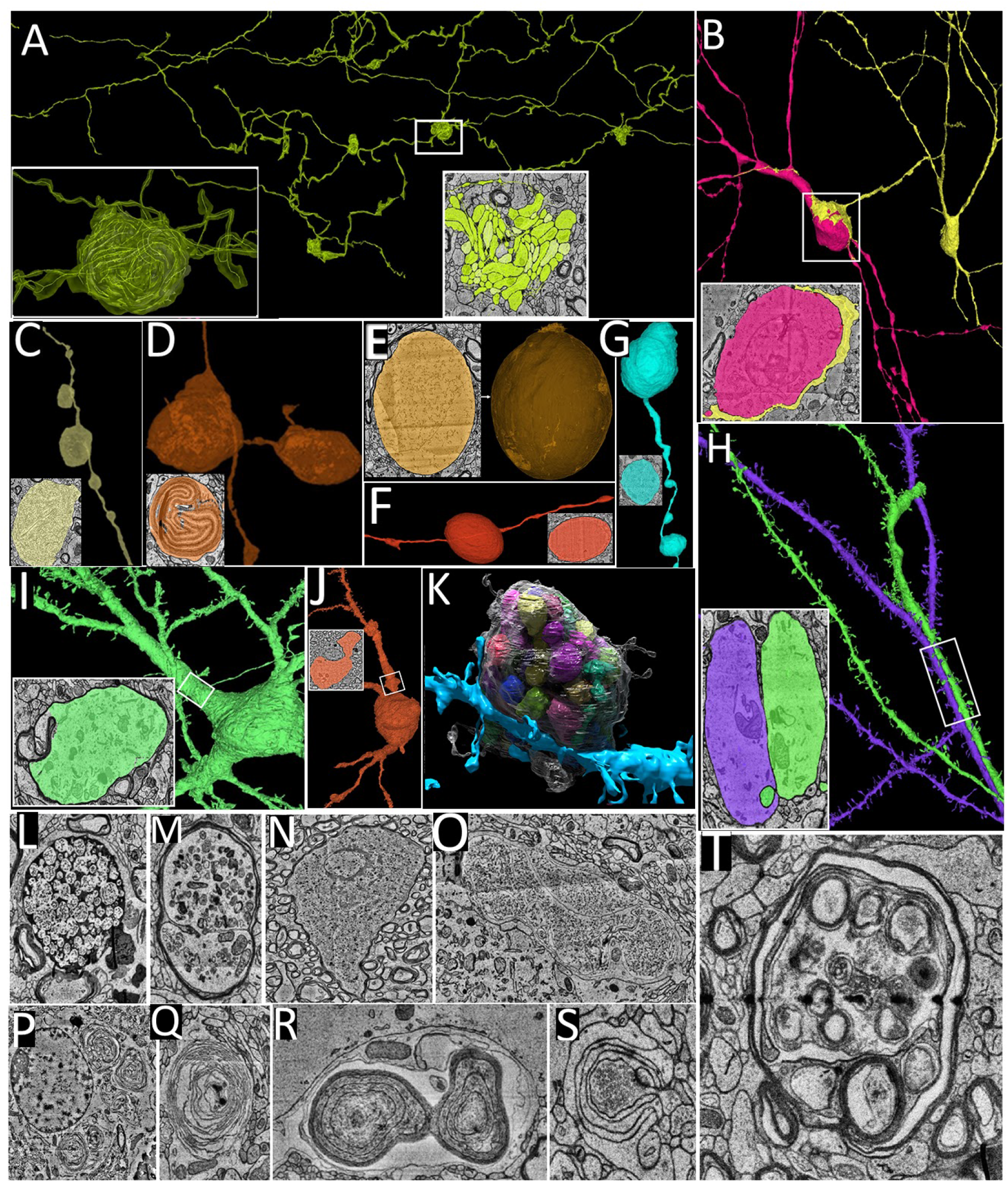
Unusual and unidentified cortical objects. **(A)** Whorled axon that makes inhibitory synapses on cell soma and dendritic shafts in L5, L6 (<u>link</u>)**. (B)** Soma of interneuron engulfed by the dendritic process of another interneuron (<u>link</u>)**. (C)** Unidentified object filled with fibrous substance (<u>link</u>**). (D)** Whorls of loosely coiled myelin (<u>link</u>). **(E)** Egg shaped object with no associated processes (<u>link</u>). **(F)** Myelinated oval object. **(G)** Greyish object. **(H)** Interacting climbing dendrites <u>link</u> **(I)** Myelinated dendritic trunk in grey matter (<u>link</u>). **(J)** Myelinated dendritic trunk in white matter (<u>link</u>). **(K)** Swollen dendritic spine containing 150 intramembranous objects (segmented manually; <u>link</u>). **(L)** Unidentified object filled with small spherical objects (<u>link</u>). **(M)** Myelinated axon filled with unidentified substance (<u>link</u>). **(N)** Myelinated object filled with small debris (<u>link</u>). **(O)** Compartmentalized fibrous object (<u>link</u>). **(P)** Whorls within the soma surrounding a cell nucleus (<u>link</u>). **(Q)** Random lying whorled substance (<u>link</u>). **(R)** Conjoint whorls (<u>link</u>). **(S)** Synapse wrapped by astrocytic processes (<u>link</u>). **(T)** Myelinated object with multiple membranous rings (<u>link</u>).

**Supplementary Figure 8:**
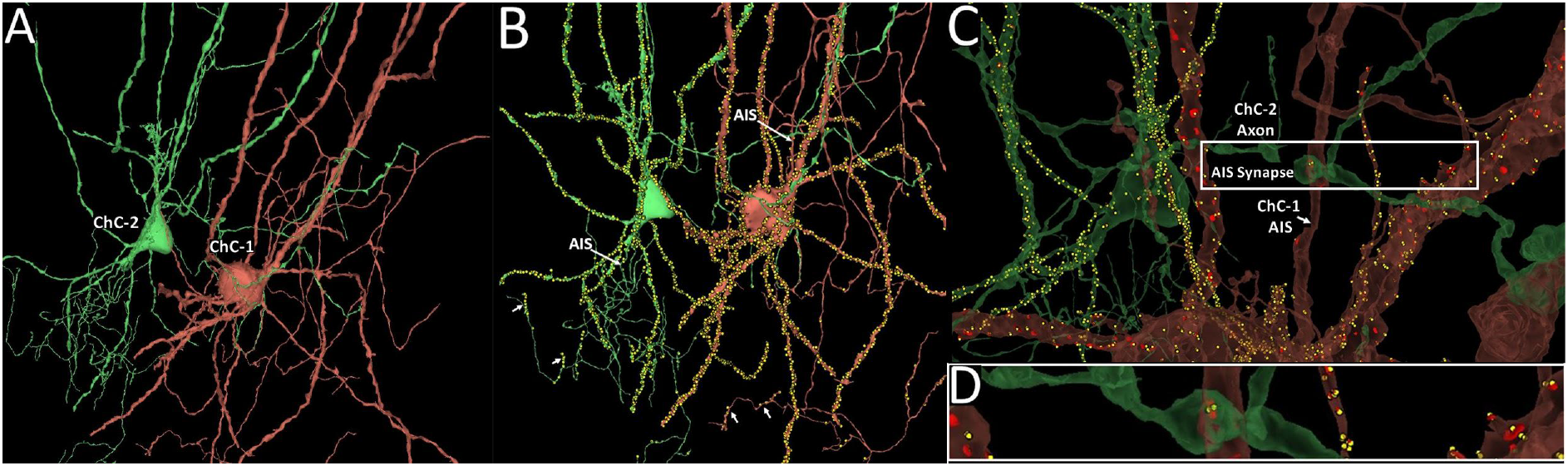
Chandelier cells. **(A)** Two chandelier cells, ChC-1 and ChC-2. **(B)** Incoming and outgoing synapses on both ChC-1 and ChC-2 chandelier cells. The arrows indicate the characteristic cartridges made by chandelier cell axons. **(C)** Incoming inhibitory synapse on AIS of ChC- 1 made by the axon of ChC-2 (<u>link</u>). **(D)** Close-up of the ChC-ChC AIS synapse shown in panel **C**.

**Supplementary Figure 9:**
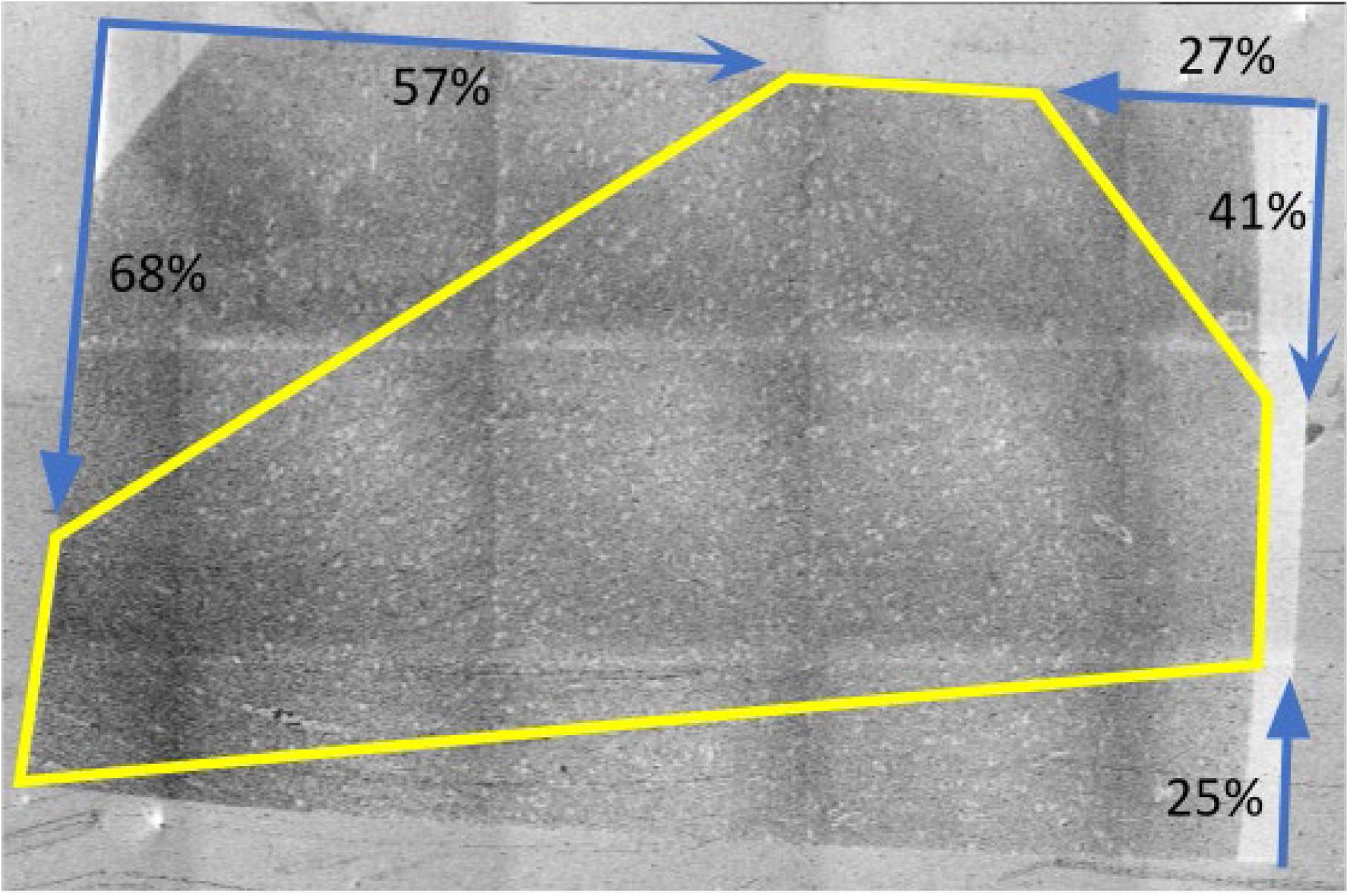
Low-resolution electron microscopy overview of a representative section. Region of interest (ROI) is indicated by a yellow polygon. Blue arrows and corresponding percentages indicate the minimum distance (expressed as a proportion of that edge of the section, including any resin) between the corner of the section at the foot of the arrow, and the point of the ROI at its head.

**Supplementary Figure 10:**
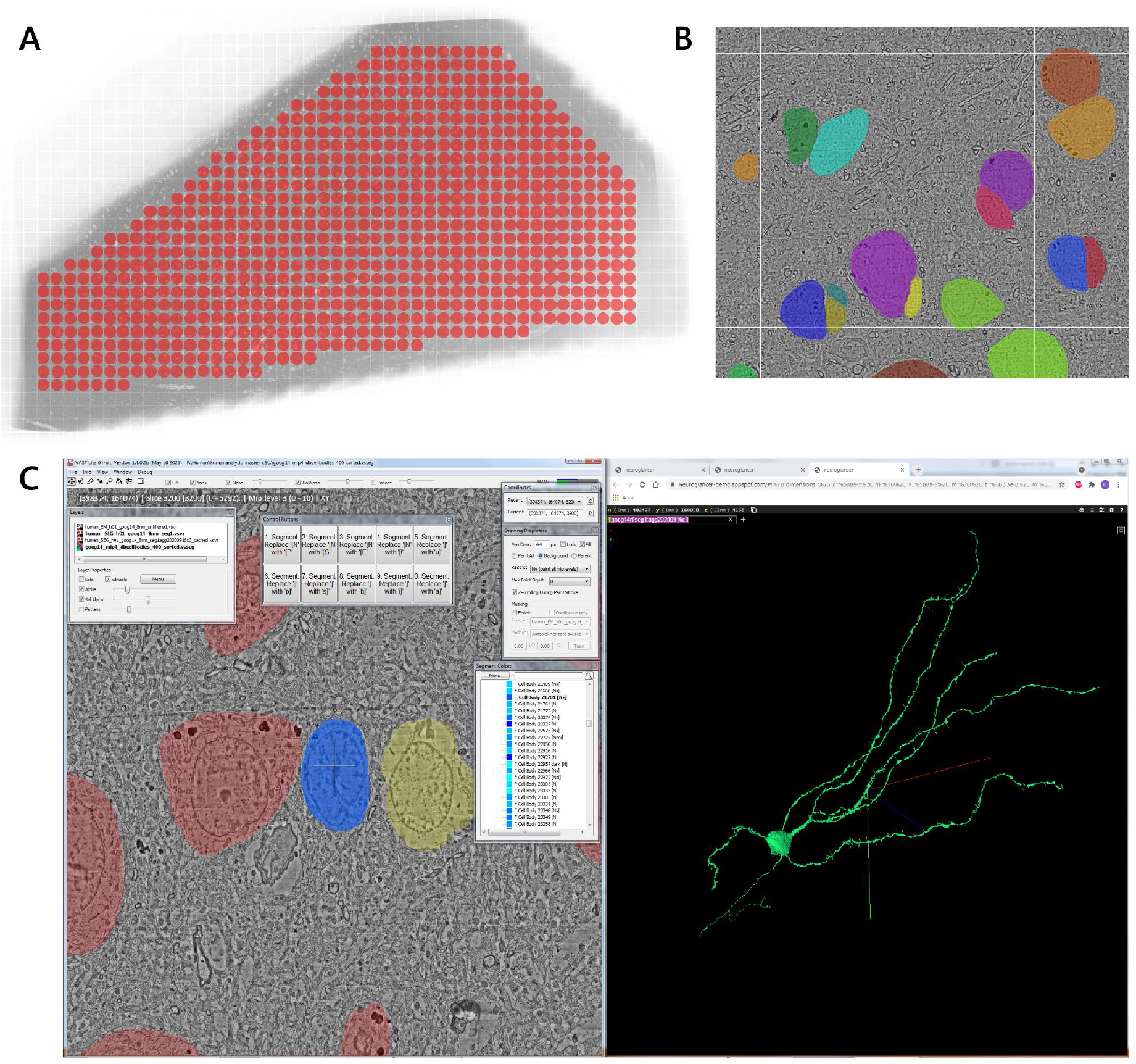
Manual cell body labeling and classification. **(A)** Systematic search for all cell bodies in the tissue was done in VAST by using an overlaid grid (image layer). Each block which was surveyed was marked by a red dot (separate segmentation layer). The area of red dots shows where very close to 100% of all cell bodies should be labeled. **(B)** Shows the resolution at which manual cell body labeling in VAST was done (mip 4, every 128th section) and the approximate accuracy of manual labels. **(C)** Cell body classification. VAST (left) is linked to Matlab via its API and vasttools.m. A Matlab script is running in the background which polls for SPACE key presses in VAST. Once the user presses the SPACE key, Matlab remote-controls VAST to advance to the next segment in the list and moves the 2D view in VAST to that location. At the same time it queries the corresponding ID in the automated segmentation and opens a Neuroglancer 3D view of the same cell in a browser window. The user can then inspect the cell in 2D and 3D and make a decision for classification. The classification key is added to the name of the cell segment in the VAST ‘Segment Colors’ list as a list of tags in square brackets. Specific tags can be added by clicking the buttons in VASTs configurable ‘Control Buttons’ window. The names list can later be exported together with other metadata and used for analysis. (<u>See an interactive view of all annotated cells</u>.)

**Supplementary Figure 11:**
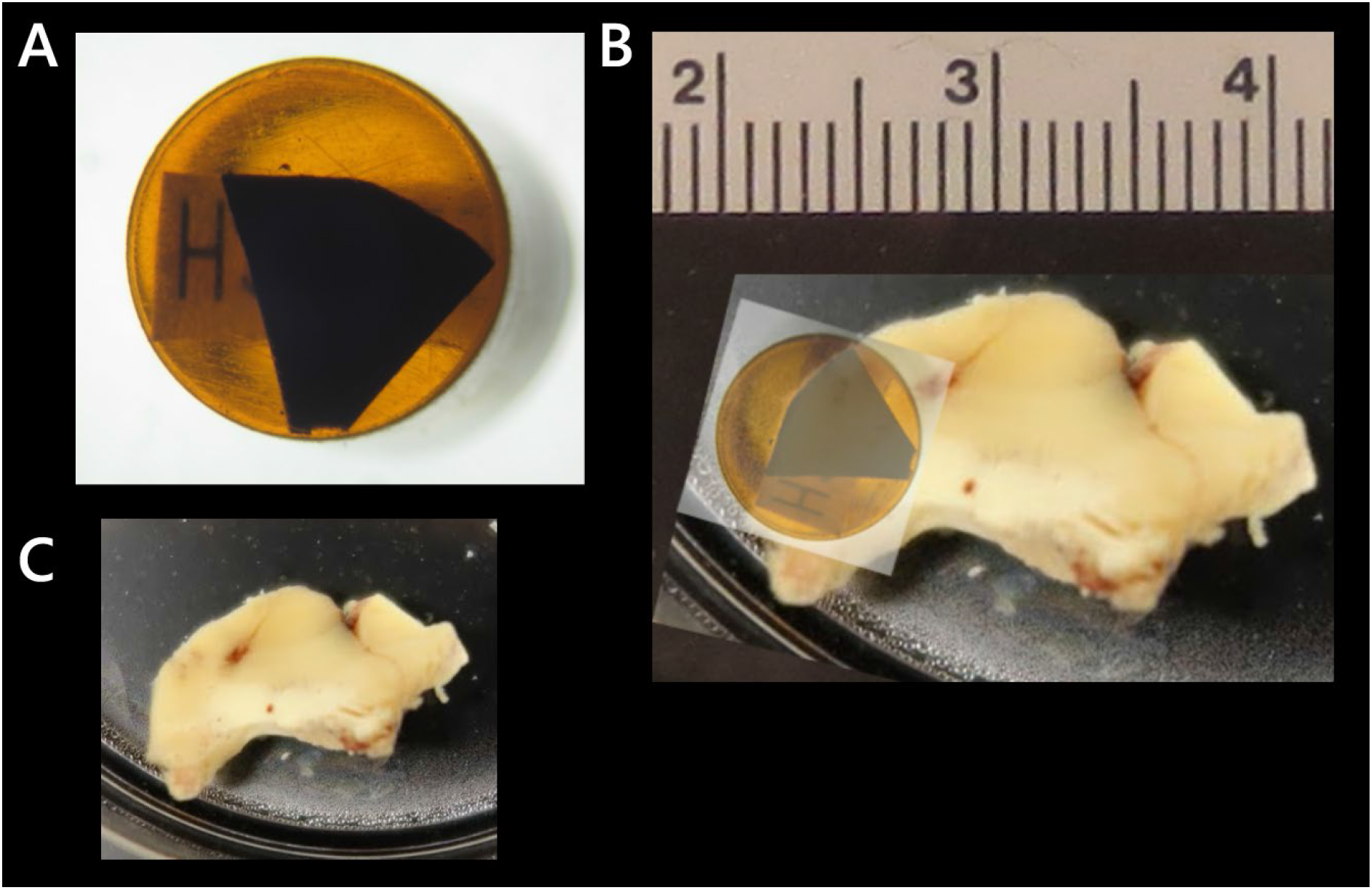
Tissue sample. (**A**) The human brain vibratome section in its resin block, stained, embedded and ready for trimming and sectioning. This vibratome section was taken from a larger brain biopsy sample of the human temporal cortex (**B**) which was approximately 2 x 1 cm in size. **(C)** The fixed sample before vibratoming.

**Supplementary Figure 12.**
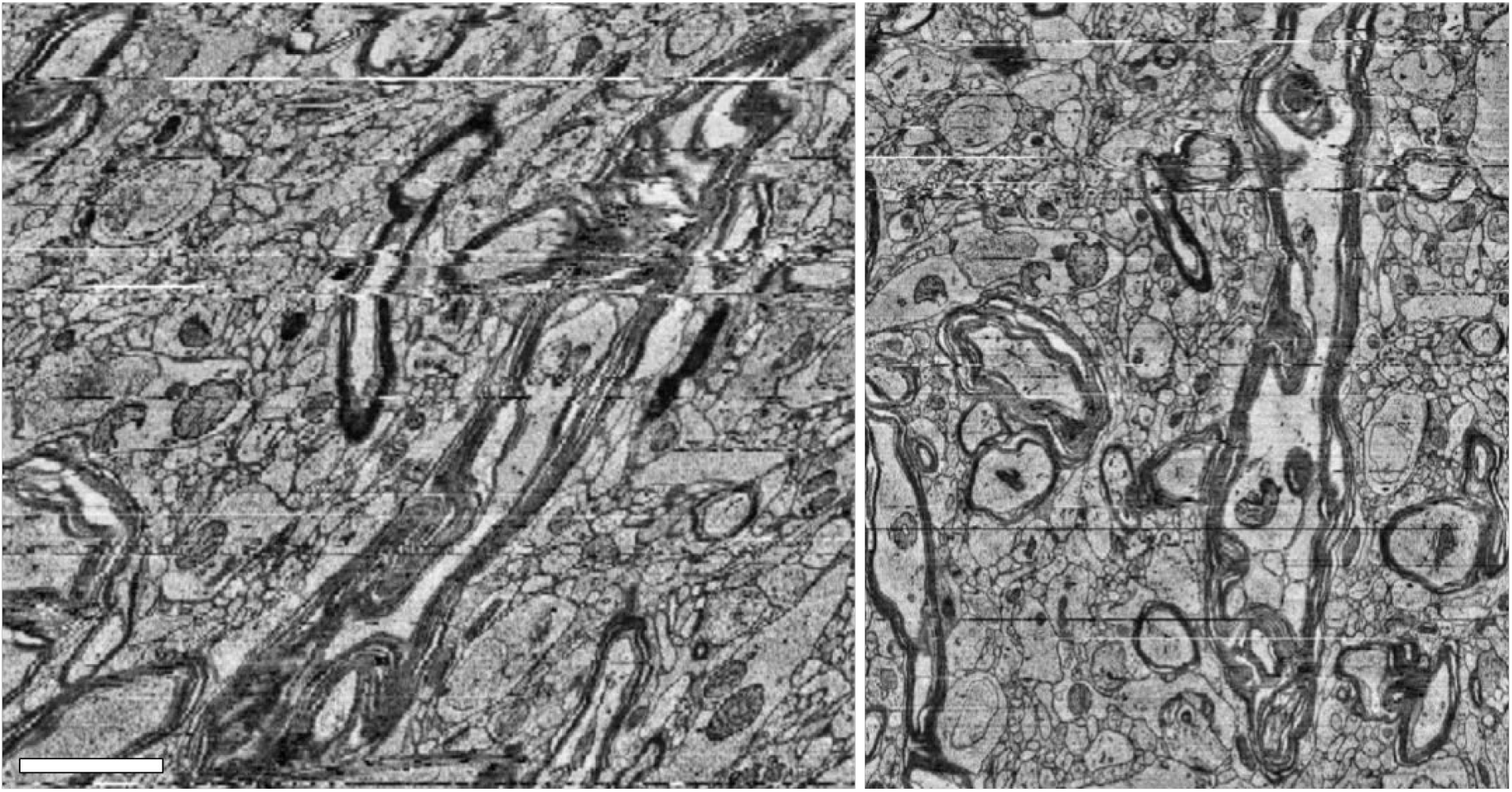
Optical flow field correction. XZ section of a part of the dataset exhibiting drift and some misalignments before (left) and after optical flow field correction (right). Scale bar 2 μm.

**Supplementary Figure 13.**
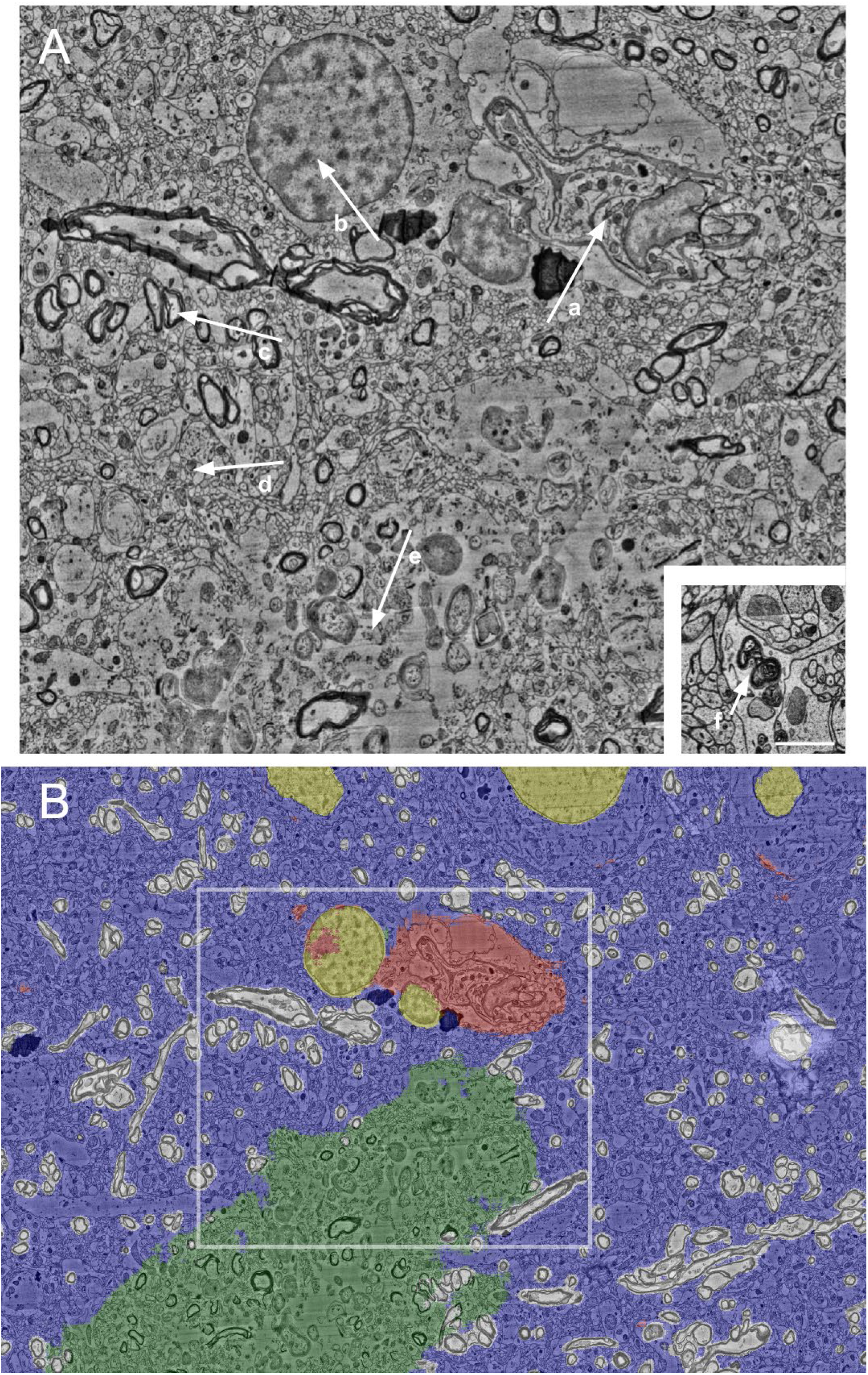
Semantic segmentation predictions. **A:** tissue types detected by the semantic segmentation model. (a) Blood vessel (b) Nucleus (c) Myelin (d) Neuropil (e) Fissure (f) Myeloid bodies within a dendritic branch. Scale bar is 1 μm. **B:** predicted masks indicating cell nuclei (yellow), blood vessels (red), tissue fissure artefact (green), myelin (light gray), other neuropil (dark blue). White frame indicates the region shown in A.

**Supplementary Figure 14.**
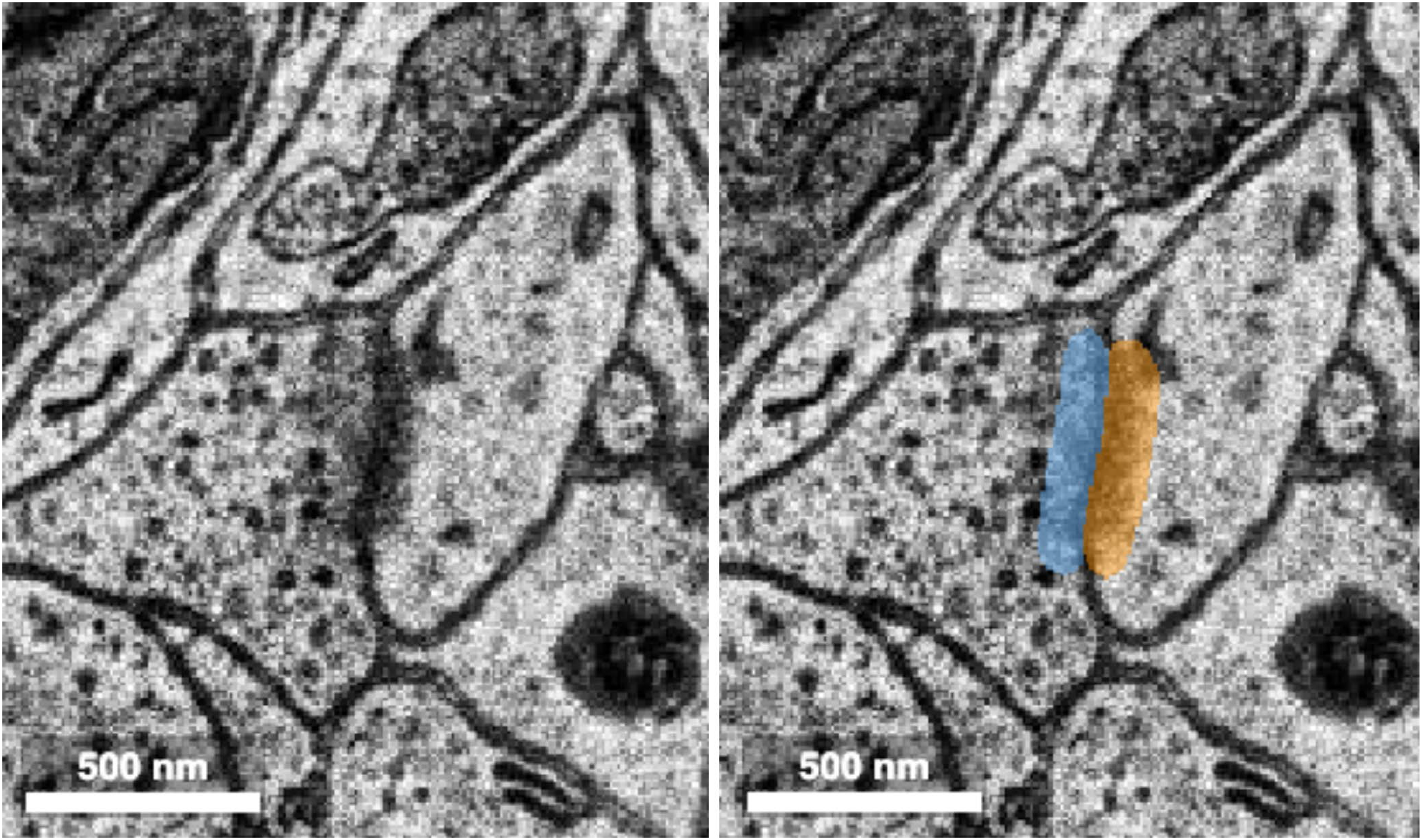
Synapse prediction. Left: Typical example of a chemical synapse, showing vesicles in the axon, dark post-synaptic density at the cleft, and a lack of vesicles in the receiving dendrite. Right: voxel-wise annotations denoting presynaptic site (blue) and postsynaptic site (orange). Other voxels are assumed to be background.

**Supplementary Figure 15.**
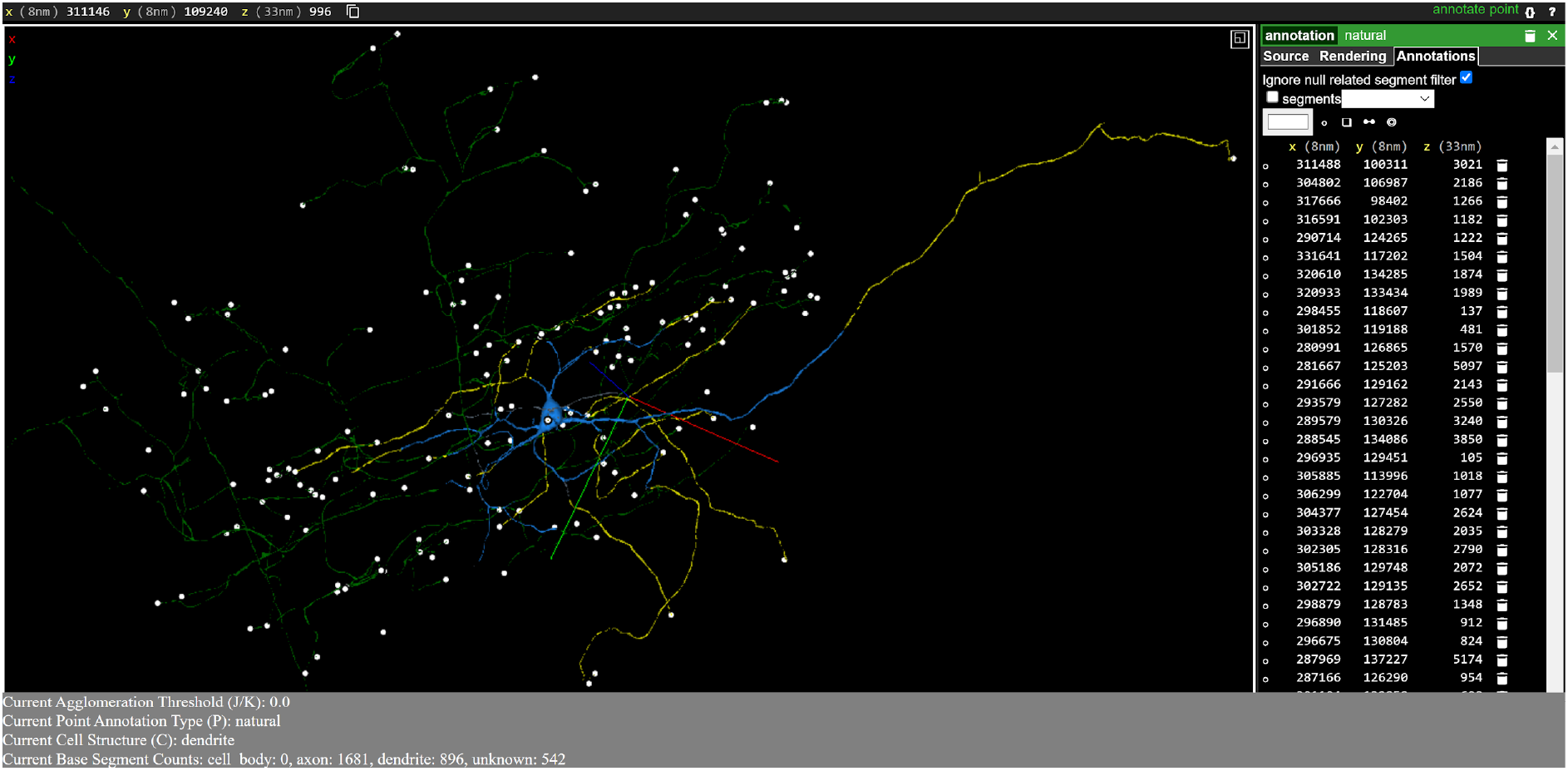
Neuron proofreading interface in Neuroglancer with CREST. A fully proofread neuron is shown, with added dendritic and axonal base segments indicated in yellow and green, respectively. The base segment containing the soma is indicated in blue. White dots are point annotations added to indicate the ends of branches. Numbers of classified and unclassified base segments are displayed in a count near the bottom of the screen.

**Supplementary Figure 16.**
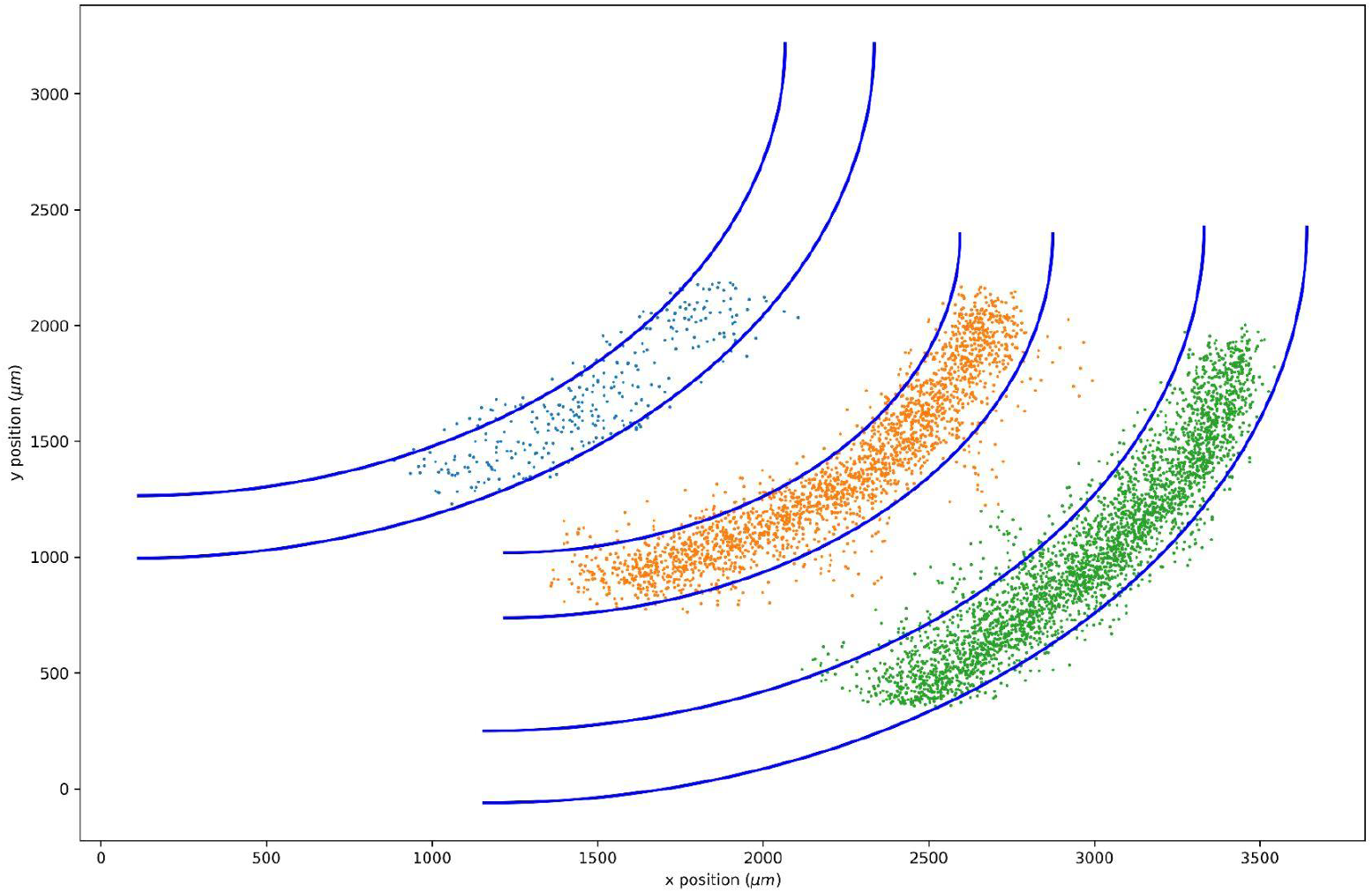
Clusters of neurons with fitted upper and lower bounds. Blue lines: fitted layer bounds. Points: individual neurons defining a cluster, colored by cluster membership.

**Supplementary Figure 17.**
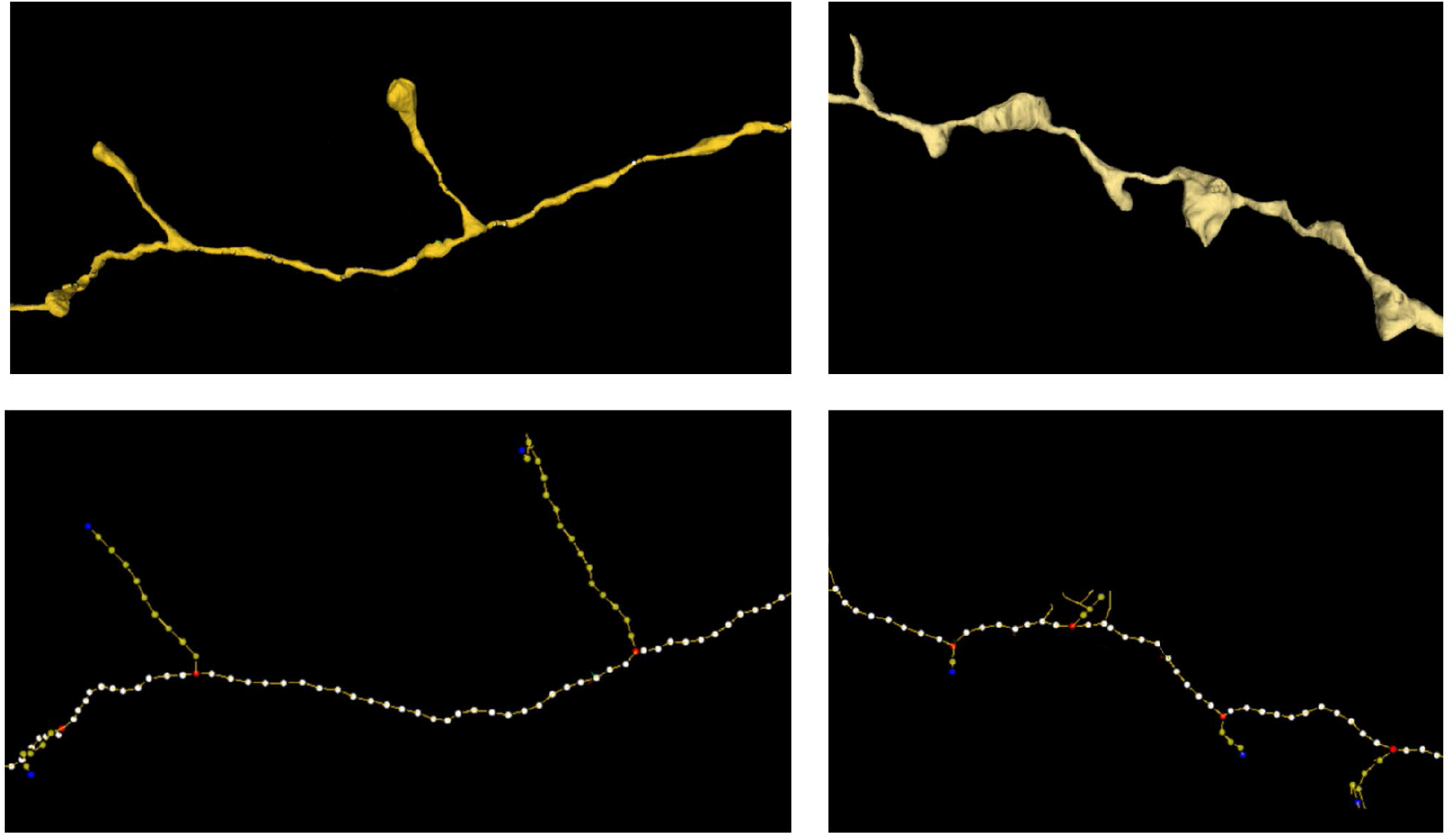
Identification of shaft, synapse and terminal bouton stalk components of axonal skeletons. Upper panels: Segmentation views of axons. Lower panels: Skeleton views of axons. Right panels: axon with three ‘en-passant’ synapses. Left panels: axon with two terminal bouton synapses and one ‘en passant’ synapse. Blue dots: synapse-associated skeleton nodes. White dots: shaft skeleton nodes. Yellow dots: skeleton nodes connecting synapse nodes to the shaft.

**Supplementary Figure 18.**
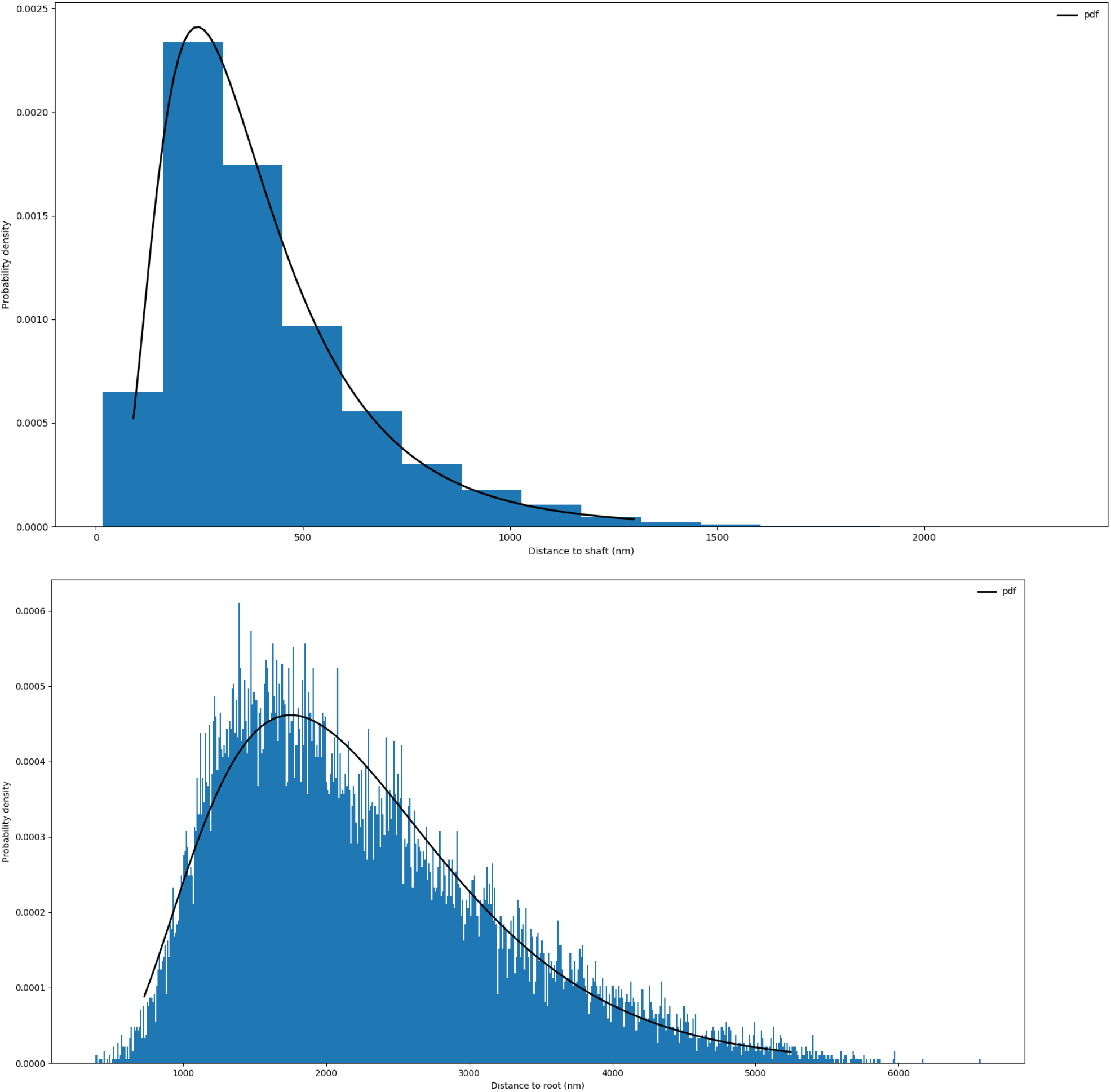
Distribution of synapse distances from axonal shaft. Blue columns indicate distribution of observed distances and the black curve indicates the fitted probability density function. Upper panel: distribution of distances from synapse location to closest point on shaft for ‘en passant’ (‘shaft-type’) synapses, with fitted scaled, shifted lognormal distribution (shape: 0.577, location: 2.42, scale: 338, see https://docs.scipy.org/doc/scipy/reference/generated/scipy.stats.lognorm.html for details). Lower panel: Distribution of distances from synapse location to ‘root point’ on shaft for ‘terminal bouton stalk’’ (‘stalk-type’) synapses, with fitted scaled, shifted chi-sq distribution (df: 7.43, location: 369, scale: 254, see https://docs.scipy.org/doc/scipy/reference/generated/scipy.stats.chi2.html for details).

**Supplementary Figure 19.**
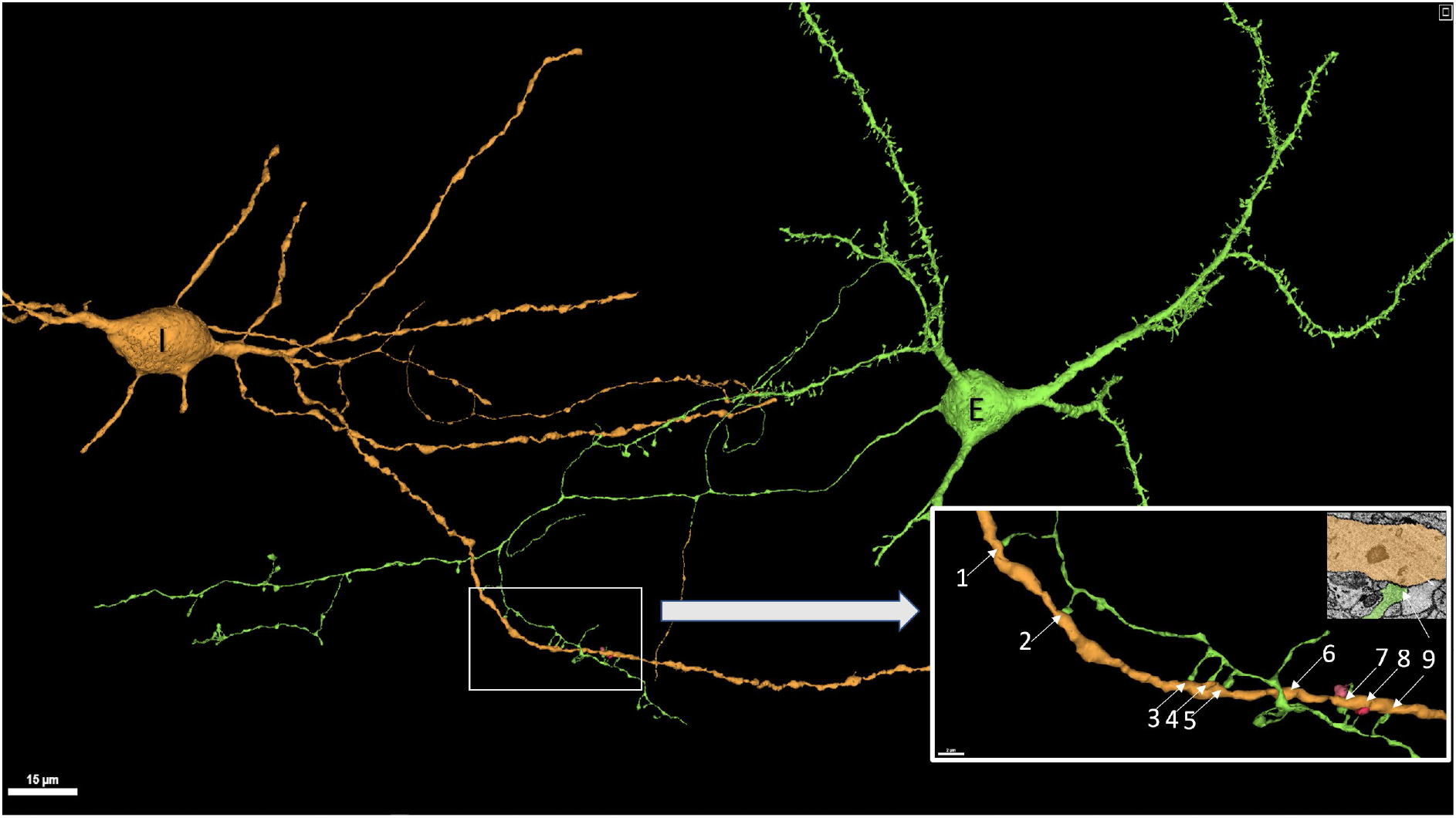
Excitatory cell making multiple synapses on inhibitory cell dendrite. E: excitatory cell. I: inhibitory cell. Numbered arrows: individual synapses.

**Supplementary Figure 20.**
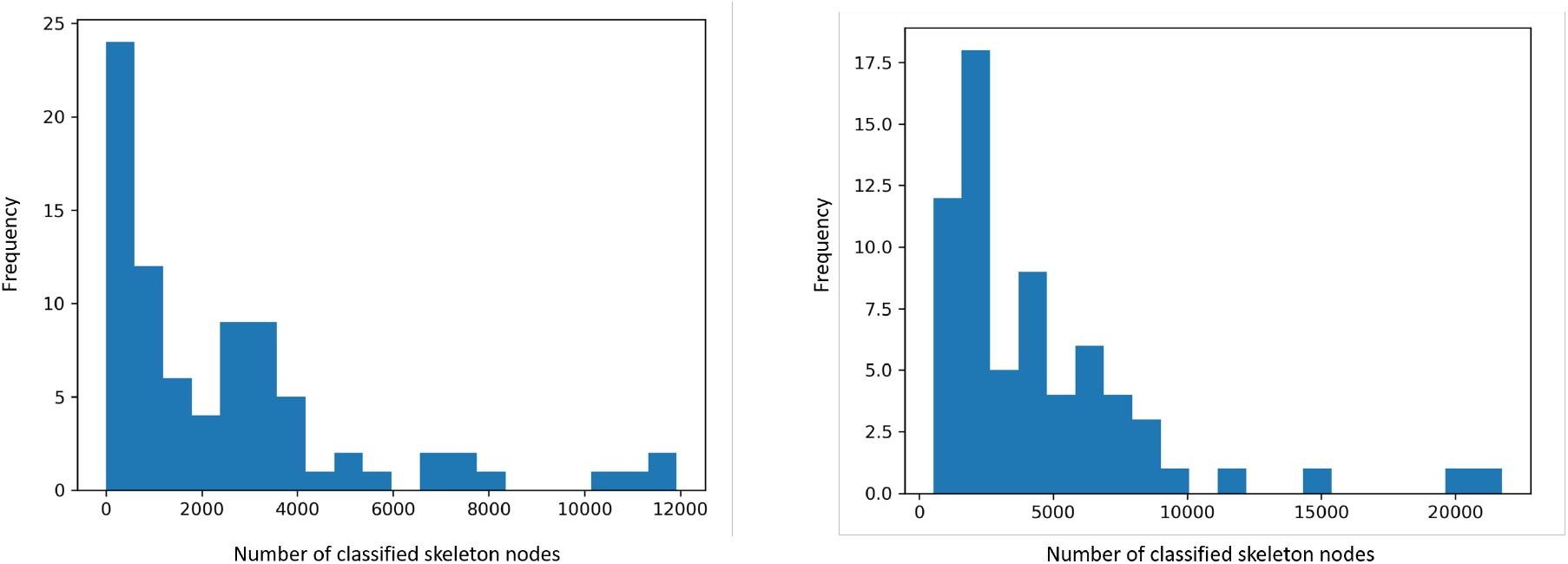
Difference in number of classified skeleton nodes in ‘forward’ and ‘reverse’ layer 6 pyramidal neuron basal dendrites. ‘Reverse’ basal dendrites (upper panel) have less mass as measured by number of classified skeleton nodes (mean: 2514) than ‘forward’ basal dendrites (lower panel, mean: 4437). Skeleton nodes are classified as described in ‘Cellular subcompartment classification and merge error correction’.

**Supplementary Figure 21:**
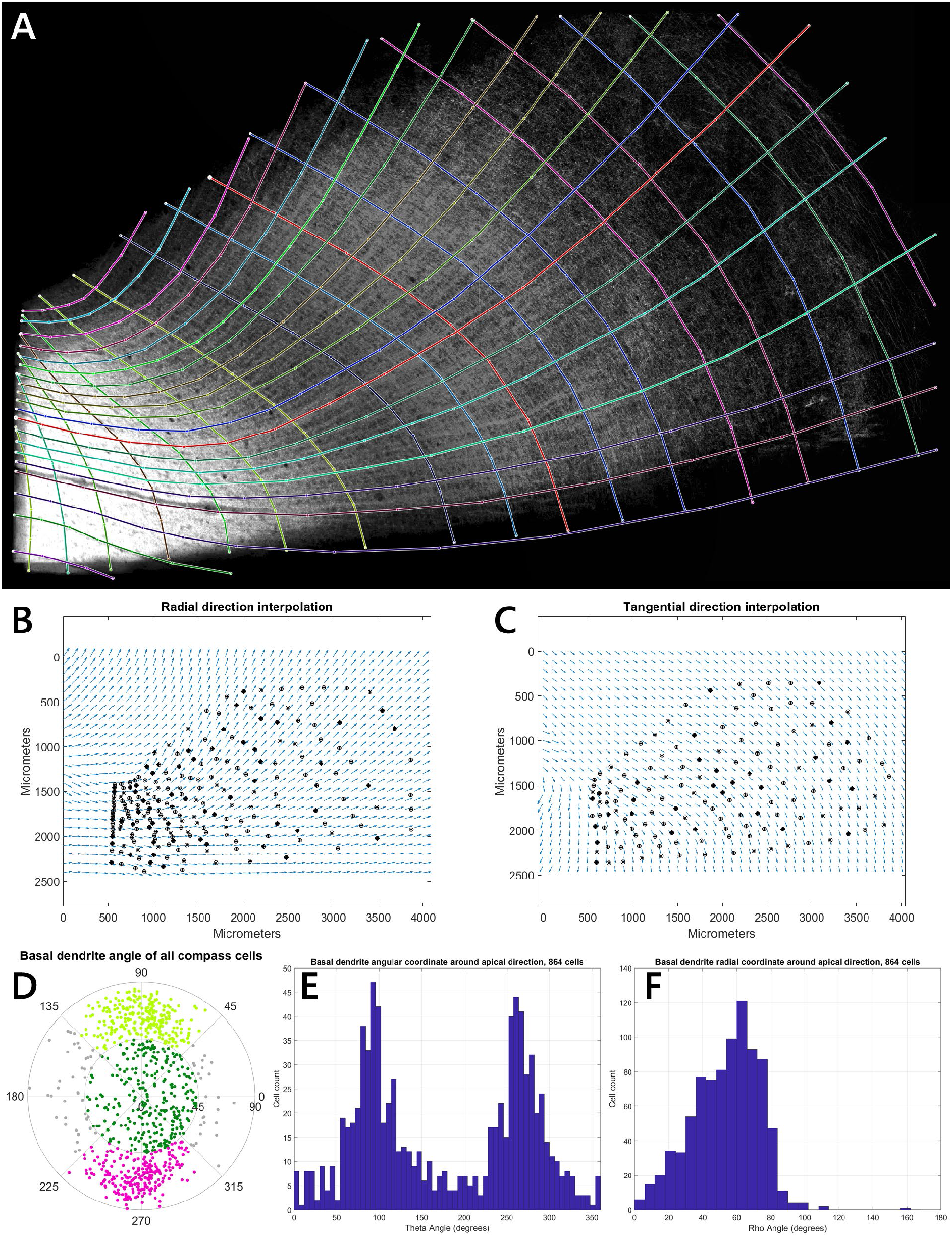
Layer 5/6 triangular (or compass) cells. **(A)** Estimation of tissue topology by fitting a grid of tangential and radial lines to 2D projection images of different extracted properties of the sample in VAST. **(B)**, **(C)** Interpolated vector fields from these lines. **(D)-(F)** Basal dendrite directions around the local radial direction with respect to the topology field, as polar plot (D) and histograms of angular (E) and radial (F) coordinate.

**Supplementary Figure 22:**
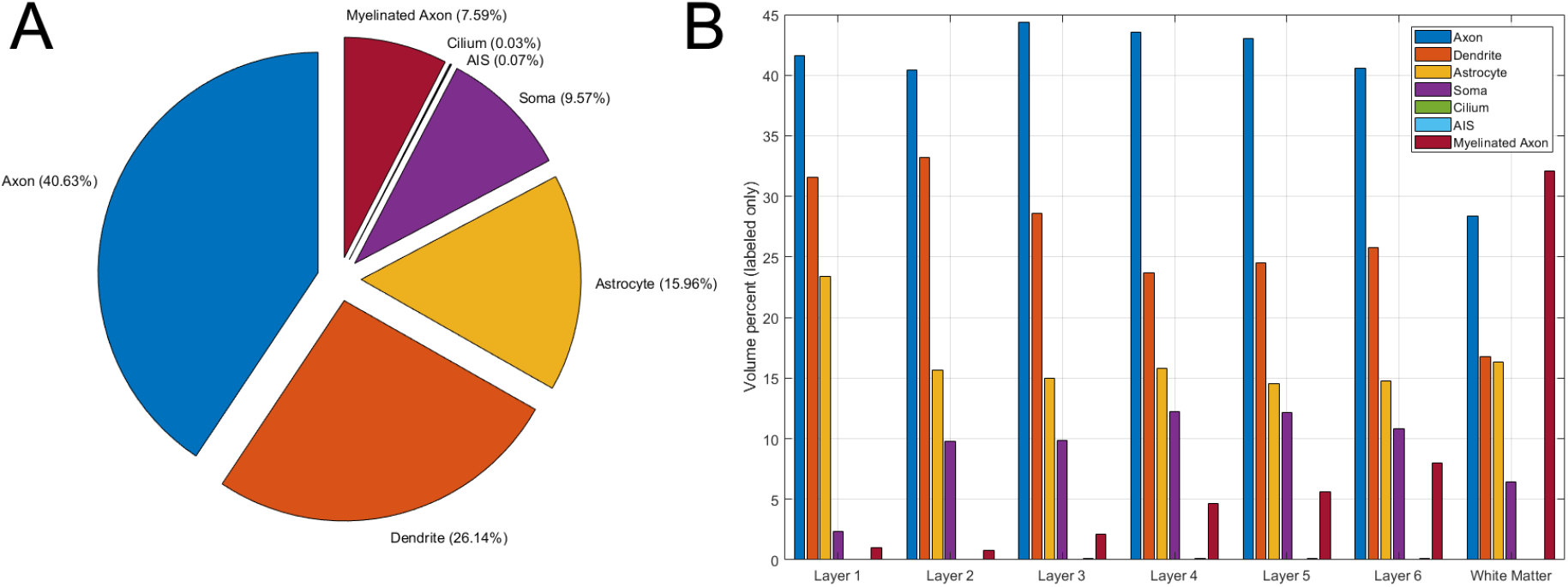
Volume occupancy by layer. All objects in the C3 segmentation were classified into seven classes based on the classification of their skeletons. The resulting voxel data was sampled at 2048 x 2048 x 2112 nm per pixel (mip 8) and masked by a manually generated tissue mask to remove all voxels outside the segmented volume. **(A)** Overall statistics of the volume percentage of different classes in the complete segmented volume. 36.82% of the voxels within the tissue were unlabeled and are excluded from the analysis. Unlabeled regions include: blood vessels, myelin sheaths, regions of fissures or other image defects, and extracellular space. **(B)** Since tissue statistics vary by cortical layer, we split the volume into regions based on the circular layer estimates (see **Supplementary Fig. 16**) and analyzed volume occupancy separately per layer. See **Supplementary Table 9** for the data values.

**Supplementary Figure 23:**
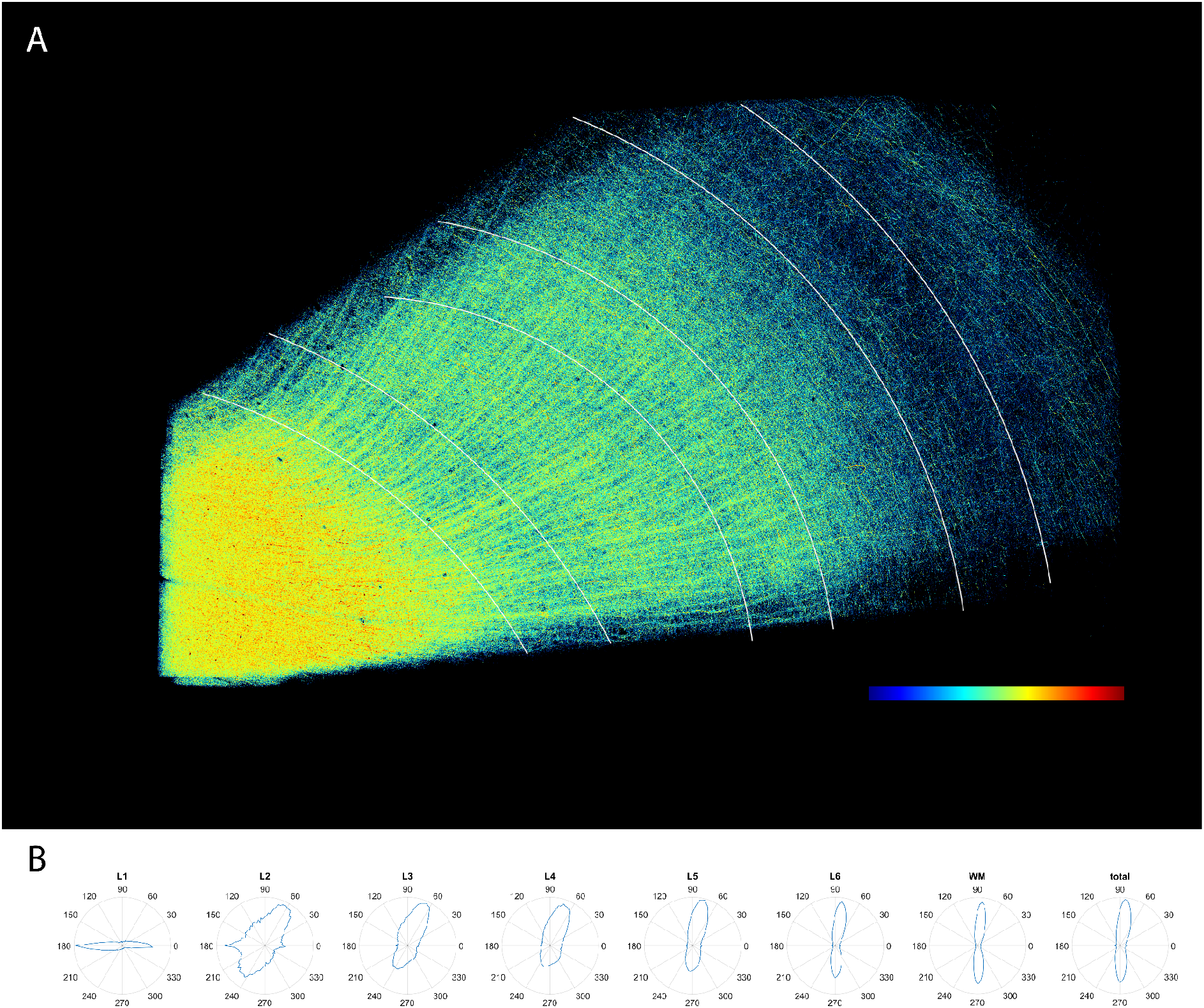
Myelinated axons from skeletonization. Statistics about the myelinated axons were extracted from skeletonization. **(A)** Maximum projection view of the myelinated axon calibers (excluding myelin sheath). The colors were mapped linearly to the axon calibers, with the blue end corresponding to 0 and red end corresponding to a caliber of 1 micron. **(B)** The angular distribution of the myelinated axons by layer in the perpendicular-tangential plane defined in **Fig. 3H**. The vertical axes of the polar plots correspond to the perpendicular direction (perpendicular to the sectioning plane) and the horizontal axes correspond to the tangential direction (tangential to the pia). Only axons with an angle larger than 45 degrees away from the radial direction were counted.

**Supplementary Figure 24:**
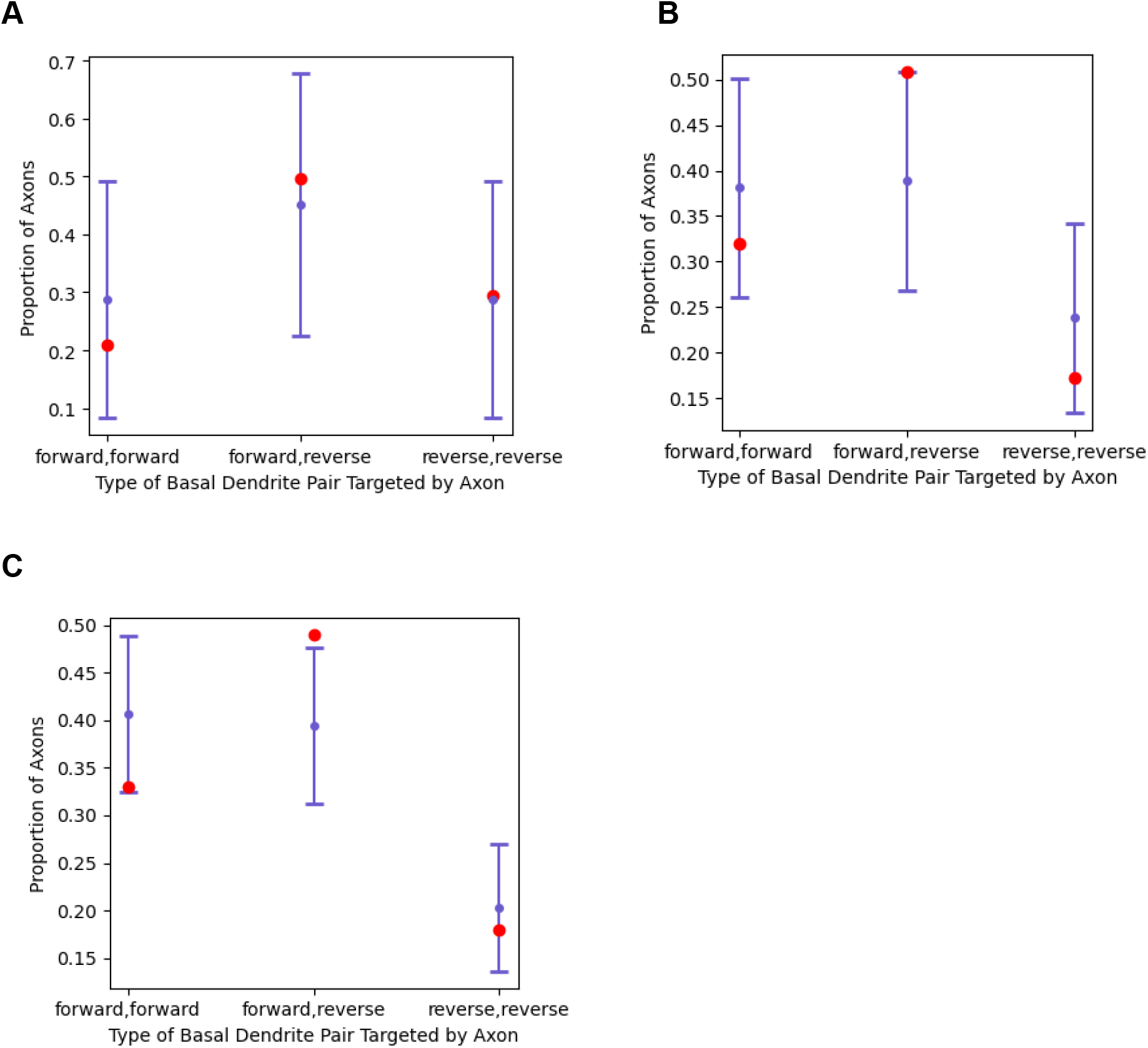
Basal dendrite type pair preferences of basal dendrite-targeting axons. **(A)** Axon initial segment (AIS) targeting inhibitory axons (p = 0.319, n = 174). **(B)** Largest basal dendrite (non-AIS) targeting inhibitory axons (p = 9.89 · 10^-6^, n=372). **(C)** Largest basal dendrite (non-AIS) targeting excitatory axons (p = 1.60 · 10^-7^, n=808). Red dots indicate expected proportions, blue bars indicate 95% confidence intervals around observed proportions, P-values are calculated using the Chi-Squared Test to compare expected and observed counts.

**Supplementary Figure 25:**
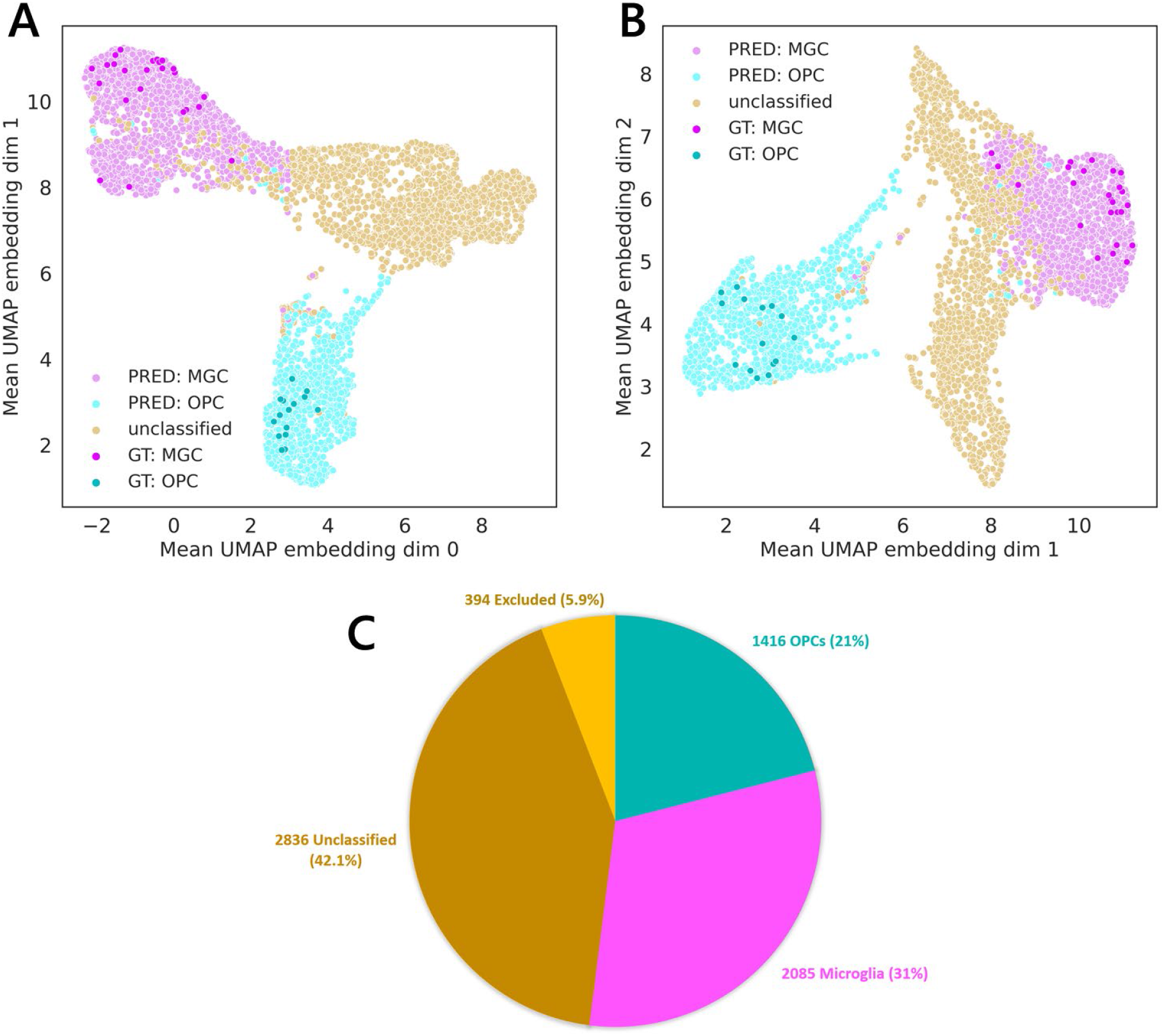
Classifying Microglia (MGCs) vs. Oligodendrocyte Precursor Cells (OPCs) by using linear embeddings. (A), (B): Two-dimensional views of three-dimensional UMAP space derived from the 32-dimensional embedding space. Ground-truth examples used for training a linear classifier shown in darker color. ‘Unclassified’ cells could not be reliably classified as either microglia or OPC. **(C)**: Pie chart shows that more than half of the 6731 candidate cells could be classified by this method.

**Supplementary Figure 26:**
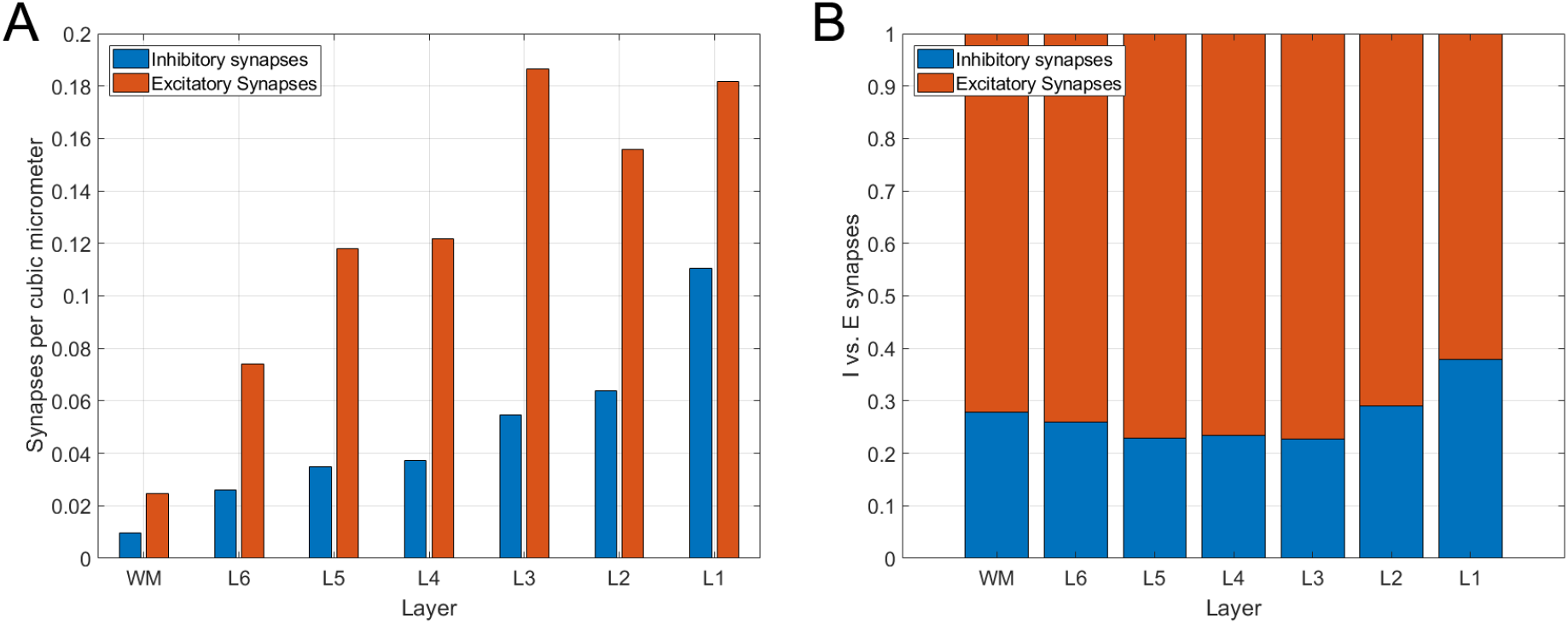
Excitatory and inhibitory synapses in different layers. (A): Densities of excitatory and inhibitory synapses in different layers. The values of this plot can be found in Supplementary Table 10. **(B):** E/I balance of synapse numbers in different layers. This analysis includes all automatically detected synapses within a region of 0.65 mm^3^ of the dataset, using the layer boundaries and volumetric tissue mask used for estimating cell densities (Supplementary Figure 5). This statistics is based on 116.92 million synapses, of which 86.08 million were classified as excitatory (73.6%) and 30.84 million as inhibitory (26.4%). The average number of synapses is 0.1789 per μm^3^ in the analyzed volume.

## Supplementary Tables

**Supplementary Table 1.** Imaging metrics. See <u>h01_imaging_metrics.csv</u>

**Supplementary Table 2.**
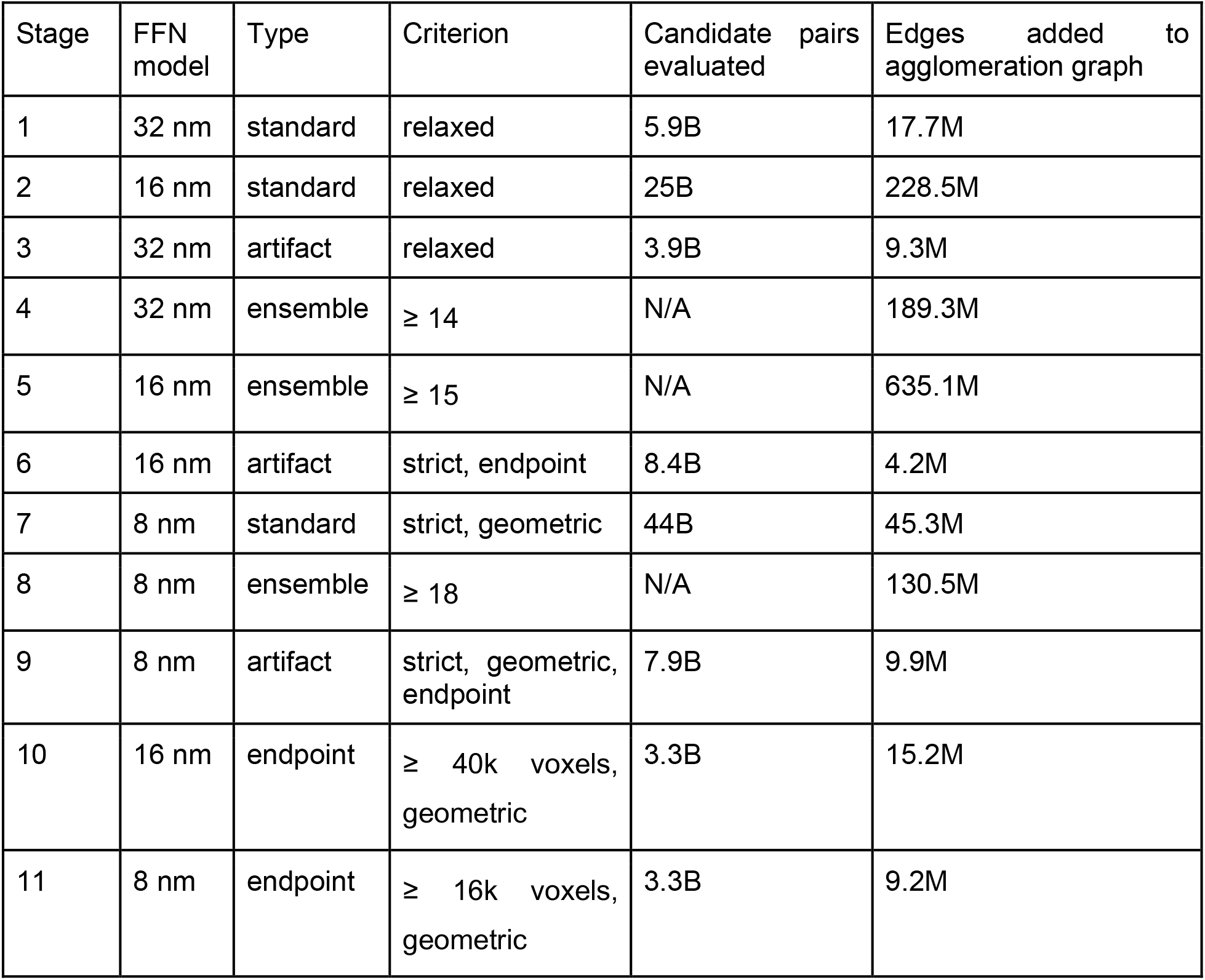
Agglomeration graph edge selection criteria by agglomeration stage. The criteria are: strict = ((dA ≤ 0.02 ∨ dB ≤ 0.02) ∧ (f** ≥ 0.6 ∧ JAB ≥ 0.8)), relaxed = strict ∨ (f** ≥ 0.9 ∧ JAB ≥ 0.9). Ensemble criteria refer to the number of alternative segmentations contributing a given (A, B) Candidate edge. Endpoint size criteria refer to object sizes in the base segmentation. The endpoint criterion excluded all segment pairs for which skeleton endpoints for both segments in the pair were not present within a 500 nm of the pair center. Geometric criteria excluded points for which y < 181053 + (276044 - 181053) * ((x - 104865) / (204645 - 104865)) (white matter, only applied when the 8 nm FFN model was used), (x - (328769 + (z - 1919) / (5251 - 1919) * (343919 - 328769))2 + (y - (75082 + (z - 1919) / (5251 - 1919) * (65206 - 75082))2 >= 37452 (fissure), or (x - (338167 + (z - 1904) / (5286 - 1904) * (339636 - 338167))2 + (y - (138052 + (z - 1904) / (5286 - 1904) * (142197 - 138052))2 >= 17592 (fissure).

| Stage | FFN model | Type | Criterion | Candidate pairs evaluated | Edges added to agglomeration graph |
| --- | --- | --- | --- | --- | --- |
| 1 | 32 nm | standard | relaxed | 5.9B | 17.7M |
| 2 | 16 nm | standard | relaxed | 25B | 228.5M |
| 3 | 32 nm | artifact | relaxed | 3.9B | 9.3M |
| 4 | 32 nm | ensemble | $\geq 14$ | N/A | 189.3M |
| 5 | 16 nm | ensemble | $\geq 15$ | N/A | 635.1M |
| 6 | 16 nm | artifact | strict, endpoint | 8.4B | 4.2M |
| 7 | 8 nm | standard | strict, geometric | 44B | 45.3M |
| 8 | 8 nm | ensemble | $\geq 18$ | N/A | 130.5M |
| 9 | 8 nm | artifact | strict, geometric, endpoint | 7.9B | 9.9M |
| 10 | 16 nm | endpoint | $\geq 40k$ voxels, geometric | 3.3B | 15.2M |
| 11 | 8 nm | endpoint | $\geq 16k$ voxels, geometric | 3.3B | 9.2M |

**Supplementary Table 3.** Manual counts of all cells in H01, Phase 1 & 2.

|  |  |  |  |
| --- | --- | --- | --- |
| Pyramidal Cells [P] |  |  | 8803 |
| Spiny Cells [E] |  |  | 1535 |
| Spiny Stellate [S] |  |  | 193 |
| Interneurons [I] |  |  | 4688 |
| Unclassified Neurons [N] |  |  | 868 |
| Astrocytes [A] |  |  | 5474 |
| Oligodendrocytes [O] |  |  | 20139 |
| Microglia / OPC [G] |  |  | 6536 |
| C-shaped MG/OPC [C] |  |  | 166 |
| Unclassified Cells [U] |  |  | 678 |
| All Excitatory Neurons |  |  | 10531 |
| All Inhibitory Neurons |  |  | 4688 |
| All Neurons |  |  | 16087 |
| All Glia |  |  | 32315 |
| Blood Vessel Cells in DB cells |  |  | 285 |
| BV Endothelial Cells |  |  | 4604 |
| BV Pericytes |  |  | 3549 |
| BV Circulating Immune |  |  | 46 |
| All DB Cells w/o BV |  |  | 49080 |
| All ES BV cells |  |  | 8136 |
| All Cells |  |  | 57216 |
| E / (E+I) |  |  | 0.692 |
| I / (E+I) |  |  | 0.308 |

**Supplementary Table 4.**
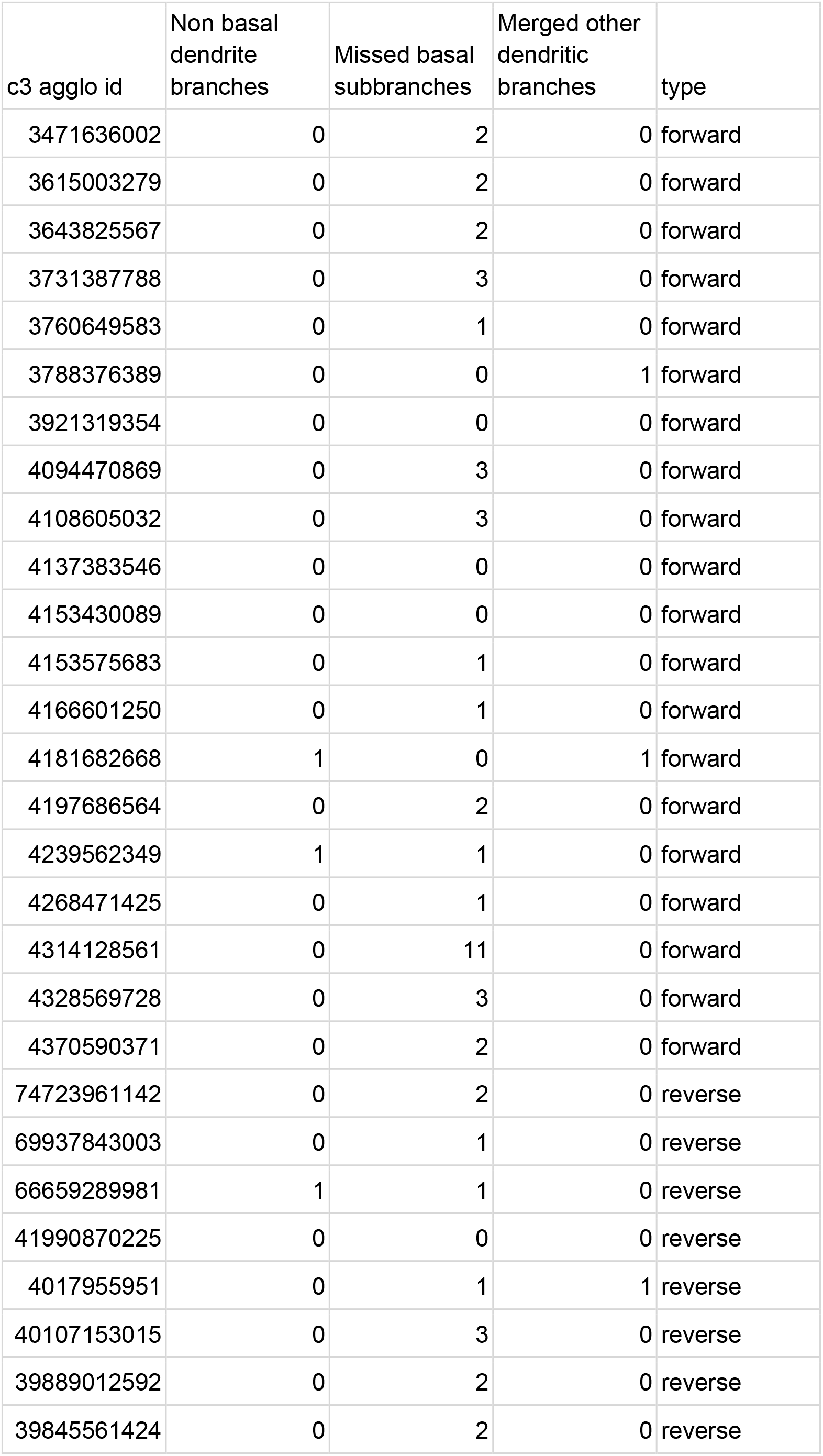
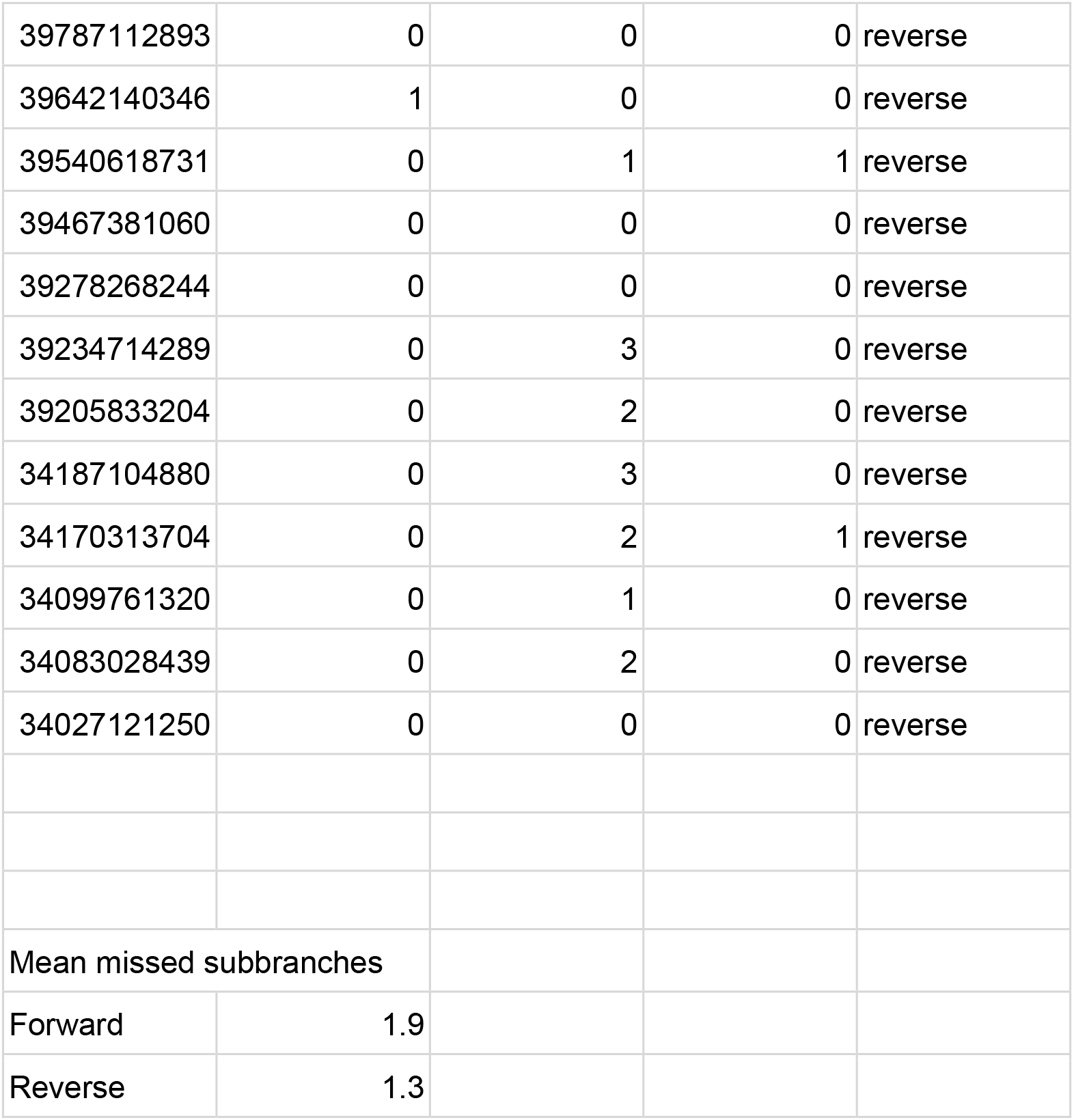
Basal dendrite branch identification performance.

**Supplementary Table 5.**
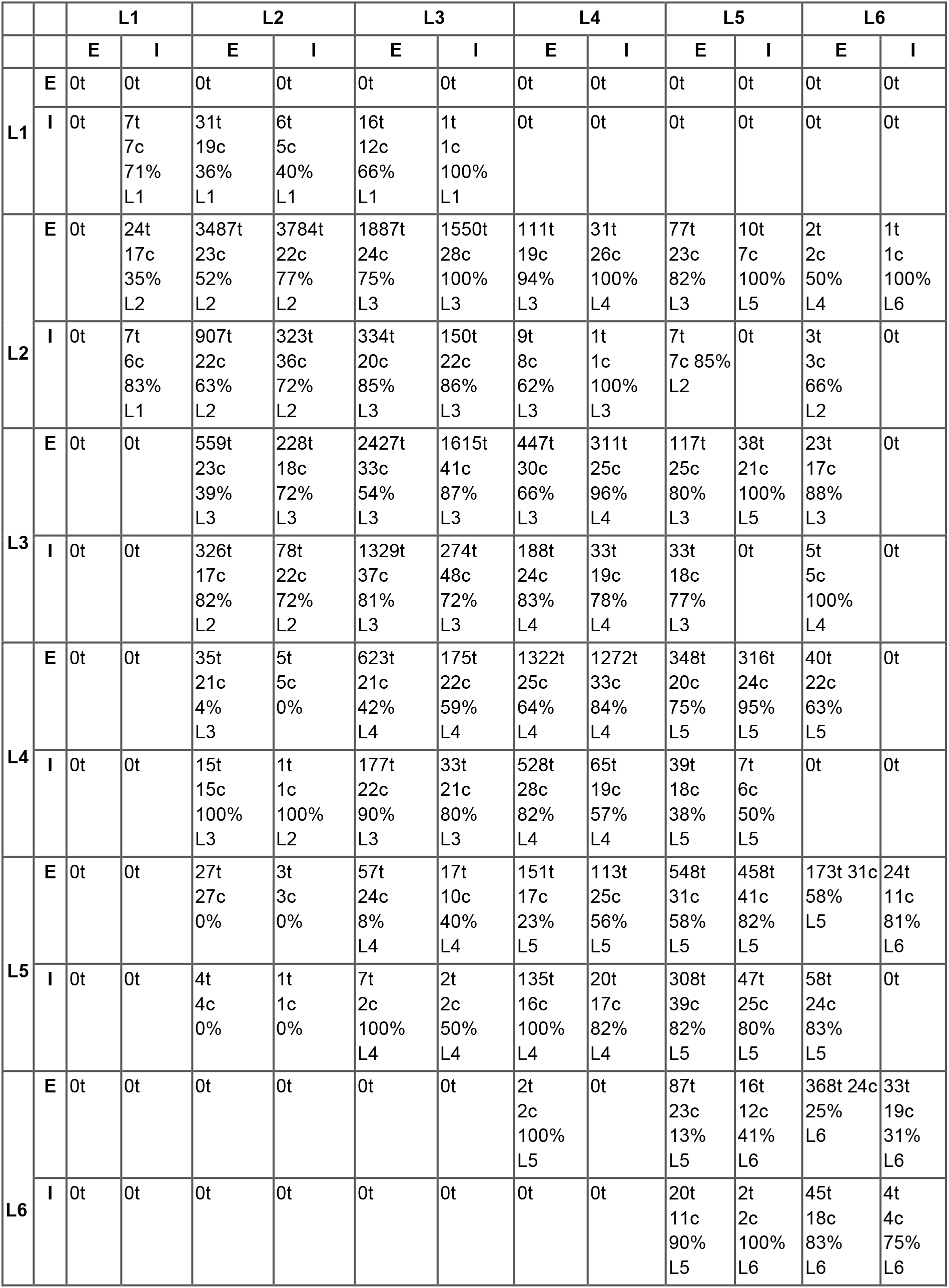
Summary of Machine Learning-identified connections. Pre- and post-synaptic cell type (E or I) and Layer (L) are indicated in row and column headings, respectively. Each cell contains the total number of connections (t), number manually checked (c), the percentage of those checked that are valid, and the most common layer for the connection to occur in (L), for that connection type.

**Supplementary Table 6.** Summary of manually-identified connections. Rows indicate pre-synaptic neuron types and columns indicate post-synaptic neuron types. L numbers indicate layer membership of cell and E and I indicate excitatory and inhibitory cell types, respectively. Counts indicate numbers of connections between the pre-synaptic type indicated by the row and the post-synaptic type indicated by the column, where one connection consists of all of the synapses between two individual neurons

|  |  |  | Post-synaptic |  |  |  |  |  |  |  |  |  |  |  |
| --- | --- | --- | --- | --- | --- | --- | --- | --- | --- | --- | --- | --- | --- | --- |
|  |  |  | L1 |  | L2 |  | L3 |  | L4 |  | L5 |  | L6 |  |
|  |  |  | E | I | E | I | E | I | E | I | E | I | E | I |
| Pre-synaptic | L1 | E | 0 | 0 | 0 | 0 | 0 | 0 | 0 | 0 | 0 | 0 | 0 | 0 |
|  |  | I | 0 | 0 | 0 | 0 | 0 | 0 | 0 | 0 | 0 | 0 | 0 | 0 |
|  | L2 | E | 0 | 0 | 45 | 60 | 18 | 3 | 1 | 0 | 0 | 0 | 0 | 0 |
|  |  | I | 0 | 0 | 27 | 8 | 6 | 1 | 0 | 0 | 0 | 0 | 0 | 0 |
|  | L3 | E | 0 | 0 | 0 | 0 | 9 | 20 | 7 | 6 | 0 | 1 | 1 | 0 |
|  |  | I | 0 | 0 | 18 | 2 | 76 | 11 | 7 | 1 | 1 | 0 | 0 | 0 |
|  | L4 | E | 0 | 0 | 0 | 0 | 18 | 7 | 27 | 27 | 3 | 4 | 0 | 0 |
|  |  | I | 0 | 0 | 0 | 0 | 0 | 0 | 0 | 1 | 0 | 0 | 0 | 0 |
|  | L5 | E | 0 | 0 | 0 | 0 | 0 | 0 | 0 | 0 | 15 | 13 | 3 | 0 |
|  |  | I | 0 | 0 | 2 | 1 | 9 | 1 | 24 | 3 | 18 | 3 | 8 | 0 |
|  | L6 | E | 0 | 0 | 0 | 0 | 0 | 0 | 0 | 0 | 0 | 0 | 1 | 0 |
|  |  | I | 0 | 0 | 0 | 0 | 0 | 0 | 0 | 0 | 0 | 0 | 0 | 0 |

**Supplementary Table 7.**
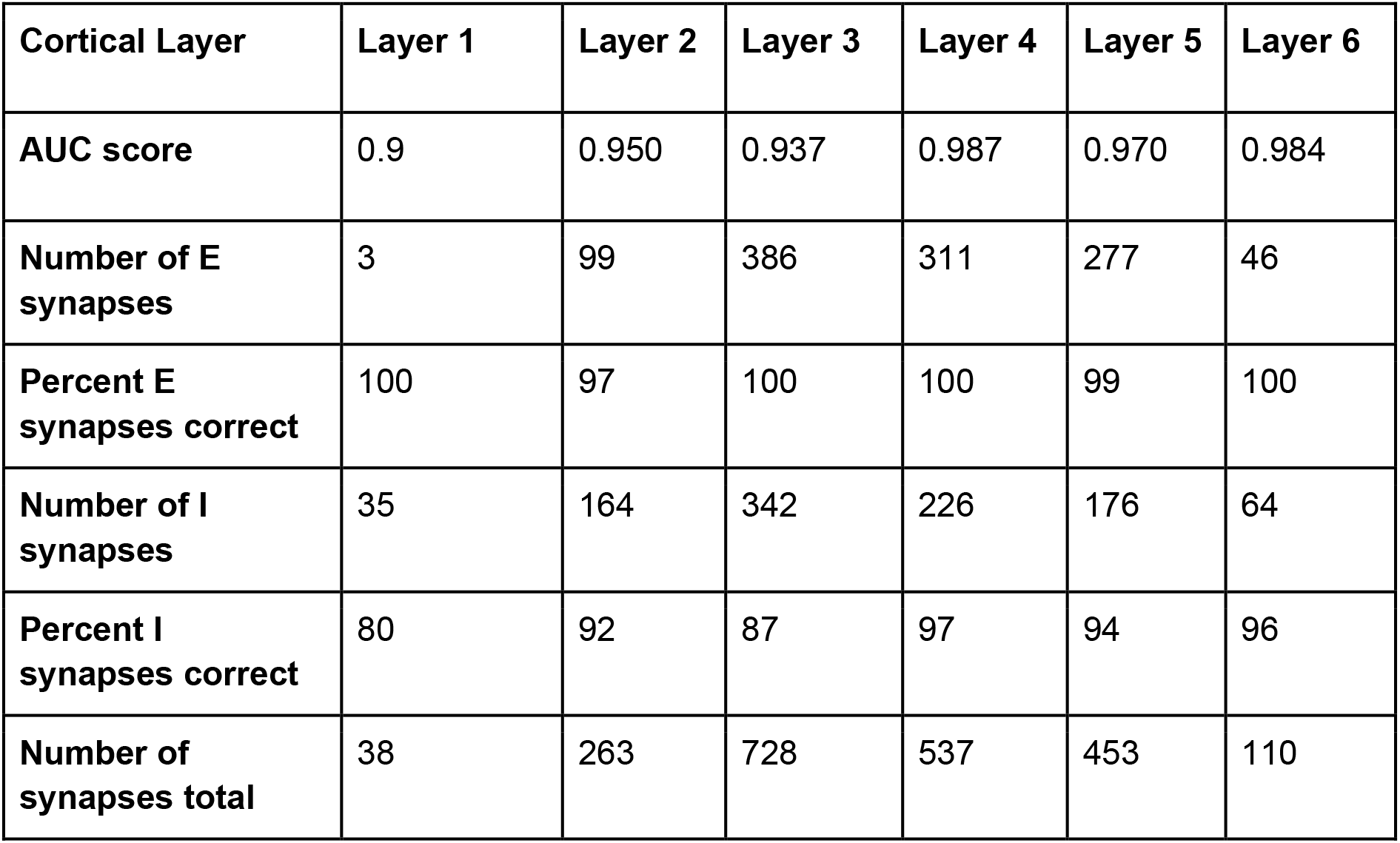
Excitatory and inhibitory synapse classification accuracy. Columns indicate cortical layers in which synapses were formed. E and I refer to excitatory and inhibitory, respectively.

**Supplementary Table 8.** Estimated number of cells per cubic millimeter for different layers. These are the values of the plot shown in Supplementary Figure 5A.

|  | Layer 1 | Layer 2 | Layer 3 | Layer 4 | Layer 5 | Layer 6 | White Matter |
| --- | --- | --- | --- | --- | --- | --- | --- |
| Spiny Neurons | 36.05 | 21194 | 8628.3 | 19724 | 10072 | 7678.4 | 2189.6 |
| Interneurons | 5299.3 | 11522 | 4980.8 | 5184.2 | 2905.8 | 1220.4 | 679.52 |
| Unclassified Neurons | 144.2 | 624.51 | 226.4 | 678.22 | 463.25 | 254.25 | 616.6 |
| Astrocytes | 12293 | 3567.5 | 4427.4 | 4589.6 | 3986.7 | 3496 | 4089.7 |
| Oligodendrocytes | 5641.8 | 3942.2 | 8540.3 | 14131 | 16880 | 24141 | 61471 |
| Microglia/OPCs | 6326.7 | 6510.5 | 6490.1 | 6001.8 | 5790.6 | 6038.5 | 8053.6 |
| Unknown | 198.27 | 234.19 | 176.09 | 761.83 | 1052.8 | 978.87 | 478.18 |

**Supplementary Table 9.** Estimated volume occupancy percentages for different layers. These are the values of the plot shown in Supplementary Figure 22B.

|  | Layer 1 | Layer 2 | Layer 3 | Layer 4 | Layer 5 | Layer 6 | White Matter |
| --- | --- | --- | --- | --- | --- | --- | --- |
| Axon | 41.6515 | 40.4082 | 44.3686 | 43.5128 | 43.0305 | 40.5566 | 28.3573 |
| Dendrite | 31.5978 | 33.2242 | 28.5824 | 23.6832 | 24.4849 | 25.786 | 16.7716 |
| Astrocyte | 23.3594 | 15.664 | 14.9801 | 15.7906 | 14.5567 | 14.73 | 16.2967 |
| Soma | 2.3623 | 9.7743 | 9.8245 | 12.2161 | 12.1522 | 10.8323 | 6.4454 |
| Cilium | 0.0068 | 0.0498 | 0.0308 | 0.0561 | 0.0433 | 0.0269 | 0.0135 |
| AIS | 0.02 | 0.0706 | 0.0861 | 0.0979 | 0.0908 | 0.0874 | 0.0299 |
| Myelinated Axon | 1.0022 | 0.8088 | 2.1276 | 4.6433 | 5.6416 | 7.9809 | 32.0855 |

**Supplementary Table 10.** Estimated synapse densities for different layers. These are the values of the plot shown in Supplementary Figure 26A.

|  | Layer 1 | Layer 2 | Layer 3 | Layer 4 | Layer 5 | Layer 6 | White Matter |
| --- | --- | --- | --- | --- | --- | --- | --- |
| Excitatory synapses per $\mu\text{m}^3$ | 0.1817 | 0.15596 | 0.18654 | 0.12195 | 0.1179 | 0.074138 | 0.024861 |
| Inhibitory synapses per $\mu\text{m}^3$ | 0.11062 | 0.063762 | 0.054729 | 0.037229 | 0.034986 | 0.026043 | 0.0095933 |

## References

1. Zhou, Y. et al. Atypical behaviour and connectivity in SHANK3-mutant macaques. Nature 570, 326–331 (2019).

2. Birey, F. et al. Assembly of functionally integrated human forebrain spheroids. Nature 545, 54–59 (2017).

3. Quadrato, G. et al. Cell diversity and network dynamics in photosensitive human brain organoids. Nature 545, 48–53 (2017).

4. Velasco, S. et al. Individual brain organoids reproducibly form cell diversity of the human cerebral cortex. Nature vol. 570 523–527 (2019).

5. Engel, J. Early Surgical Therapy for Drug-Resistant Temporal Lobe Epilepsy. JAMA vol. 307 922 (2012).

6. . Tapia, J. C., et al. High-contrast en bloc staining of neuronal tissue for field emission scanning electron microscopy. Nat. Protoc. 7, 193–206 (2012).

7. Hayworth, K. J. et al. Imaging ATUM ultrathin section libraries with WaferMapper: a multi-scale approach to EM reconstruction of neural circuits. Front. Neural Circuits 8, 68 (2014).

8. Saalfeld, S., Fetter, R., Cardona, A. & Tomancak, P. Elastic volume reconstruction from series of ultra-thin microscopy sections. Nat. Methods 9, 717–720 (2012).

9. Januszewski, M. et al. High-precision automated reconstruction of neurons with flood- filling networks. Nat. Methods 15, 605–610 (2018).

10. Li, H., Januszewski, M., Jain, V. & Li, P. H. Neuronal Subcompartment Classification and Merge Error Correction. doi:10.1101/2020.04.16.043398.

11. Keller, D., Erö, C. & Markram, H. Cell Densities in the Mouse Brain: A Systematic Review. Front. Neuroanat. 12, 83 (2018).

12. Bae, J. A., Baptiste, M., Bodor, A. L., Brittain, D. & Buchanan, J. A. Functional connectomics spanning multiple areas of mouse visual cortex. bioRxiv (2021).

13. Lanjakornsiripan, D. et al. Layer-specific morphological and molecular differences in neocortical astrocytes and their dependence on neuronal layers. Nat. Commun. 9, 1623 (2018).

14. Hodge, R. D. et al. Conserved cell types with divergent features in human versus mouse cortex. Nature 573, 61–68 (2019).

15. Bayraktar, O. A. et al. Astrocyte layers in the mammalian cerebral cortex revealed by a single-cell in situ transcriptomic map. Nat. Neurosci. 23, 500–509 (2020).

16. Colombo, J. A. & Reisin, H. D. Interlaminar astroglia of the cerebral cortex: a marker of the primate brain. Brain Res. 1006, 126–131 (2004).

17. Bushong, E. A., Martone, M. E., Jones, Y. Z. & Ellisman, M. H. Protoplasmic astrocytes in CA1 stratum radiatum occupy separate anatomical domains. J. Neurosci. 22, 183– 192 (2002).

18. Ogata, K. & Kosaka, T. Structural and quantitative analysis of astrocytes in the mouse hippocampus. Neuroscience 113, 221–233 (2002).

19. Halassa, M. M., Fellin, T., Takano, H., Dong, J.-H. & Haydon, P. G. Synaptic islands defined by the territory of a single astrocyte. J. Neurosci. 27, 6473–6477 (2007).

20. Marques, S. et al. Oligodendrocyte heterogeneity in the mouse juvenile and adult central nervous system. Science 352, 1326–1329 (2016).

21. Harris, J. J. & Attwell, D. The energetics of CNS white matter. J. Neurosci. 32, 356–371 (2012).

22. Vanlandewijck, M. et al. A molecular atlas of cell types and zonation in the brain vasculature. Nature 554, 475–480 (2018).

23. Hall, C. N. et al. Capillary pericytes regulate cerebral blood flow in health and disease. Nature 508, 55–60 (2014).

24. Hartmann, D. A. et al. Brain capillary pericytes exert a substantial but slow influence on blood flow. Nat. Neurosci. 24, 633–645 (2021).

25. Lapenna, A., De Palma, M. & Lewis, C. E. Perivascular macrophages in health and disease. Nat. Rev. Immunol. 18, 689–702 (2018).

26. Duvernoy, H. M., Delon, S. & Vannson, J. L. Cortical blood vessels of the human brain. Brain Res. Bull. 7, 519–579 (1981).

27. Brown, W. R. A Review of String Vessels or Collapsed, Empty Basement Membrane Tubes. Journal of Alzheimer’s Disease vol. 21 725–739 (2010).

28. Mendes-Jorge, L. et al. Intercapillary bridging cells: immunocytochemical characteristics of cells that connect blood vessels in the retina. Exp. Eye Res. 98, 79–87 (2012).

29. Alarcon-Martinez, L. et al. Interpericyte tunnelling nanotubes regulate neurovascular coupling. Nature 585, 91–95 (2020).

30. Soares, J. M., Marques, P., Alves, V. & Sousa, N. A hitchhiker’s guide to diffusion tensor imaging. Front. Neurosci. 7, 31 (2013).

31. DeFelipe, J., Alonso-Nanclares, L. & Arellano, J. I. Microstructure of the neocortex: comparative aspects. J. Neurocytol. 31, 299–316 (2002).

32. Felleman, D. J. & Van Essen, D. C. Distributed hierarchical processing in the primate cerebral cortex. Cereb. Cortex 1, 1–47 (1991).

33. Callaway, E. M. Local circuits in primary visual cortex of the macaque monkey. Annu. Rev. Neurosci. 21, 47–74 (1998).

34. Szentágothai, J. The ‘module-concept’ in cerebral cortex architecture. Brain Res. 95, 475–496 (1975).

35. Schneider-Mizell, C. M. et al. Chandelier cell anatomy and function reveal a variably distributed but common signal. doi:10.1101/2020.03.31.018952.

36. Briggs, F. Organizing principles of cortical layer 6. Front. Neural Circuits 4, 3 (2010).

37. Callaway, E. M. Cell types and local circuits in primary visual cortex of the macaque monkey. The visual neurosciences 1, 680–694 (2004).

38. Ramón y Cajal, S. Histologie du système nerveux de l’homme et des vertébrés. (A. Maloine, 1909).

39. De Crinis, M. Uber die Spezialzellen in der menschlichen Grosshirnrinde. J Psych Neurol 45, 439–449 (1934).

40. Braak, H. Architectonics of the Human Telencephalic Cortex. Studies of Brain Function (1980) doi:10.1007/978-3-642-81522-5.

41. Burkhalter, A., Bernardo, K. L. & Charles, V. Development of local circuits in human visual cortex. J. Neurosci. 13, 1916–1931 (1993).

42. Lichtman, J. W. & Denk, W. The big and the small: challenges of imaging the brain’s circuits. Science 334, 618–623 (2011).

43. Fischl, B. & Dale, A. M. Measuring the thickness of the human cerebral cortex from magnetic resonance images. Proc. Natl. Acad. Sci. U. S. A. 97, 11050–11055 (2000).

44. Braitenberg, V. & Schüz, A. Cortex: Statistics and Geometry of Neuronal Connectivity. (Springer Science & Business Media, 2013).

45. Mishchenko, Y. Automation of 3D reconstruction of neural tissue from large volume of conventional serial section transmission electron micrographs. J. Neurosci. Methods 176, 276–289 (2009).

46. Morgan, J. L. & Lichtman, J. W. Digital tissue and what it may reveal about the brain. BMC Biol. 15, 101 (2017).

47. Kasthuri, N. et al. Saturated Reconstruction of a Volume of Neocortex. Cell 162, 648– 661 (2015).

48. Eberle, A. L. et al. High-resolution, high-throughput imaging with a multibeam scanning electron microscope. J. Microsc. 259, 114–120 (2015).

49. Briggman, K. L., Helmstaedter, M. & Denk, W. Wiring specificity in the direction-selectivity circuit of the retina. Nature 471, 183–188 (2011).

50. Bock, D. D. et al. Network anatomy and in vivo physiology of visual cortical neurons. Nature vol. 471 177–182 (2011).

51. Helmstaedter, M. et al. Connectomic reconstruction of the inner plexiform layer in the mouse retina. Nature 500, 168–174 (2013).

52. Takemura, S.-Y. et al. A visual motion detection circuit suggested by Drosophila connectomics. Nature 500, 175–181 (2013).

53. Lee, W.-C. A. et al. Anatomy and function of an excitatory network in the visual cortex. Nature 532, 370–374 (2016).

54. Motta, A. et al. Dense connectomic reconstruction in layer 4 of the somatosensory cortex. Science 366, (2019).

55. Scheffer, L. K. et al. A connectome and analysis of the adult Drosophila central brain. Elife 9, (2020).

56. Dorkenwald, S., McKellar, C., Macrina, T. & Kemnitz, N. FlyWire: Online community for whole-brain connectomics. bioRxiv (2020).

57. Yin, W. et al. A petascale automated imaging pipeline for mapping neuronal circuits with high-throughput transmission electron microscopy. Nat. Commun. 11, 4949 (2020).

58. Gour, A. et al. Postnatal connectomic development of inhibition in mouse barrel cortex. Science 371, (2021).

59. Abbott, L. F. et al. The mind of a mouse. Cell 182, 1372–1376 (2020).

60. Witvliet, D. et al. Connectomes across development reveal principles of brain maturation. Nature 596, 257–261 (2021).

61. Seung, H. S. Reading the book of memory: sparse sampling versus dense mapping of connectomes. Neuron 62, 17–29 (2009).

62. Morgan, J. L. & Lichtman, J. W. Why not connectomics? Nat. Methods 10, 494–500 (2013).

63. Fitts, P. M. & Posner, M. I. Human performance. (1967).

64. Tapia, J. C. et al. Pervasive synaptic branch removal in the mammalian neuromuscular system at birth. Neuron 74, 816–829 (2012).

65. Lichtman, J. W. & Colman, H. Synapse elimination and indelible memory. Neuron 25, 269–278 (2000).

66. Walsh, M. K. & Lichtman, J. W. In vivo time-lapse imaging of synaptic takeover associated with naturally occurring synapse elimination. Neuron vol. 37 67–73 (2003).

67. Wilson, A. M. et al. Developmental Rewiring between Cerebellar Climbing Fibers and Purkinje Cells Begins with Positive Feedback Synapse Addition. Cell Rep. 29, 2849– 2861.e6 (2019).

68. Sheehan, D. C. & Hrapchak, B. B. Theory and practice of histochemistry. CV Mosby, St Louis (1980).

69. OpenCV: cv::SimpleBlobDetector Class Reference. https://docs.opencv.org/3.4/d0/d7a/classcv_1_1SimpleBlobDetector.html.

70. Lowe, D. G. Distinctive image features from scale-invariant keypoints. Int. J. Comput. Vis. 60, 91–110 (2004).

71. Fischler, M. A. & Bolles, R. C. Random Sample Consensus: A Paradigm for Model Fitting with Applications to Image Analysis and Automated Cartography. Readings in Computer Vision 726–740 (1987) doi:10.1016/b978-0-08-051581-6.50070-2.

72. Toldo, R. & Fusiello, A. Robust Multiple Structures Estimation with J-Linkage. Lecture Notes in Computer Science 537–547 (2008) doi:10.1007/978-3-540-88682-2_41.

73. Rublee, E., Rabaud, V., Konolige, K. & Bradski, G. ORB: An efficient alternative to SIFT or SURF. 2011 International Conference on Computer Vision (2011) doi:10.1109/iccv.2011.6126544.

74. Nealen, A., Müller, M., Keiser, R., Boxerman, E. & Carlson, M. Physically Based Deformable Models in Computer Graphics. Computer Graphics Forum vol. 25 809–836 (2006).

75. Swope, W. C., Andersen, H. C., Berens, P. H. & Wilson, K. R. A computer simulation method for the calculation of equilibrium constants for the formation of physical clusters of molecules: Application to small water clusters. The Journal of Chemical Physics vol. 76 637–649 (1982).

76. Berger, D. R., Seung, H. S. & Lichtman, J. W. VAST (Volume Annotation and Segmentation Tool): Efficient Manual and Semi-Automatic Labeling of Large 3D Image Stacks. Front. Neural Circuits 12, 88 (2018).

77. Sato, M., Bitter, I., Bender, M. A., Kaufman, A. E. & Nakajima, M. TEASAR: tree- structure extraction algorithm for accurate and robust skeletons. Proceedings the Eighth Pacific Conference on Computer Graphics and Applications doi:10.1109/pccga.2000.883951.

78. seung-lab. seung-lab/kimimaro. https://github.com/seung-lab/kimimaro.

79. He, K., Zhang, X., Ren, S. & Sun, J. Deep residual learning for image recognition. arXiv preprint arXiv:1512. 03385 (2015).

80. Chen, T., Kornblith, S., Norouzi, M. & Hinton, G. A Simple Framework for Contrastive Learning of Visual Representations. (2020).

81. McInnes, L., Healy, J., Saul, N. & Großberger, L. UMAP: Uniform Manifold Approximation and Projection. Journal of Open Source Software vol. 3 861 (2018).

82. Çiçek, Ö., Abdulkadir, A., Lienkamp, S. S., Brox, T. & Ronneberger, O. 3D U-Net: Learning Dense Volumetric Segmentation from Sparse Annotation. arXiv [cs.CV] (2016).

83. Yuste, R. Dendritic Spines. (MIT Press, 2010).

84. Campello, R. J. G. B., Moulavi, D. & Sander, J. Density-Based Clustering Based on Hierarchical Density Estimates. in Advances in Knowledge Discovery and Data Mining 160–172 (Springer Berlin Heidelberg, 2013).

85. Goodman, L. A. On Simultaneous Confidence Intervals for Multinomial Proportions. Technometrics 7, 247–254 (1965).

